# Evidence of functional connectivity disruptions between auditory and non-auditory regions in adolescents living with HIV

**DOI:** 10.1101/2025.02.16.638287

**Authors:** Joanah Madzime, Marcin Jankiewicz, Ernesta M. Meintjes, Peter Torre, Barbara Laughton, Martha J. Holmes

## Abstract

Children living with perinatally acquired HIV (CPHIV) demonstrate hearing impairments and language processing delays even in the presence of combination antiretroviral therapy (cART). Even though efficient processing of sound relies on the proper function of both the peripheral and central auditory auditory systems (PAS; CAS), investigations on the effect of HIV on the auditory system have predominantly focused on the PAS. Additionally, language processing requires the efficient interaction between CAS brain regions and non-auditory regions. Investigating the functional connectivity (FC) within the CAS and between the CAS and non-auditory regions may reveal the influence of HIV on regions involved in auditory function.

Within a Bayesian stastical framework, we used resting-state functional magnetic resonance imaging (RS-fMRI) to map FC in the CAS as well as between CAS regions and non-auditory regions of 11-year-old CPHIV. Graph theory was used to investigate the regional effects of HIV on functional brain network properties. Finally we explored the relationships between FC and neurocognitive outcomes relating to auditory and language processing. We hypothesized that CPHIV would show disruptions in FC between CAS regions as well as between CAS and non-auditroy regions. Secondly, we hypothesized that in CPHIV, regional brain network properties would be altered compared to their uninfected peers (CHUU). Finally we hypothesized that FC and functional network regional outcomes would be related to neurocognitive outcomes.

Our investigation revealed lower FC of the primary auditory cortex (PAC) in CPHIV as well as disruptions in FC between CAS regions (including the PAC) and non-auditry regions including hippocampal sub-regions, the lingual gyri and basal ganglia. Functional network analysis reavealed lower nodal degree and efficiency in CAS regions including the cochlear nucleus/superior olivary complex (CN/SOC) and the inferior colliculus (IC). We also report associations between the nodal efficiency of middle temporal and superior frontal regions and delayed recall, a neurocognitive marker of working memory, present in CHUU but not in CPHIV.

Our results demonstrate FC alterations in the PAC and between CAS regions and non-auditory regions involved in limbic, visual and motor processing, as well as disruptions to the regional properties of the CAS regions as part of the functional brain network. These results provide insight into the state of the CAS FC in the presence of HIV and its possible role in the hearing and language impairments seen in this population.

## Introduction

The efficient processing of sound involves both the perception of sound by the peripheral nervous system and the processing of auditory signals in the central nervous system (CNS) (Litovsky, 2015). While basic auditory abilities are developed at birth, continuous exposure to auditory stimulation in childhood results in functional changes in the central auditory system (CAS) (Fitzroy et al., 2015; Skoe et al., 2015). The CAS, which comprises the cochlear nucleus (CN) and inferior colliculus (IC) in the brainstem, medial geniculate nucleus (MGN) in the thalamus and primary auditory cortex (PAC) in the superior temporal gyrus (STG), interacts heavily with other brain areas, including the temporal and frontal lobe regions, for efficient language processing and communication (Maffei et al., 2015; Middlebrooks et al., 2016).

Children perinatally infected with the human immunodeficiency virus (CPHIV) are at risk of hearing impairments and disruptions in language development (Chao et al., 2012; Maro et al., 2017; Rice et al., 2013; Torre et al., 2012, 2015). Associations between auditory function and HIV status have been reported (Niemczak et al., 2021). Further, CPHIV are at risk of lower working memory and verbal fluency outcomes (Lima et al., 2023; Morgan et al., 2015). Investigating the function of the CAS may provide a window into the relationship between HIV, language and the brain in these at-risk populations.

Resting-state functional magnetic resonance imaging (RS-fMRI) is a tool to investigate co-activation of anatomically separate brain areas, which reflects functional communication between areas (van den Heuvel & Hulshoff Pol, 2010). In the literature, functional co-activation is often referred to as functional connectivity (FC). Coupled with a graph theory analysis, RS-fMRI can provide insight into information transfer and integration in the functional brain network. In a RS-fMRI based graph-theoretical framework, a graph is a set of brain regions (nodes) functionally connected by edges. Using this approach, we can investigate network measures of integration and segregation at the regional/nodal level as well as at a global level revealing the arrangement or topology of the network and its regions (Bullmore et al., 2016).

To date, there are only a handful of RS-fMRI studies in the context of children living with HIV and none focused on the CAS. Yadav et al (2018) reported lower FC in the middle temporal gyrus MTG of CPHIV, which could potentially affect auditory processing (Yadav et al., 2018). Another study showed HIV-related altered FC between brain regions in the DMN and regions belonging to RSNs that work with the auditory network, including the visual and somatosensory networks (Herting et al., 2015). Our group has used neuroimaging techniques to study brain development since age 5 years in a cohort of CPHIV who initiated antiretroviral therapy (ART) in infancy. At 7 years old, we observed reduced FC in the inferior frontal gyrus of CPHIV, a region that is associated with the auditory RSN (Toich et al., 2018). Although these findings allude to potential alterations in auditory functional regions in CPHIV, investigating the specific regions involved in auditory function as well as their interactions with other brain regions that participate in language processing and communication is necessary.

We used RS-fMRI and a graph-theoretical framework to assess FC of the CAS in a cohort of 11-year old children. We hypothesized significant differences in FC in CPHIV as compared to CHUU children. Furthermore, we posited that the CAS in CPHIV would have reduced functional strength and efficiency compared to CHUU. Lastly, to better understand the possible relationship to auditory outcomes, we explored associations between functional deficits and cognitive outcomes related to language and hearing.

## Methods

### Study cohort

Participants were 84 (11–12-year-old) children (25 CHUU, 59 CPHIV) who completed structural and RS-fMRI scans. The children living with HIV were recruited from the randomized Children with HIV Early Antiretroviral Therapy (CHER) trial and have been monitored since birth at the Family Centre for Research with Ubuntu (FAMCRU), Tygerberg Children’s Hospital, in Cape Town, South Africa (Cotton et al., 2014; Violari et al., 2008). For the CHER trial, infants were randomized into one of three treatment groups; two arms involved early (before 12 weeks of age) ART and one deferred ART until CD4%<25% in the first year or CD4%<20% thereafter, or if clinical disease progression criteria presented (Cotton et al., 2014). The uninfected children were recruited from a parallel vaccine study (Madhi et al., 2010).

### T1-weighted (T1w) structural and RS-fMRI Image acquisition

T1w structural and RS-fMRI acquisitions were done in the same session using a 3 Tesla Siemens Skyra MRI scanner (Erlangen, Germany) at University of Cape Town’s Cape Universities Body Imaging Centre (CUBIC). Children were first familiarized with the scanning procedures on a mock scanner. The T1w structural was acquired for 6 mins using a multiecho magnetization prepared rapid gradient echo (ME-MPRAGE) sequence with TR = 2530 ms, TEs = 1.69/3.54/5.39/7.24ms, inversion time (TI) 1,100ms, flip angle = 7°, and FOV = 224×224×176mm^3^. RS-fMRI data were acquired for 6 mins using an interleaved multi-slice 2D gradient echo, echo planar imaging (EPI) sequence with the following parameters: 33 interleaved slices, slice thickness 4mm, slice gap 1mm, voxel size 3×3×4mm^3^, FOV = 250×250×164mm^3^, TR/TE = 2,000/30ms, flip angle 90°, 180 EPI volumes.

### Neurodevelopmental measures: KABC-II Auditory/language-related scores

Participants were administered the Kaufman Assessment Battery for Children – Second Edition (KABC-II) (Kaufman & Kaufman, 2014) by trained research assistants supervised by a licensed psychologist. The neurodevelopmental scores were part of the Kaufman Assessment Battery for Children – Second Edition (KABC-II) (Kaufman & Kaufman, 2014) and were measured in the same group of children at 11 years old. Subtests of the battery include Learning, (Atlantis, Atlantis delayed, Rebus, Rebus delayed, Delayed Recall) which assesses the ability to store and retrieve information, Planning (Story completion and Pattern reasoning) which assesses the ability to solve nonverbal problems, Sequential processing (Number recall, Word order and Hand movement) which assesses ability to solve problems using auditory working memory and visual information presented serially, and Simultaneous processing (Rover, Block Counting, Conceptual Thinking, Triangles, Face Recognition, and Gestalt Closure) which assesses the ability to solve spatial or logistical problems that require simultaneous processing of multiple related stimuli (Kaufman & Kaufman, 2014). The sum of all scores generates a global cognitive score called the Mental Processing index. Moreover, summing the subtests that do not require verbal expression (Hand Movements, Block Counting, Triangles, Pattern Reasoning, Story Completion, Conceptual Thinking, and Face Recognition) generates the Nonverbal Index score (Kaufman & Kaufman, 2014). For the current study we chose the tests related to auditory and language processing: Number recall, Word order, Learning, Atlantis, Atlantis delayed, Rebus, Rebus delayed and Delayed Recall, Story completion, Sequential processing, Mental processing and Nonverbal index. Moreover, we included Animal naming, a neurocognitive score relating to word generation and categorization (Strauss et al., 2010).

### Ethical Considerations

The study protocol was approved by the Human Research Ethics Committees of the University of Cape Town and Stellenbosch University. Parents/guardians provided written informed consent and children oral assent.

### RS-fMRI pre-processing

The process described here was done for each subject in the Analysis of Functional Neuroimages (AFNI) software. The first 5 EPI volumes were removed to account for the time it takes for signal stabilization. Then we ‘despiked’ each volume to truncate the signal in voxels with artificially high signal, and performed temporal interpolation to adjust the acquisition time of each slice to that of the first slice of the volume. Next each EPI volume was co-registered to the reference EPI volume (6^th^ EPI volume in the current study), generating 6 motion parameters (3 rotations and 3 translations) per volume. The motion estimates were de-meaned by removing the mean per run, and per-TR motion derivates were computed for each volume as the backward difference between the motion estimates of said volume and the prior volume (i.e. Δx(t)=x(t)-x(t-1), where x is any one of the 6 motion parameters). The EPI volumes were spatially smoothed with a Gaussian kernel of 6mm full width at half maximum (FWHM), and a brain mask of the EPI dataset created. The T1w structural volume was next segmented into white matter (WM), gray matter (GM) and cerebrospinal fluid (CSF). We then aligned the subject’s EPI volumes to his/her T1w structural volume, and used the resulting co-registered CSF and WM masks to extract WM and CSF signals from the EPI data. Finally, signals from non-neuronal sources were removed from the EPI data by regressing out the mean WM and CSF signals, as well as the 6 de-meaned motion parameters and their per-TR derivatives. We censored pairs of TRs where the Euclidean norm (i.e. the square root of the sum of squares) of the per-TR motion derivatives exceeded 0.3 mm, as well as TRs for which 5% or more of the brain mask was classified as outliers by afni_proc. Notably, a per-TR Euclidean norm of 0.3 mm corresponds to 0.3 mm motion in a single direction, or translations of 0.12 mm and rotations of 0.11° in each direction. Participants with fewer than 160 EPI volumes after truncation were excluded from further analyses. EPI time series data of remaining participants were truncated to 160 volumes.

### GM regions of interest

To fully examine the CAS, we included both manually traced and automatically segmented regions of interest (ROI). ROIs included 4 bilateral manually traced GM regions that form part of the auditory pathway and 126 automatically segmented cortical and subcortical GM regions. The manual tracings were performed on the T1w structural volumes by an expert neuroanatomist using Freeview software (FreeSurfer v7.1.0 image analysis suite: (http://surfer.nmr.mgh.harvard.edu/) on a Lenovo ThinkPad Yoga370 tablet. The 4 manually traced auditory ROIs included a region at the junction between brainstem pons and medulla and consists of the cochlear nucleus and superior olivary complex (CN/SOC), the IC in the midbrain, the MGN in the thalamus and the PAC. For automatic parcellation we used Connectome mapper (CMP) 3 (v3.0.0-RC4), an imaging processing pipeline to produce multi-resolution morphological subject atlases (Daducci et al., 2012; Tourbier et al., 2022). We used the lowest resolution which consisted of a set of 126 subcortical and cortical ROIs. A list of all the ROIs in the atlas and their abbreviations can be found in *Appendix A: Automatically segmented regions of interest.* We then merged the manually traced and automatically segmented ROIs. Automatically segmented ROIs that overlapped with manually traced ROIs were excluded. The number of excluded ROIs varied between 81 and 122 across subjects. Finally, 80 ROIs were common to all participants. Masks of these ROIs were mapped to the RS-fMRI mask and used as seeds for the FC analyses.

### FC and functional network definition

The process described here was done for each subject. The time series signals within each seed were extracted to compute the mean time series for each ROI. We calculated a Pearson correlation coefficient between the mean time series of each ROI and every other ROI (*3dNetCorr)* (Taylor & Saad, 2013). The correlation coefficients were Fisher Z-transformed to obtain approximately normally distributed correlations. This resulted in an 80×80 correlation matrix for each subject with 3160 connections. Of those connections, 604 pairs included at least one auditory ROI. This matrix was the input for our FC analysis.

To define each subject’s functional network, we discarded weak and/or negative edge weights (Fisher Z-transformed r values < 0.1003 equivalent to Pearson r values < 0.1000) from each subject’s 80×80 Fisher Z-transformed matrix (Figure 1). Removing negative edge weights allows for an intuitive interpretation of the relationship between nodes i.e. strong positive weights mean nodes work together in a correlated way (Fornito et al., 2016; Rubinov & Sporns, 2010). We then computed nodal graph measures using iGraph and braingraph packages in R (v 4.1.3). We computed nodal degree and nodal strength – measures of the number and weightings of connections to a node, respectively. Nodes with high degree or strength tend to be identified as network hubs as they are generally positioned where they can make strong contributions to the global network (Hoff et al., 2013; van den Heuvel & Sporns, 2013). In addition we computed nodal transitivity – a measure of densely connected groups of nodes. Nodes with high transitivity in the brain are surrounded by highly interconnected nodes forming clusters that may translate as functional coordination (Fornito et al., 2016). Finally we computed nodal efficiency and nodal local efficiency. Nodal efficiency is a measure of the efficiency of communication between a node and all the other nodes in the network and nodes that have high degree tend to have high efficiency (Fornito et al., 2016; Sporns, 2018). On the other hand, nodal local efficiency is a measure of the efficiency of the graph formed in the absence of the node. This measure shows the graph’s level of fault tolerance and nodes with high nodal local efficiency reflect efficient integration with their neighbors (Fornito et al., 2016).

**Figure 1:**
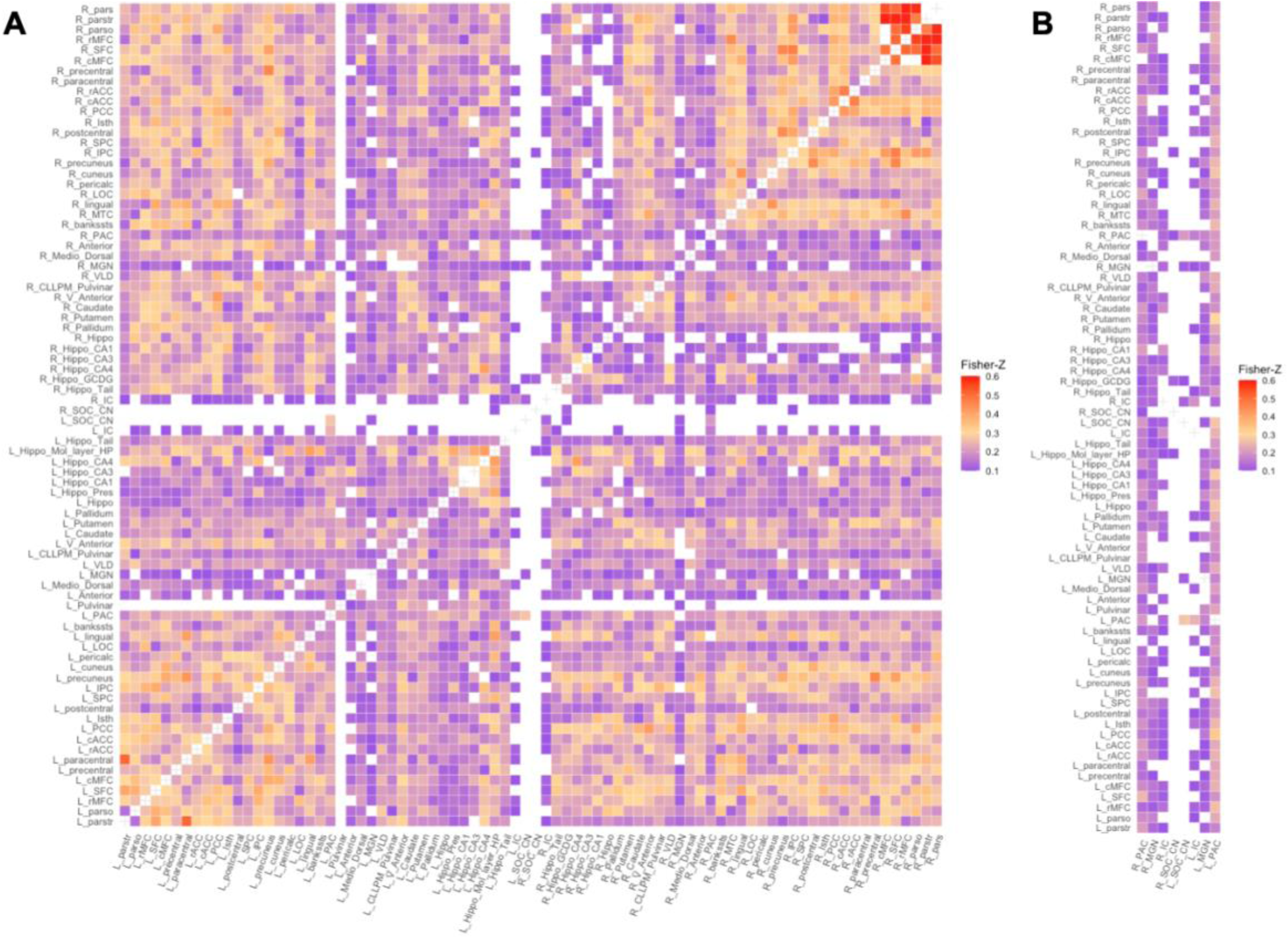
Functional network of a random control subject: Fisher Z-transformed r values of weak and negative connections (Fisher Z-transformed r value < 0.1003) were discarded. A – Full 80×80 RS connectivity matrix, B – the columns containing auditory ROIs. From left column to right: (R) PAC, (R) MGN, (R) IC, (R) SOC/CN, (L)_ SOC/CN, (L) IC, (L) MGN, (L) PAC. (R/L) – Right/Left.

### Statistical analyses

To investigate the effects of living with HIV on both FC and functional network organization we employed Bayesian multilevel modelling (BML) (Chen, Bürkner, et al., 2019; Chen, Xiao, et al., 2019). An advantage of the Bayesian framework is its non-dichotomous interpretation of results, unlike binary significance of the p-value statistic in conventional null-hypothesis significance testing (NHST) (Kruschke & Liddell, 2018). Briefly, consider a scenario where we are investigating the difference between two group means (effect), CHUU-CPHIV. Under the null hypothesis (H0), the effect equals 0. A Bayesian analysis estimates a probability distribution (posterior distribution) of the alternative hypothesis (H1: effect > 0), based on the data, i.e., P(effect>0|data). The data is assigned a prior probability distribution based on an assumption, such as that the ROIs are connected to each other through Gaussian or Student’s t-distribution. The area under the posterior distribution curve that is greater than 0, the posterior probability (P+), is interpreted as the strength of evidence that the effect is positive, given the data. That is, the strength of evidence that the reference group (CHUU) has a higher mean than the test group (CPHIV), based on the data. A very high P+ represents a strong probability that the effect is greater than 0, while a very low P+ means there is a weak probability the effect is greater than 0. Note, a very low P+ corresponds with a strong probability of the effect being less than 0. Figure 2 shows the posterior probability distribution for the nodal strength of a random brain ROI, for the comparison between CHUU and CPHIV.

**Figure 2:**
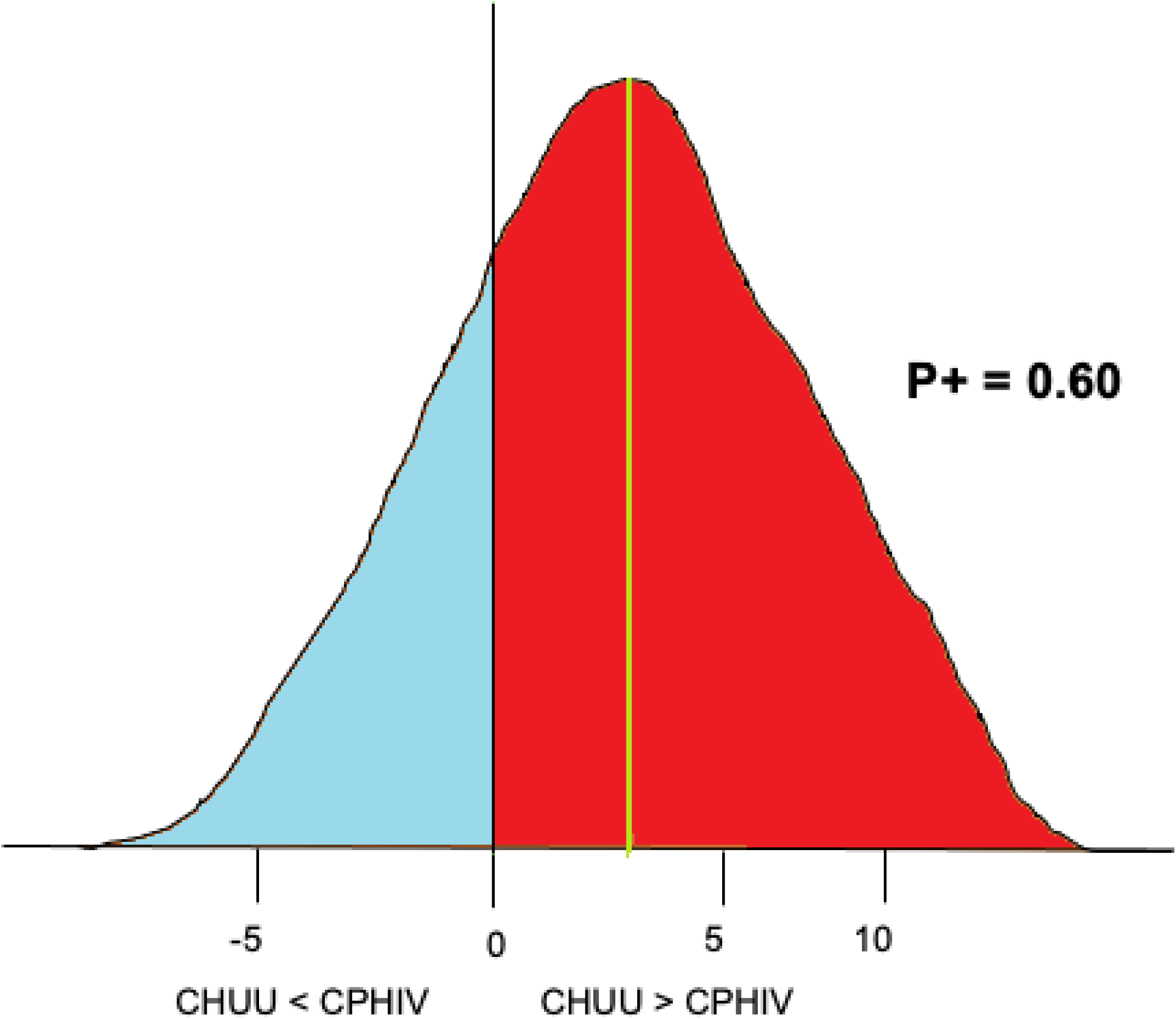
An example of posterior probability distribution of the nodal strength of a randomly chosen brain ROI for the comparison CHUU-CPHIV. The horizontal axes indicates the effect size, the vertical axes (black) indicates zero effect or difference between the two group means and the vertical green line indicates the mean difference. Here, the area under the curve greater than 0 (red), P+ = 0.60, indicates we found with 60% probability that the CHUU group has a bigger mean than CPHIV given the data.

Furthermore, using the framework, a multilevel model is formed which integrates variability estimates at the group and subject levels in estimating the uncertainty of effect at each brain region or ROI. This integrative model is done via Monte-Carlo Markov chains (MCMC) at a predetermined number of iterations resulting in the partial pooling of the effect of interest towards the centre (Chen, Bürkner, et al., 2019; Chen, Xiao, et al., 2019). BML for investigating effects is particularly well suited for neuroimaging studies as we know brain regions are not independent i.e. one ROI could be involved in multiple pairwise connections. In addition, unlike the NHST assumption that the likelihood of an effect is equally anywhere between negative infinity and positive infinity for every brain region, BML assumes the likelihood of an effect more likely follows a Gaussian distribution (Chen et al., 2021). In NHST models this is corrected for via multiple comparisons correction but the BML approach not only negates the need for multiple comparisons correction, it also captures the covariant structure of the brain, allowing for a more robust and efficient inference.

To investigate potential FC group differences, we ran the matrix-based analysis (MBA) program which uses BML to pool and share variability estimates of ROIs in each subject’s Fisher Z-transformed correlation matrix. The contribution of each ROI in the matrix is captured in the model, showing its overall contribution to connections in the matrix. MBA offers effect estimations of both region and region pairs or connections (Chen, Bürkner, et al., 2019). The reporting of ROI effects is a unique feature of MBA, outputting information about regional contributions. Region effects characterize regional contributions within a multilevel framework, complementing paired FC. We included sex as a confounding variable in all our models because studies find biological sex is related to different patterns of functional activation, which may be confounded by pubertal hormones during adolescence (Gur & Gur, 2016; Lenroot & Giedd, 2010). Our final MBA model was of the form:

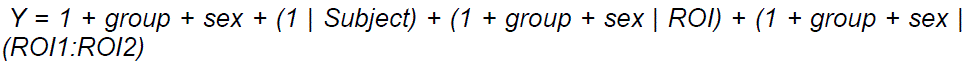

where ROI1:ROI2 represent a connection. As such the resulting effect estimations were at the ROI level i.e. 80 ROIs and region pair level i.e. 3160 region pairs.

To investigate functional network organization between CHUU and CPHIV, we ran the region-based analysis (RBA) program for each graph theory measure (Chen, Xiao, et al., 2019). RBA uses BML to pool and share information among brain regions or nodes. Here we also included sex as a confounding variable and our final RBA model was of the form:

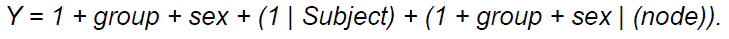

The resulting effect estimates were for every node.

To assess the quality of MCMC we used the split Ř statistic which gauges the consistency of an ensemble of Markov chains. Fully converged chains correspond to Ř = 1. In the present study we used an arbitrary threshold of Ř < 1.1 for chains that converged well in both the MBA and RBA analyses.

#### Associations between functional outcomes and auditory/language-related scores

We examined the association between functional outcomes (FC between ROIs/graph measure outcomes for ROIs) and KABC-II auditory/language-related neurodevelopmental scores. Associations were investigated among connections or ROI pairs that included an auditory ROI i.e. 604 connections. Graph measure associations were investigated with all 5 graph measures (i.e. node degree, strength, transitivity, nodal and local efficiency) for each of the 80 nodes.

To investigate the associations we ran Pearson correlation statistical tests. Where a variable was not normally distributed (Shapiro-Wilk p <0.05), we investigated the relationship via Spearman correlation statistical test as it is resistant to outliers. We included sex as a confounding variable and investigated handedness as potential confounder due to evidence of functional asymmetry in language processing areas that develops with age (Olulade et al., 2020; Vassal et al., 2016). We corrected for multiple comparisons using the False Discovery Rate (FDR) method and considered FDR q < 0.05 as our significance level.

#### Associations between functional outcomes and immune health markers

We ran analyses to investigate the association between functional outcomes and immune health markers within the CPHIV group. Correlation tests were either Pearson or Spearman depending on normality. We investigated early immune health (CD4% at study enrolment), immune health of participant at the time of 11 year scan and treatment related health markers (age at cART initiation and first VL suppression). We corrected for multiple comparisons using the FDR method and considered FDR q < 0.05 as our significance level.

## Results

Out of 84 participants who underwent MRI scanning, a total of 6 participants (2 CHUU; 4 CPHIV) were excluded due to poor image quality and/or not meeting motion inclusion criteria. We present results from a sample of 78 children (23 CHUU; 55 CPHIV), demographics of which are presented in Table 1.

**Table 1:**
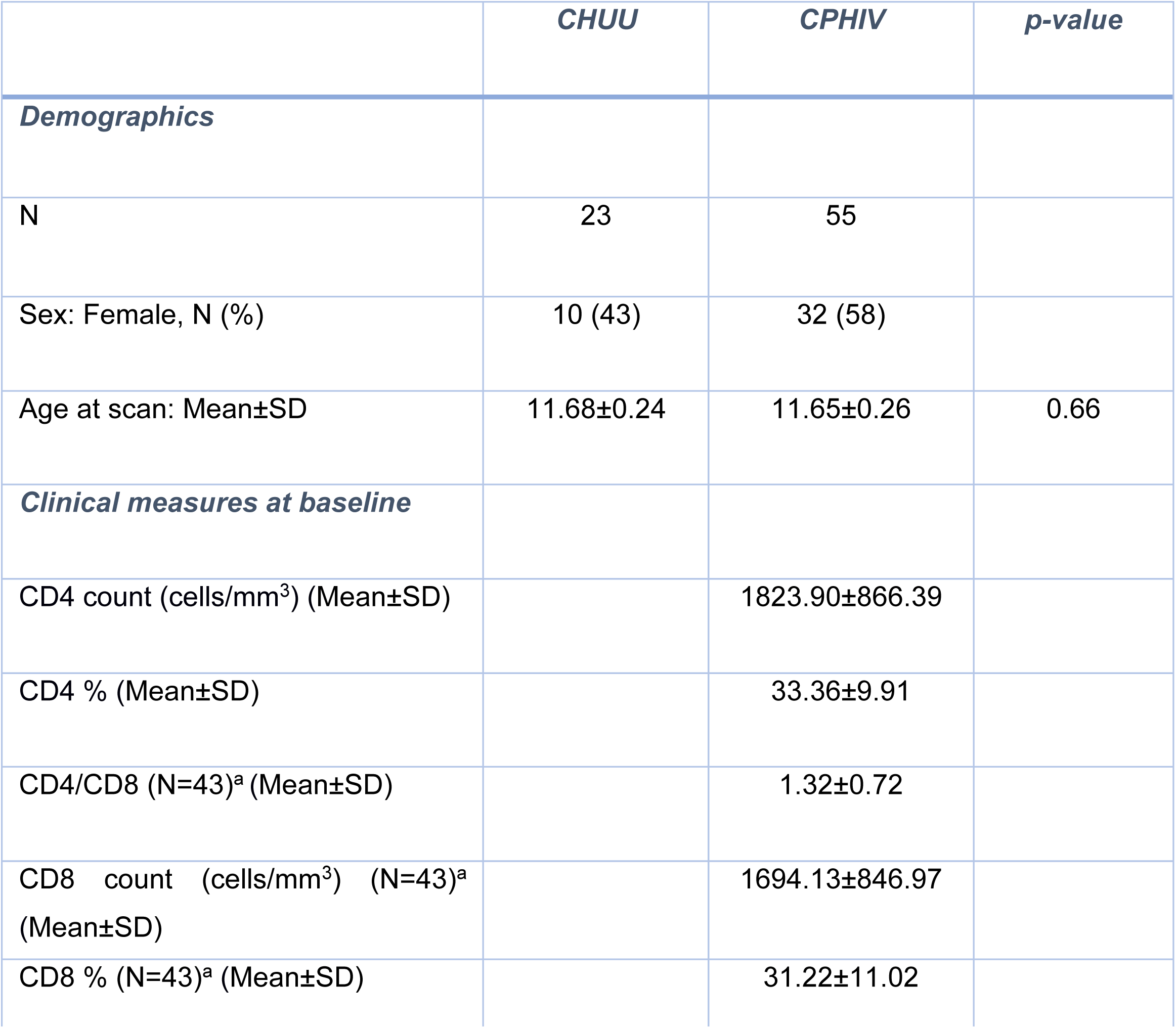
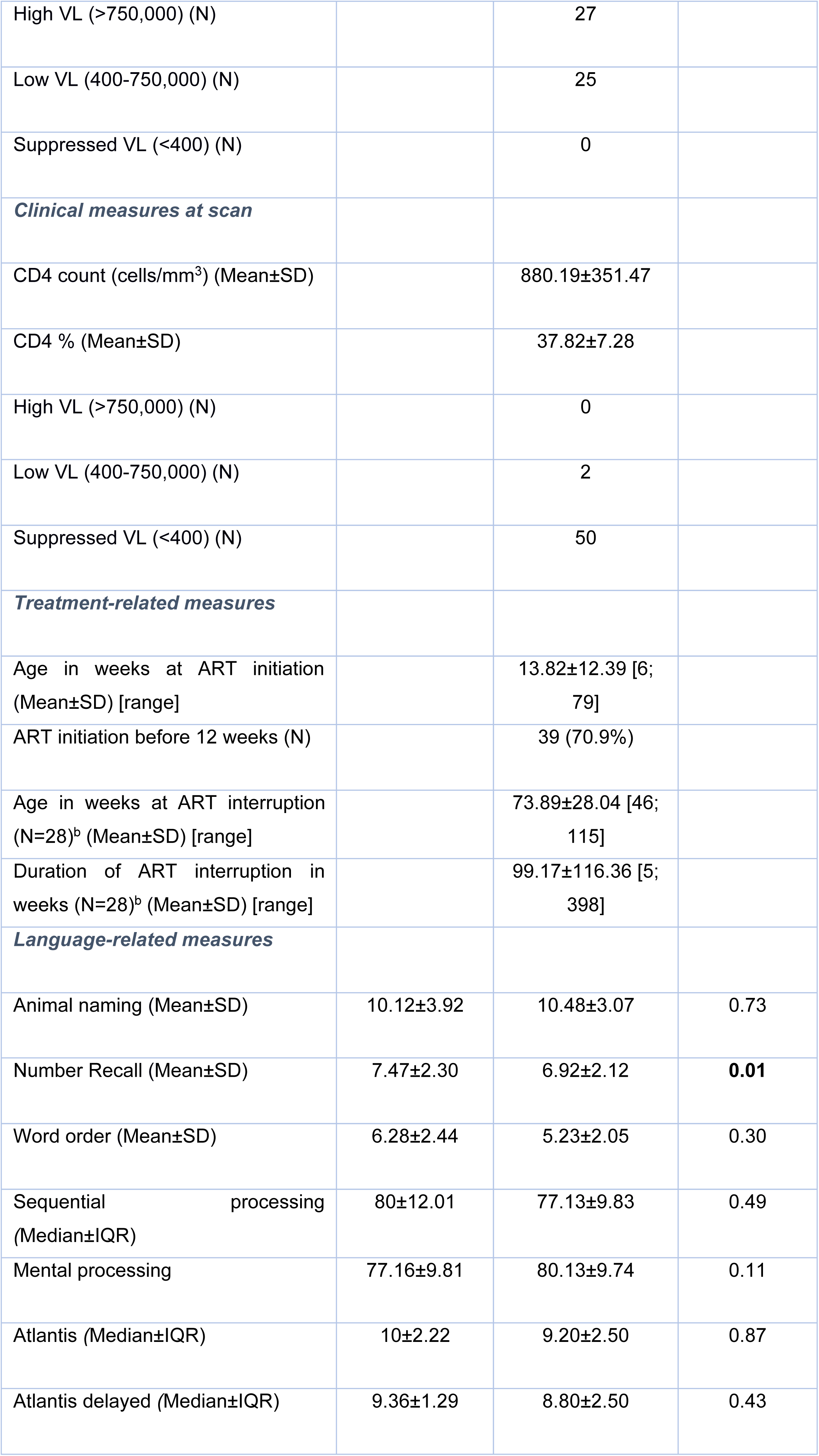
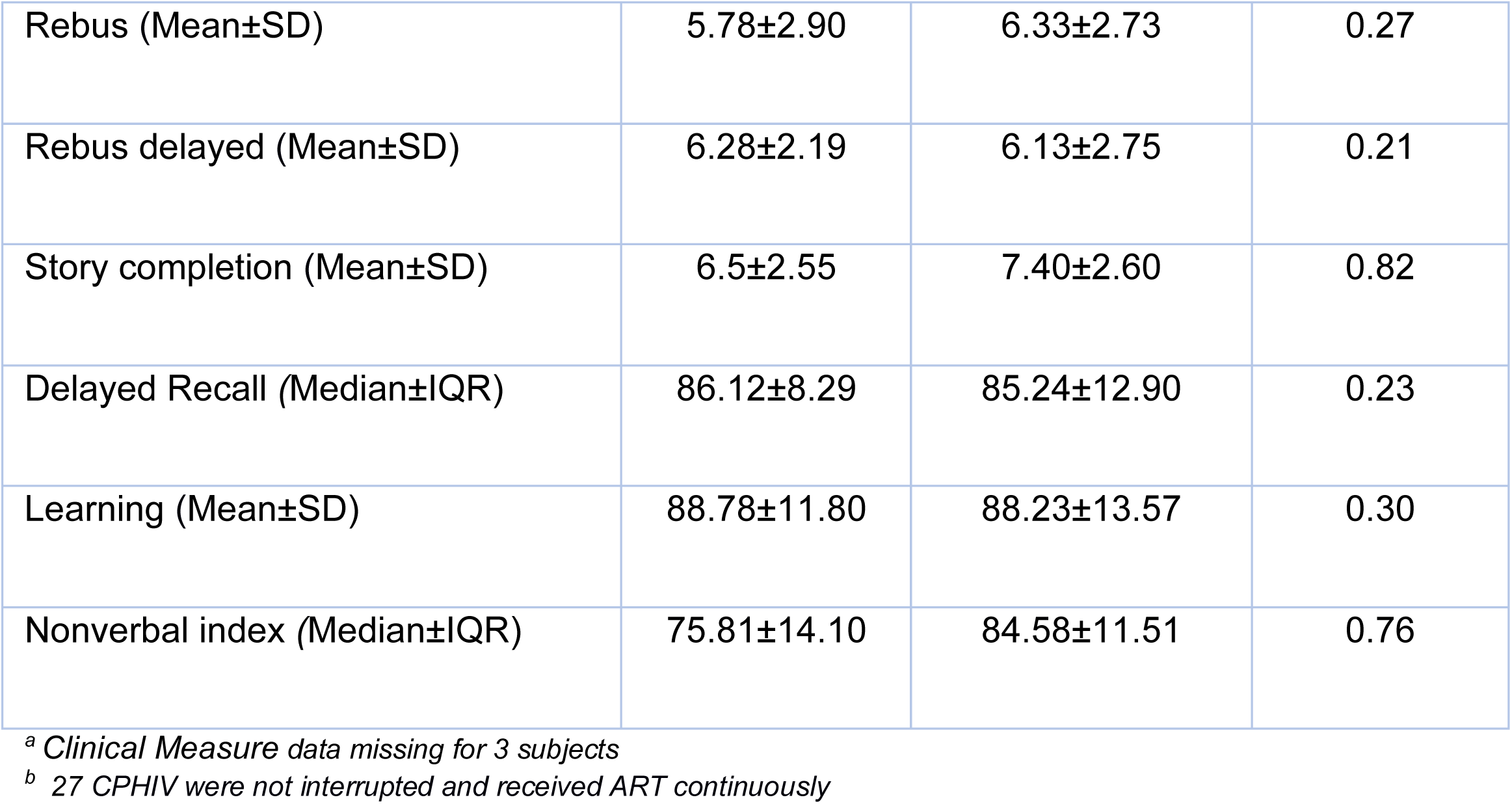
Sample characteristics of children perinatally infected and living with (CPHIV) and uninfected controls (CHUU) (N = 78). Where a variable was not normally distributed (Shapiro-Wilk p <0.05) we summarize with median and IQR.

|  | **CHUU** | **CPHIV** | **p-value** |
| --- | --- | --- | --- |
| **Demographics** |  |  |  |
| N | 23 | 55 |  |
| Sex: Female, N (%) | 10 (43) | 32 (58) |  |
| Age at scan: Mean±SD | 11.68±0.24 | 11.65±0.26 | 0.66 |
| **Clinical measures at baseline** |  |  |  |
| CD4 count (cells/mm<sup>3</sup>) (Mean±SD) | | 1823.90±866.39 | |
| CD4 % (Mean±SD) | | 33.36±9.91 | |
| CD4/CD8 (N=43)<sup>a</sup> (Mean±SD) | | 1.32±0.72 | |
| CD8 count (cells/mm<sup>3</sup>) (N=43)<sup>a</sup> (Mean±SD) | | 1694.13±846.97 | |
| CD8 % (N=43)<sup>a</sup> (Mean±SD) | | 31.22±11.02 | |
| High VL (>750,000) (N) |  | 27 |  |
| Low VL (400-750,000) (N) |  | 25 |  |
| Suppressed VL (<400) (N) |  | 0 |  |
| *Clinical measures at scan* |  |  |  |
| CD4 count (cells/mm<sup>3</sup>) (Mean±SD) |  | 880.19±351.47 |  |
| CD4 % (Mean±SD) |  | 37.82±7.28 |  |
| High VL (>750,000) (N) |  | 0 |  |
| Low VL (400-750,000) (N) |  | 2 |  |
| Suppressed VL (<400) (N) |  | 50 |  |
| *Treatment-related measures* |  |  |  |
| Age in weeks at ART initiation (Mean±SD) [range] |  | 13.82±12.39 [6; 79] |  |
| ART initiation before 12 weeks (N) |  | 39 (70.9%) |  |
| Age in weeks at ART interruption (N=28)<sup>b</sup> (Mean±SD) [range] |  | 73.89±28.04 [46; 115] |  |
| Duration of ART interruption in weeks (N=28)<sup>b</sup> (Mean±SD) [range] |  | 99.17±116.36 [5; 398] |  |
| *Language-related measures* |  |  |  |
| Animal naming (Mean±SD) | 10.12±3.92 | 10.48±3.07 | 0.73 |
| Number Recall (Mean±SD) | 7.47±2.30 | 6.92±2.12 | **0.01** |
| Word order (Mean±SD) | 6.28±2.44 | 5.23±2.05 | 0.30 |
| Sequential processing (Median±IQR) | 80±12.01 | 77.13±9.83 | 0.49 |
| Mental processing | 77.16±9.81 | 80.13±9.74 | 0.11 |
| Atlantis (Median±IQR) | 10±2.22 | 9.20±2.50 | 0.87 |
| Atlantis delayed (Median±IQR) | 9.36±1.29 | 8.80±2.50 | 0.43 |
| Rebus (Mean±SD) | 5.78±2.90 | 6.33±2.73 | 0.27 |
| Rebus delayed (Mean±SD) | 6.28±2.19 | 6.13±2.75 | 0.21 |
| Story completion (Mean±SD) | 6.5±2.55 | 7.40±2.60 | 0.82 |
| Delayed Recall (Median±IQR) | 86.12±8.29 | 85.24±12.90 | 0.23 |
| Learning (Mean±SD) | 88.78±11.80 | 88.23±13.57 | 0.30 |
| Nonverbal index (Median±IQR) | 75.81±14.10 | 84.58±11.51 | 0.76 |

<sup>a</sup> Clinical Measure data missing for 3 subjects
<sup>b</sup> 27 CPHIV were not interrupted and received ART continuously

We grouped our results based on strength of evidence as follows: very strong evidence of an effect – P+ >= 0.975 or P+ <= 0.025, strong evidence of an effect – P+ >= 0.95 or P+ <= 0.05 and evidence of an effect – P+ >= 0.90 or P+ <= 0.10. Our discussion mostly focuses on the connections that showed strong evidence of an effect.

### Functional connectivity results

The Ř values were less than 1.1 for the CHUU vs CPHIV FC analysis indicating all MCMC chains converged well. For a summary of model results please refer to *Appendix B: Summary information of model results for FC of CHUU vs CPHIV*.

#### ROI effects

Among the 80 ROIs, 14 ROIs showed strong evidence of lower FC in the CPHIV group. Regions include the left PAC, bilateral hippocampus, bilateral hippocampus CA1, bilateral hippocampus CA3, bilateral hippocampus CA4, left hippocampus molecular layer, left hippocampus presubiculum, right hippocampus GCDG, and bilateral lingual gyrus. In addition, 10 ROIs showed strong evidence of higher FC (P+<=0.05) in the CPHIV group including the left mediodorsal thalamic nucleus, right anterior thalamic nucleus, left caudate, left postcentral gyrus, right precentral gyrus, right rMFC, right parsopecularis, right pallidum and bilateral putamen (Figure 3). Table 3 in Appendix C shows the ROI effect estimates including uncertainty intervals and posterior probabilities of the effect being positive.

**Figure 3:**
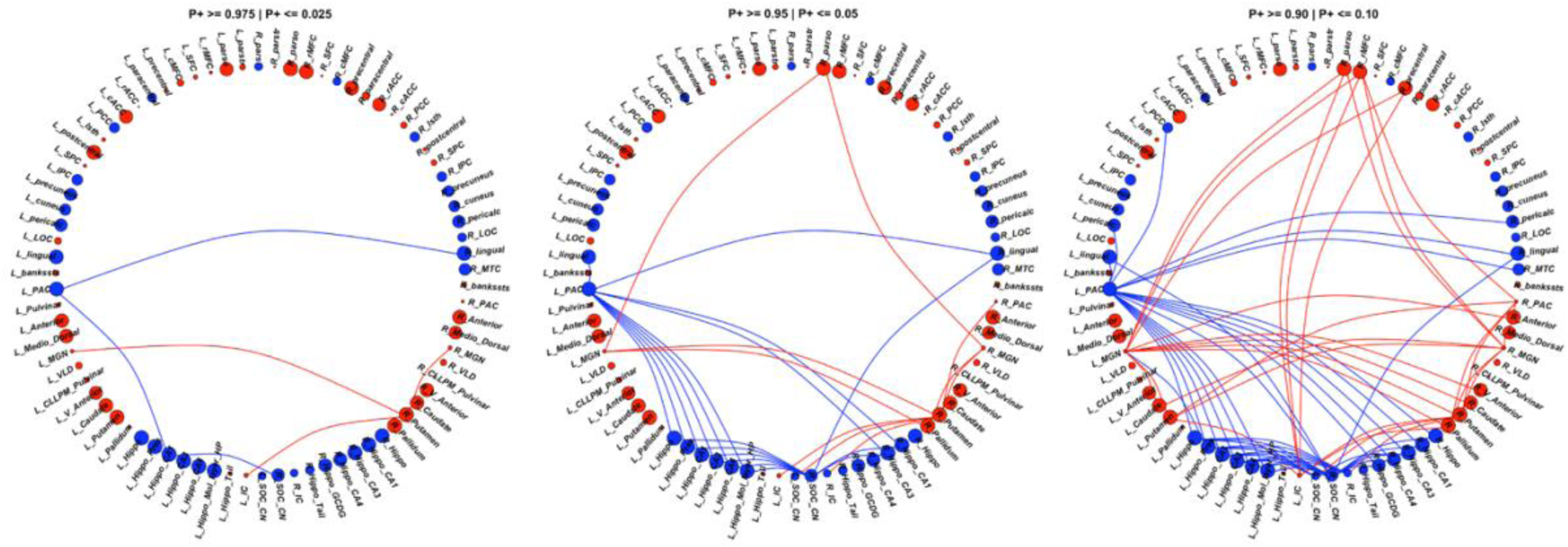
We found 73 connections showing evidence (P+>=0.90 | P+<=0.10) of altered FC between the CPHIV and CHUU groups (right). 43 connections **showed lower FC (blue links)** while 30 connections showed **higher FC (red links)** in the CPHIV group compared to CHUU. We found 29 connections showing **strong evidence (P+>=0.95 | P+<=0.05)** (middle) and 6 connections showing **very strong evidence (P+>=0.975 | P+<=0.025)** of altered FC (left) between groups. ROIs are colored according to direction of ROI effect with red and blue ROIs indicating higher and lower ROI FC in the CPHIV compared to CHUU, respectively. ROIs are sized according to the magnitude of ROI effect. Lower FC in the CPHIV group was predominantly from the left PAC and right CN/SOC to hippocampal subregions. Higher FC was predominantly from the bilateral MGN, left IC, right PAC to basal ganglia regions (putamen, pallidum and caudate) as well as to frontal lobe regions. (L/R)_parstr – parstriangularis, (L/R)_ pars – parsorbitalis, (L/R)_parso – parsopercularis, (L/R)_rMFC – rostral middle-frontal cortex, (L/R)_SFC – superior frontal cortex, (L/R)_cMFC – caudal middle-frontal cortex, (L/R)_precentral – precentral gyrus, (L/R)_paracentral – paracentral gyrus, (L/R)_rACC – rostral anterior cingulate cortex, (L/R)_cACC – caudal anterior cingulate cortex, (L/R)_PCC – posterior cingulate cortex, (L/R)_Isth – isthmus, (L/R)_postcentral – postcentral gyrus, (L/R)_SPC – superior parietal cortex, (L/R)_IPC – inferior parietal cortex, (L/R)_pericalc – pericalcarine, (L/R)_LOC – lateral occipital cortex, (L/R)_lingual – lingual gyrus, (L/R)_bankssts – banks of superior temporal sulcus, (L/R)_PAC – primary auditory cortex, (L/R)_Anterior – anterior thalamic nucleus, (L/R)_Medio_Dorsal – medio-dorsal thalamic nucleus, (L/R)_MGN – medial geniculate nucleus, (L/R)_VLD – ventral latero-dorsal thalamic nucleus, (L/R)_CLLPM_Pulvinar – central latero-lateral posterior medial pulvinar, (L/R)_V_Anterior – ventral anterior thalamic nucleus, (L/R)_Hippo – hippocampus, (L/R)_Hippo_Pres – hippocampus presubiculum, (L/R)_Hippo_CA1,3,4 – hippocampus cornu ammonis 1,3,4, (L/R)_Hippo_Mol_layer_HP – hippocampus molecular layer, (L/R)_Hippo_Tail – hippocampus tail, (L/R)_IC – inferior colliculus, (L/R)_SOC_CN – cochlear nucleus/superior olivary complex. L/R – left/right.

#### ROI pair effects

Compared to CHUU, the CPHIV showed strong evidence of lower FC between the left PAC and bilateral lingual gyrus, bilateral hippocampus CA1, bilateral hippocampus CA3, left hippocampus, left hippocampus presubiculum, and the left hippocampus molecular layer. The right CN/SOC also showed lower FC to the same regions as the left PAC in the CPHIV group. Further, we found strong evidence of higher FC (P+<=0.05) in the CPHIV group between the right PAC and right putamen, bilateral MGN and right putamen, right pallidum, right parsopercularis, bilateral IC to the right putamen, and left CN/SOC to the right putamen (Figure 3). ROI pair effect estimates for all connections to auditory regions can be found in Table 4 in Appendix C:.

### Functional network results

RBA showed strong evidence (P+ >= 0.95) of lower nodal degree of the left IC in the CPHIV group compared to CHUU. The right IC showed evidence of lower nodal degree in the same group (P+=0.94). Additionally, we found strong evidence (P+ >= 0.95) of lower nodal local efficiency of the right CN/SOC, left pulvinar and right IC in the CPHIV group compared to CHUU. The left IC showed evidence of lower nodal local efficiency in the same group (P+=0.91) (Figure 4). The nodal effect estimates for all graph measures can be found here Appendix D: Summary statistics of functional network nodal measures, CHUU-CPHIV.

**Figure 4:**
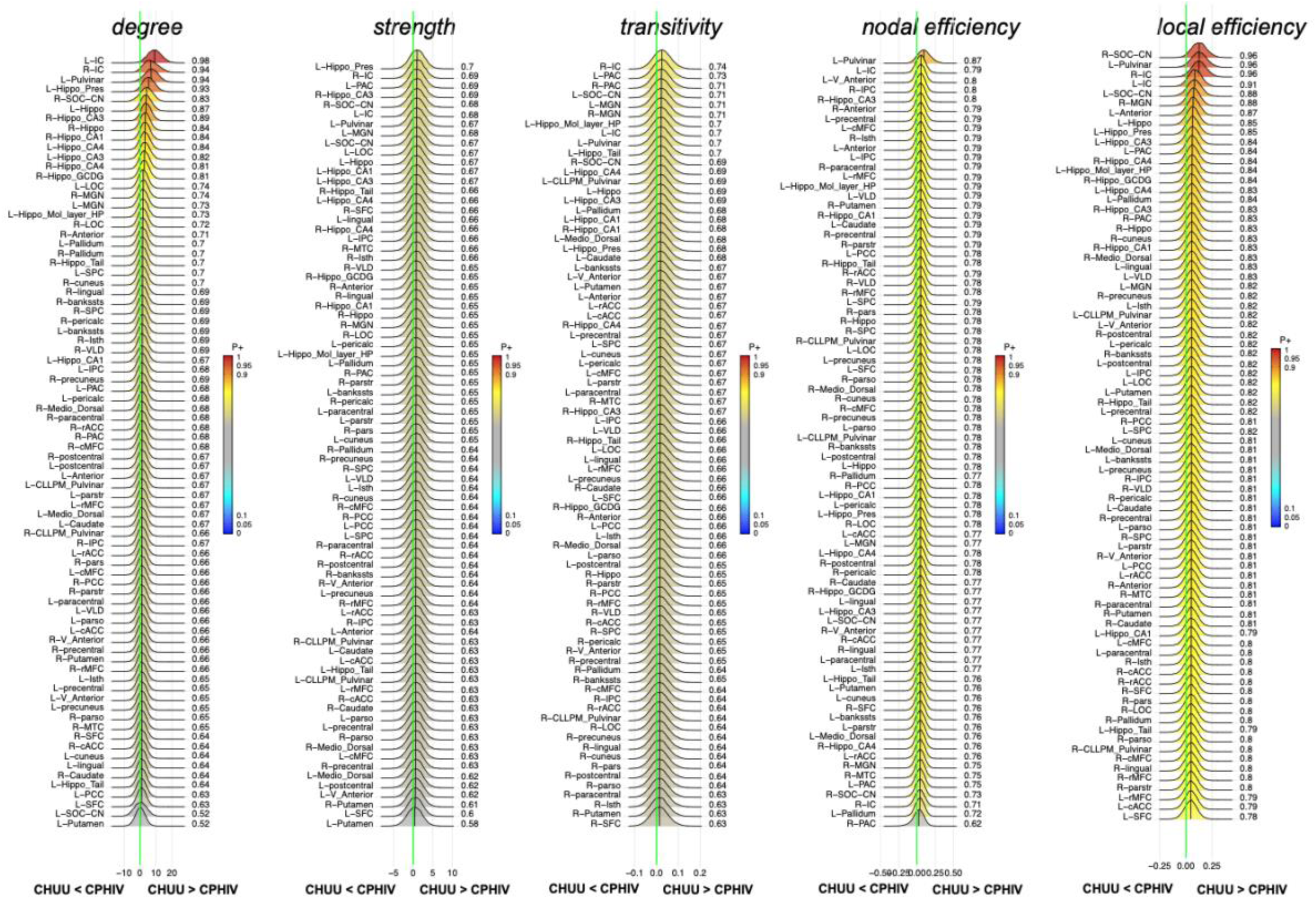
Posterior probability distribution plots for the effect magnitude of the comparison CHUU-CPHIV difference between mean graph measures. The graph measures include nodal degree, strength and transitivity, and both nodal and local efficiency. The green vertical line marks the location of no effect. L/R)_parstr – parstriangularis, (L/R)_ pars – parsorbitalis, (L/R)_parso – parsopercularis, (L/R)_rMFC – rostral middle-frontal cortex, (L/R)_SFC – superior frontal cortex, (L/R)_cMFC – caudal middle-frontal cortex, (L/R)_precentral – precentral gyrus, (L/R)_paracentral – paracentral gyrus, (L/R)_rACC – rostral anterior cingulate cortex, (L/R)_cACC – caudal anterior cingulate cortex, (L/R)_PCC – posterior cingulate cortex, (L/R)_Isth – isthmus, (L/R)_postcentral – postcentral gyrus, (L/R)_SPC – superior parietal cortex, (L/R)_IPC – inferior parietal cortex, (L/R)_pericalc – pericalcarine, (L/R)_LOC – lateral occipital cortex, (L/R)_lingual – lingual gyrus, (L/R)_bankssts – banks of superior temporal sulcus, (L/R)_PAC – primary auditory cortex, (L/R)_Anterior – anterior thalamic nucleus, (L/R)_Medio_Dorsal – medio-dorsal thalamic nucleus, (L/R)_MGN – medial geniculate nucleus, (L/R)_VLD – ventral latero-dorsal thalamic nucleus, (L/R)_CLLPM_Pulvinar – central latero-lateral posterior medial pulvinar, (L/R)_V_Anterior – ventral anterior thalamic nucleus, (L/R)_Hippo – hippocampus, (L/R)_Hippo_Pres – hippocampus presubiculum, (L/R)_Hippo_CA1,3,4 – hippocampus cornu ammonis 1,3,4, (L/R)_Hippo_Mol_layer_HP – hippocampus molecular layer, (L/R)_Hippo_Tail – hippocampus tail, (L/R)_IC – inferior colliculus, (L/R)_SOC_CN – cochlear nucleus/superior olivary complex. L/R – left/right.

### Associations between functional outcomes and auditory/language-related scores

Among the sample as a whole, we found no associations between FC and the auditory/language-related scores after correcting for multiple comparisons. However, among CHUU, we found negative associations between standardized Delayed Recall scores and nodal efficiency in the right middle temporal cortex (MTC), as well as a significantly positive association between standardized Delayed Recall and nodal efficiency in the left superior frontal cortex (SFC) (FDR q < 0.05) (Figure 5). These associations were not seen in CPHIV.

**Figure 5:**
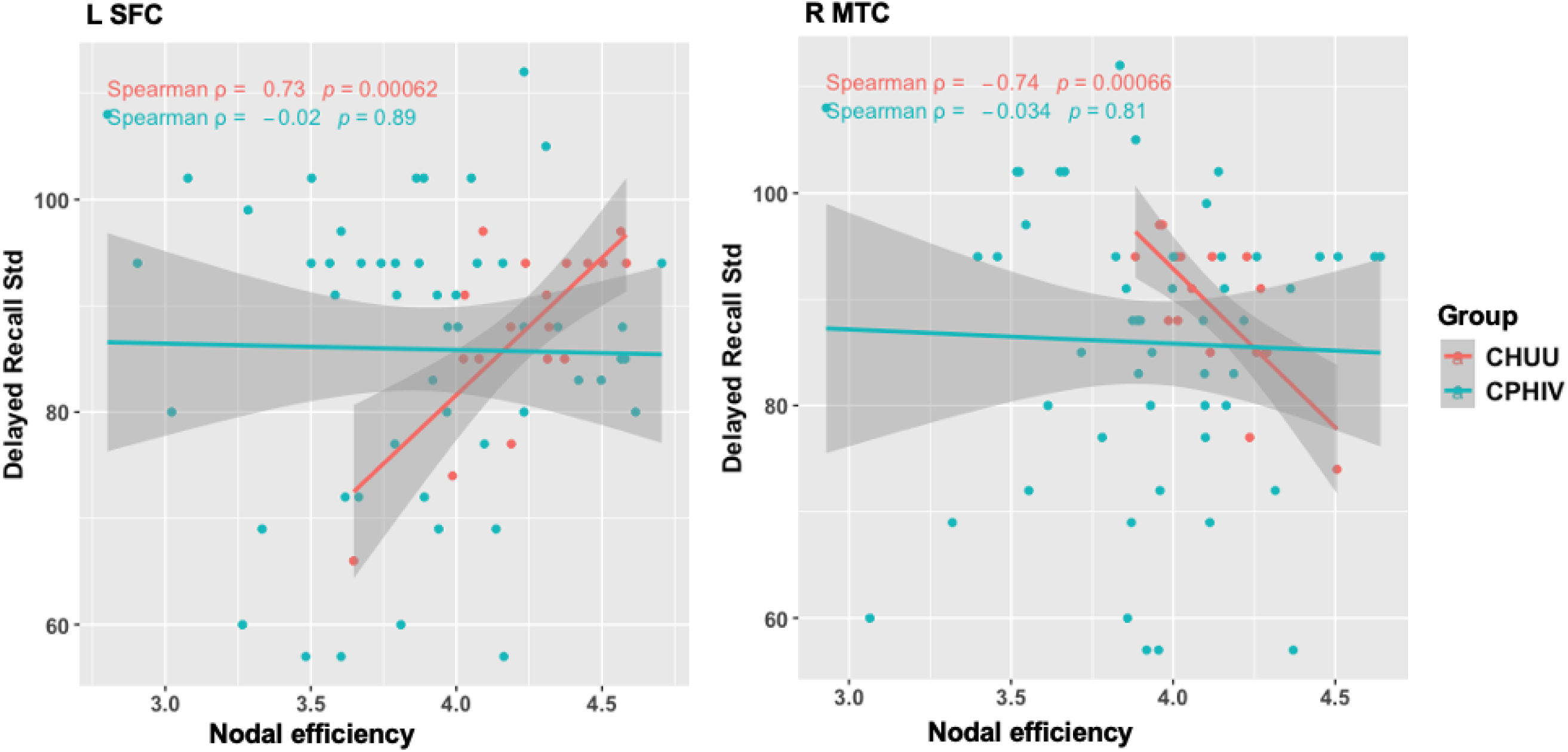
Associations of nodal efficiency in the left superior frontal cortex (SFC) and right middle temporal cortex (MTC) with Delayed Recall test scores assessed on the Kaufman Assessment Battery for Children – Second Edition (KABC-II). Std = standardized. L – right, R – right.

### Associations between functional outcomes and immune health markers

We found no significant associations between functional outcomes and immune health makers after multiple comparison correction.

## Discussion

The current study investigated the functional integrity of the CAS in relation to HIV in an adolescent cohort. CPHIV demonstrated reduced FC to the PAC, from hippocampal areas and the lingual gyrus which are involved in limbic and visual functions respectively. In addition, we reported evidence of increased FC to the bilateral MGN, with connections to the parsopercularis, an important region in speech production, most affected. Our analysis shows HIV infection influences the relationships between auditory structures and non-auditory systems, including the auditory-visual and auditory-limbic, in 11-year old children.

Both ROI and region pair analysis identified lower FC related to HIV infection in the left PAC, bilateral lingual gyrus and bilateral hippocampal regions. Altered FC of the left PAC is noteworthy as it is responsible for language processing, which displays hemisphere lateralisation (Bernal & Altman, 2012; Binder et al., 1996; Olulade et al., 2020). In fact, language lateralisation increases significantly between childhood and adolescence (Szaflarski et al., 2006). The majority of our cohort was right-handed, suggesting left hemisphere dominance for language processing. However, even though alterations to the left PAC suggest direct effects on language processing we did not find significant associations between FC to the region and language related neurodevelopmental outcomes. Instead, we found evidence of reduced FC between the left PAC and non-auditory regions including the lingual gyri and hippocampal regions.

The link between the hippocampus, a limbic system region located in the medial part of the temporal lobe, and the auditory system has been previously reported (Kraus & Canlon, 2012; Zhang et al., 2022). The hippocampus is involved in the formation of long-term auditory memories (Nyhus & Curran, 2010; Watanabe et al., 2008) as well as the comprehension of spoken language (Davis & Johnsrude, 2003; van de Ven et al., 2020).

Cortico-limbic FC is particularly vulnerable in the adolescent brain as cognition-emotion interactions are being refined (Simmonds et al., 2014). Lower FC between hippocampal regions and PAC point to possible interruptions of these relationships in the presence of HIV. Further, hippocampal neuropathology has been linked to hearing impairment (Park et al., 2018; Uchida et al., 2018) and associated with poor memory (Squire et al., 2001). Notably, CPHIV in this cohort at age 7 were found to have lower hippocampal volumes (Nwosu et al., 2018).

On the other hand, the lingual gyrus is part of the visual network and involved in visual memory and visual language processing (Palejwala et al., 2021). We have previously reported reduced FC to the lingual gyrus in the same cohort at age 7 (Toich et al., 2018). The visual system relies heavily on auditory input for efficient and precise perception of the environment around us (Gori et al., 2012; Yu et al., 2018). Additionally, adult-like visual-auditory integration only reaches its peak in late-childhood making it at risk to disruptions in young children (Gori et al., 2012). Our results point to a vulnerability of the auditory and visual memory system in the presence of HIV.

The right CN/SOC did not show evidence of altered region based FC, however its connections to hippocampal regions and the bilateral lingual gyrus demonstrated lower FC in the CPHIV group compared to CHUU. The CN and SOC are crucial auditory processing structures. The CN is the initial site of subcortical auditory processing (Hackett, 2015), where auditory signals are enhanced and sent to the SOC. In the SOC, the signals are filtered and a crossing of signals from the contralateral SOC takes place (Peterson et al., 2021). Even at this level, the CAS is interacting with other systems for cognitive processing of attentional and motor actions (Stein & Stanford, 2008) to facilitate one’s behavioural or emotional response to auditory stimulation. These results echo alterations in auditory-limbic and auditory-visual FC. In addition, graph theory analysis revealed lower nodal degree and nodal local efficiency in the CN/SOC. The local efficiency of a node is the efficiency of the graph formed by the node’s neighbours (direct edges) in its absence. Unlike nodal efficiency, which reports the integration of a node within the entire network, local efficiency measures how efficiently the neighbours can communicate in the event of damage to the node (Fornito et al., 2016). This points to reduced integration among the immediate neighbours of both the CN and SOC in the presence of HIV.

We observed several connections between CAS regions and non-auditory regions showing evidence of **higher** FC in the CPHIV group compared to CHUU. The non-auditory regions were distributed among basal ganglia, thalamic nuclei and frontal areas. The basal ganglia are of particular interest as they show alterations in CPHIV in childhood across imaging modalities (Hoare et al., 2014; Mbugua et al., 2016; Pajkrt et al., 2016; Robertson et al., 2018). Within this cohort, we’ve reported HIV-related alterations and associations in the basal ganglia at earlier ages. At ages 5 and 7, putamen volumes were larger and smaller in the CPHIV group, respectively, compared to controls (Nwosu et al., 2018; Randall et al., 2017). At age 7 years, we reported a negative association between FC and CD4 count in infancy among CPHIV pointing to higher FC linked with a weaker immune system (Toich et al., 2018). A longitudinal study from ages 5 to 11 years of neurometabolism in our cohort reported elevated choline levels in the basal ganglia, suggesting persistent inflammation (van Biljon et al., 2021). Our current results at age 11 suggest overcompensation from all levels of the CAS to the basal ganglia, which may be related to the volumetric and neurometabolic differences observed at earlier ages. Basal ganglia are broadly involved in motor control and the putamen interacts with temporal regions to initiate phonological processes (Lanciego et al., 2012).

The bilateral MGN also showed higher FC to the parsopercularis in the presence of HIV. The parsopercularis, a region part of the inferior frontal gyrus, plays a role in language and phonological processing. Interestingly altered FC in the inferior frontal gyrus has been reported previously in the same cohort, although FC was lower in CPHIV in the analysis at age 7 compared to CHUU (Toich et al., 2018). Altogether our results suggest increased interaction between the CAS and regions in other brain systems including limbic-motor, visual and language areas in the presence of HIV. Interestingly children with hearing impairments demonstrate a similar pattern of overcompensation in the auditory network, motor-limbic and visual areas (Luan et al., 2019; Shi et al., 2016; Wang et al., 2021).

Besides the reduced nodal integrity of the CN/SOC, our functional network analysis also revealed lower nodal degree and local efficiency in the IC of the CPHIV group as compared to CHUU. This result is unexpected as the IC did not show any evidence of FC alterations between groups. The reduced nodal integrity observed in the current analysis may be related to the lower degree observed in this node as well as the previously discussed CN/SOC. The nodal degree reflects the density of connections to the node, with a low degree reflecting nodes that are not as highly connected with other regions. It is worth noting that both these regions are in the brainstem where the measured BOLD signal isn’t high. Indeed, they were the structures with the lowest number of connections in Figure 1B.

Additionally, we found a positive association between nodal efficiency of the SFC and the Delayed Recall test as well as a negative association between nodal efficiency of the MTG and Delayed Recall. Both associations were seen in the CHUU group but not in CPHIV. Delayed Recall is part of the KABC Learning subscale (Kaufman & Kaufman, 2014). More specifically Delayed Recall is a composite score of Atlantis delayed, where children are asked to point to pictures they would have learnt 20 minutes earlier, and Rebus delayed where the children are taught pictures that have a word that they have to put in a sentence and recall 20 minutes later. Though both the tests have visual cues they assess the child’s language processing ability in coming up with a sentence as well as their ability to store and retrieve memories. The SFC is recruited during tasks of delayed recognition (Yee et al., 2010) while the MTG is involved in phonologic and sematic language processing (Middlebrooks et al., 2016), both processes of which interact with the CAS to facilitate speech production for communicating (Maffei et al., 2015). In addition temporo-frontal regions show altered FC in CPHIV (Toich et al., 2018; Yadav et al., 2018). Our current results point to possible disruption in the relationships between temporo-frontal nodal integrity and neurodevelopment relating to working memory in CPHIV.

## Limitations and future work

The origin and interpretation of negative FC has been an area of debate (Chen et al., 2011; Goelman et al., 2014; Mohanty et al., 2020) and the role of negative correlations as edge weights is not yet fully understood. Nodes with strong positive correlations suggest that the nodes work together whereas strong negative correlations imply the nodes have an antagonistic relationship (Fornito et al., 2016). In the present study we used only positive weights (Fisher Z-value >0.1) as the measure of pairwise connectivity in the functional network. This not only removes weak edge weights that represent spurious connections and may alter network topology, it also simplifies the interpretation of FC to cooperation between nodes. However, this approach discards information about other functional interactions in the network, particularly the very high but negative correlations. Negative correlations may imply that the pairs of regions work together in an anti-correlated way (Keller et al., 2015).

In addition, the choice of a threshold of Fisher Z-value > 0.1 was arbitrary. The threshold choice was used to improve the number of pairwise connections between subcortical CAS regions deep in the brainstem and non-auditory ROIs. As seen in Figure 1B of the functional network matrix, most of the FC to the CAS was to the PAC, then to the MGN, less to the IC and even less to the CN/SOC in the brainstem. Using a range of adjacency matrices during network definition, for example Fisher Z-value > 0.1 with 0.05 increments, as opposed to the current network definition of Fisher Z-value > 0.1 may help observe if the current results persist.

While this work examines the functional auditory system and its associations with language measures, it does not include measures of hearing. Future work is needed to link the identified alterations with hearing-related outcomes. In addition, given the dynamic nature of the brain in adolescence, longitudinal analysis is needed to examine the long-term impact of the results presented.

## Conclusions

To our knowledge this is the first study to investigate the functional integrity of the CAS and its interactions with other systems in CPHIV. Our results show altered FC between the CAS and non-auditory regions involved in limbic, visual and motor processes in CPHIV. Furthermore, CPHIV show altered nodal topology of the auditory brainstem and midbrain, with nodal measures related to cognitive scores in the CHUU group only. These results provide insight on FC in CPHIV and point to risk of auditory and language disruptions based on CAS alterations.

## Supporting information

Appendix A: Automatically segmented regions of interest.

## Appendices

## Appendix A: Automatically segmented regions of interest

**Table 2:**
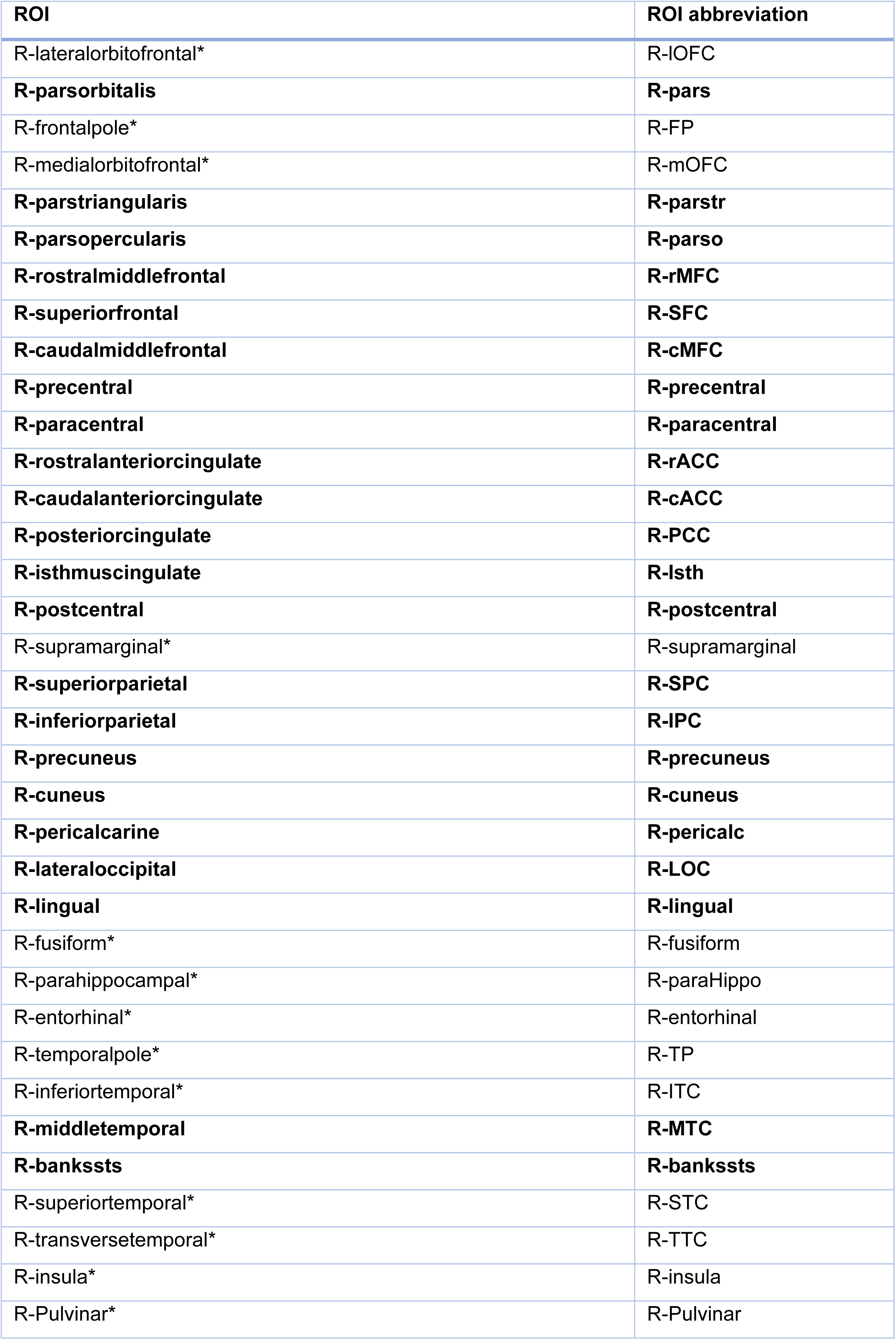
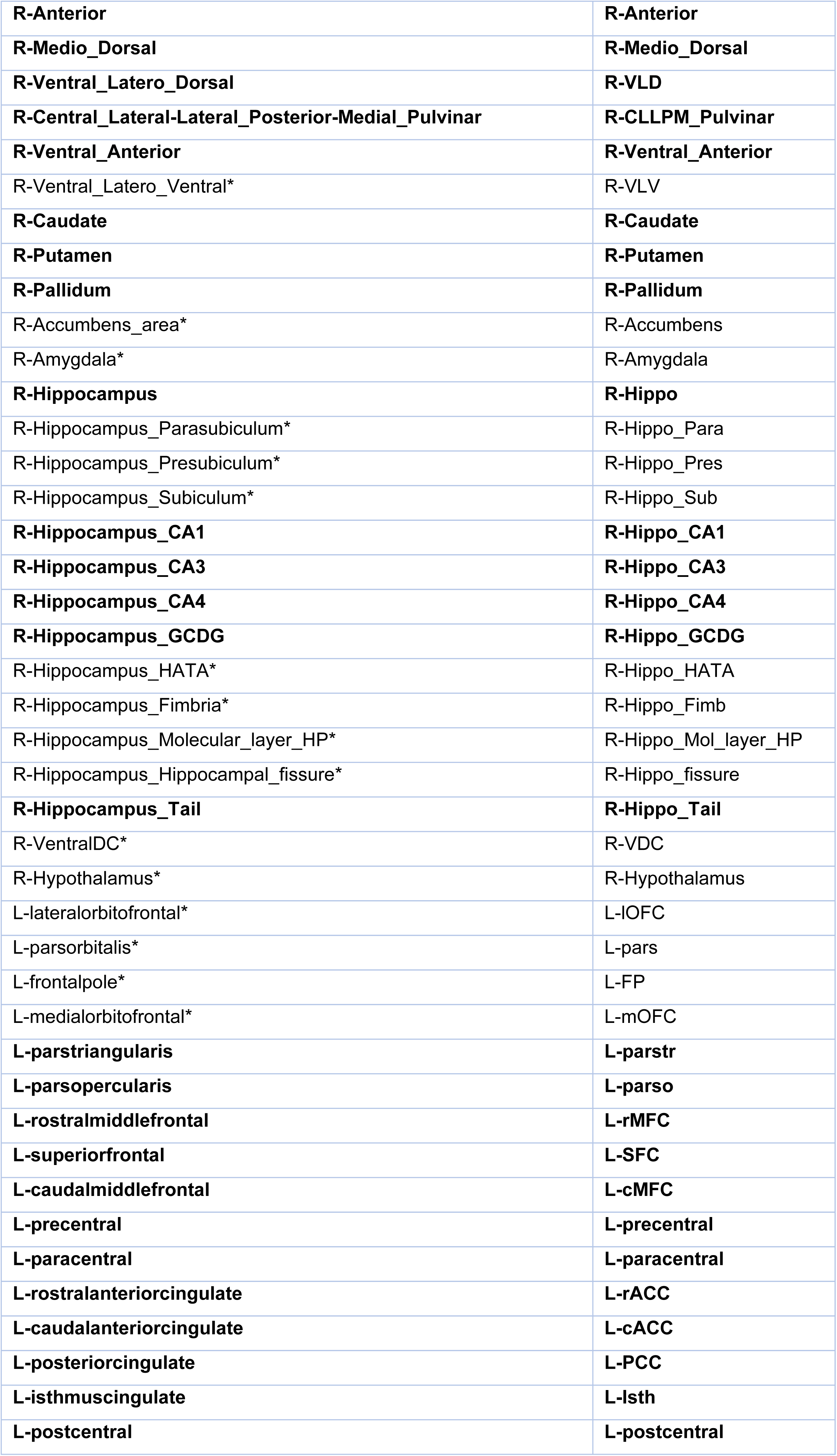
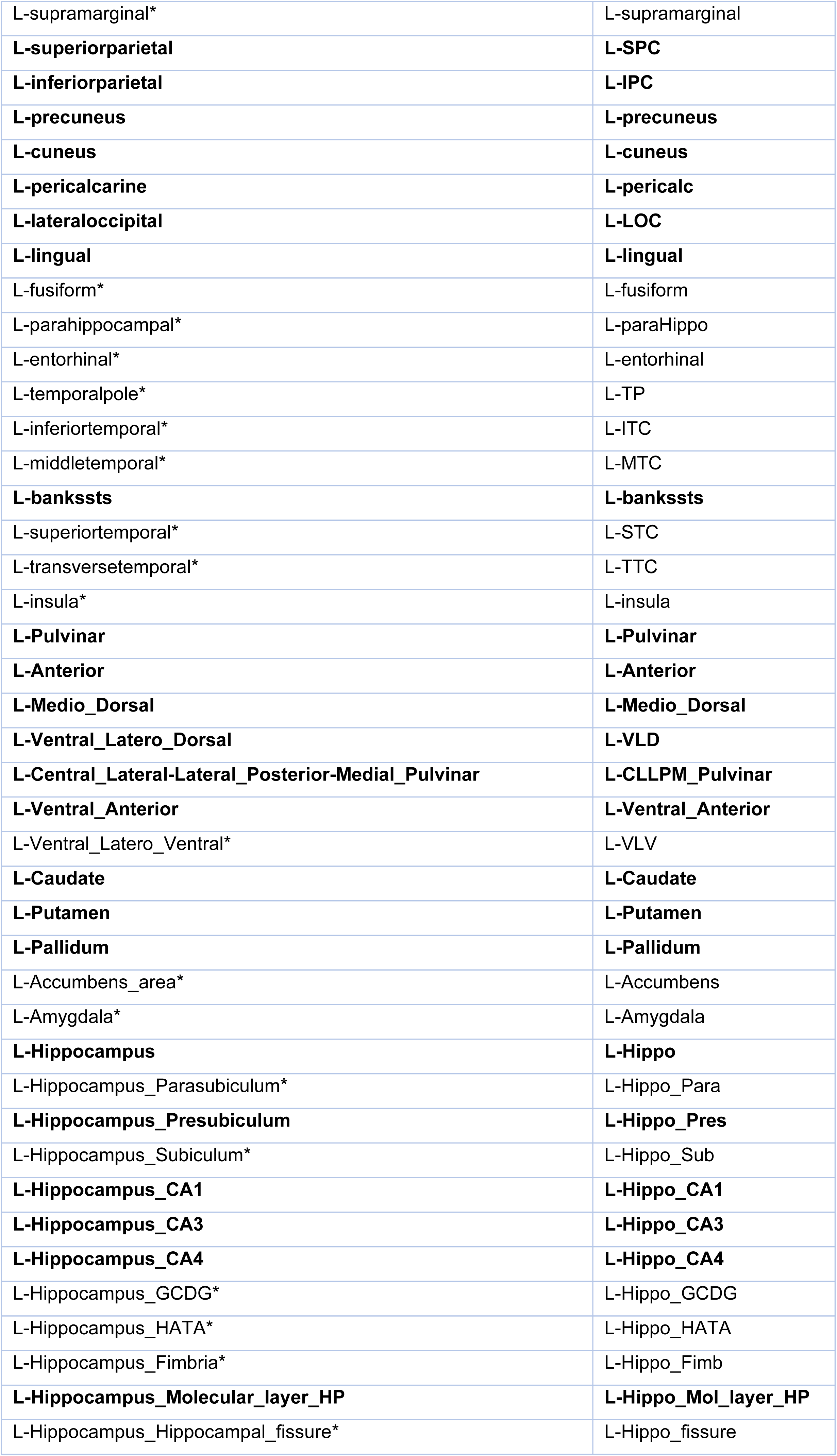
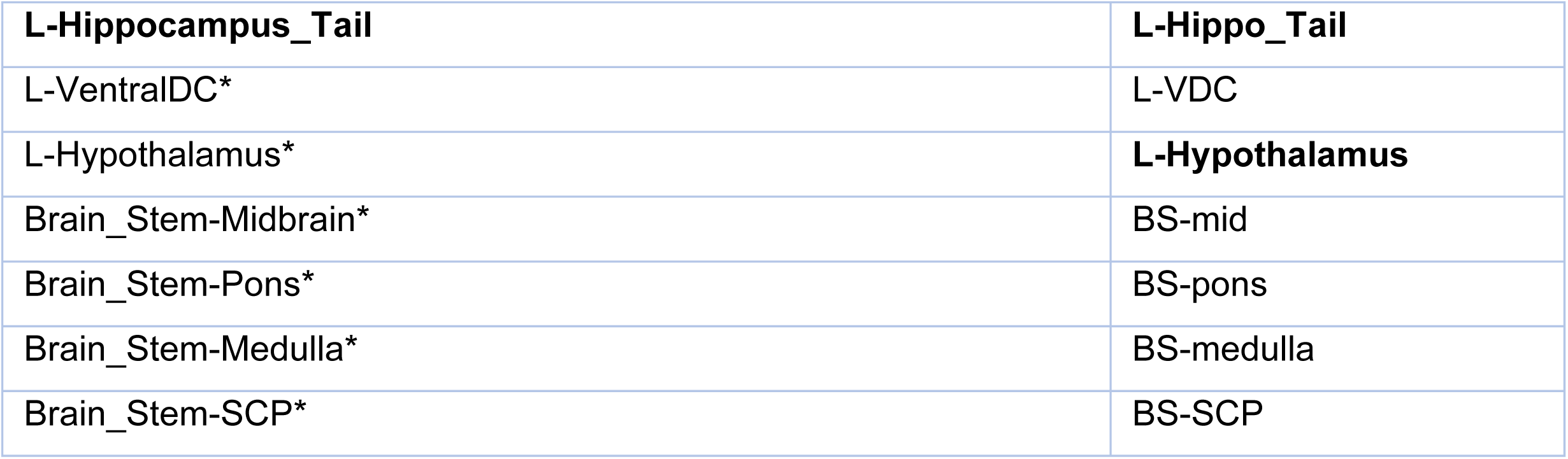
126 automatically segmented ROIs. Asterisks denote the 54 regions excluded due to overlapping with manually traced regions in at least participant. R – right, L – left.

## Appendix B: Summary information of model results for FC of CHUU vs CPHIV

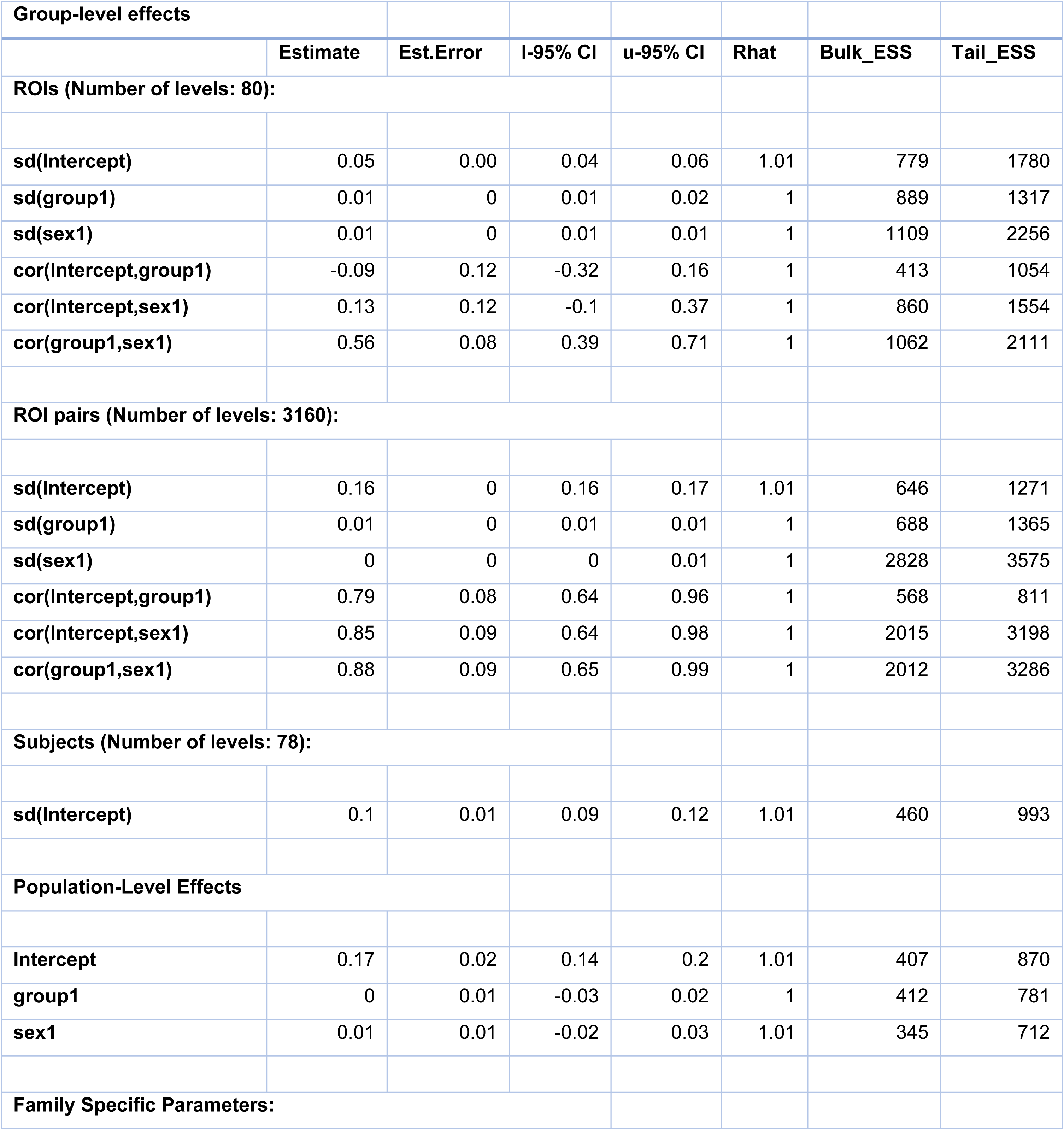
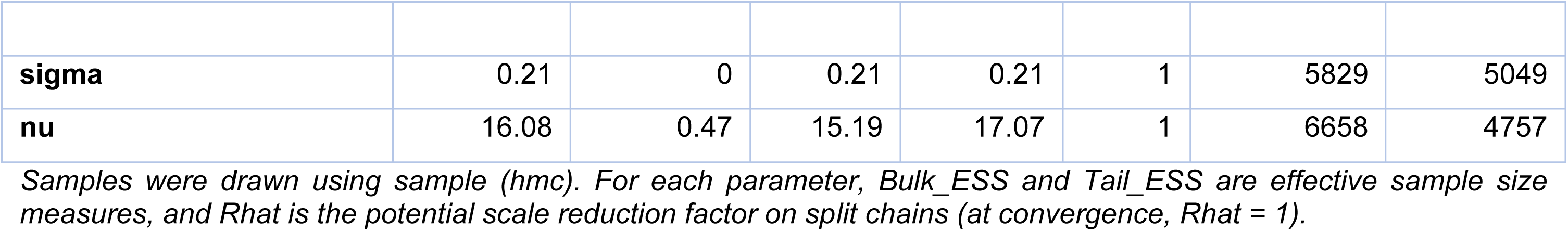

### Appendix C: Region and region pair effect estimates

**Table 3:**
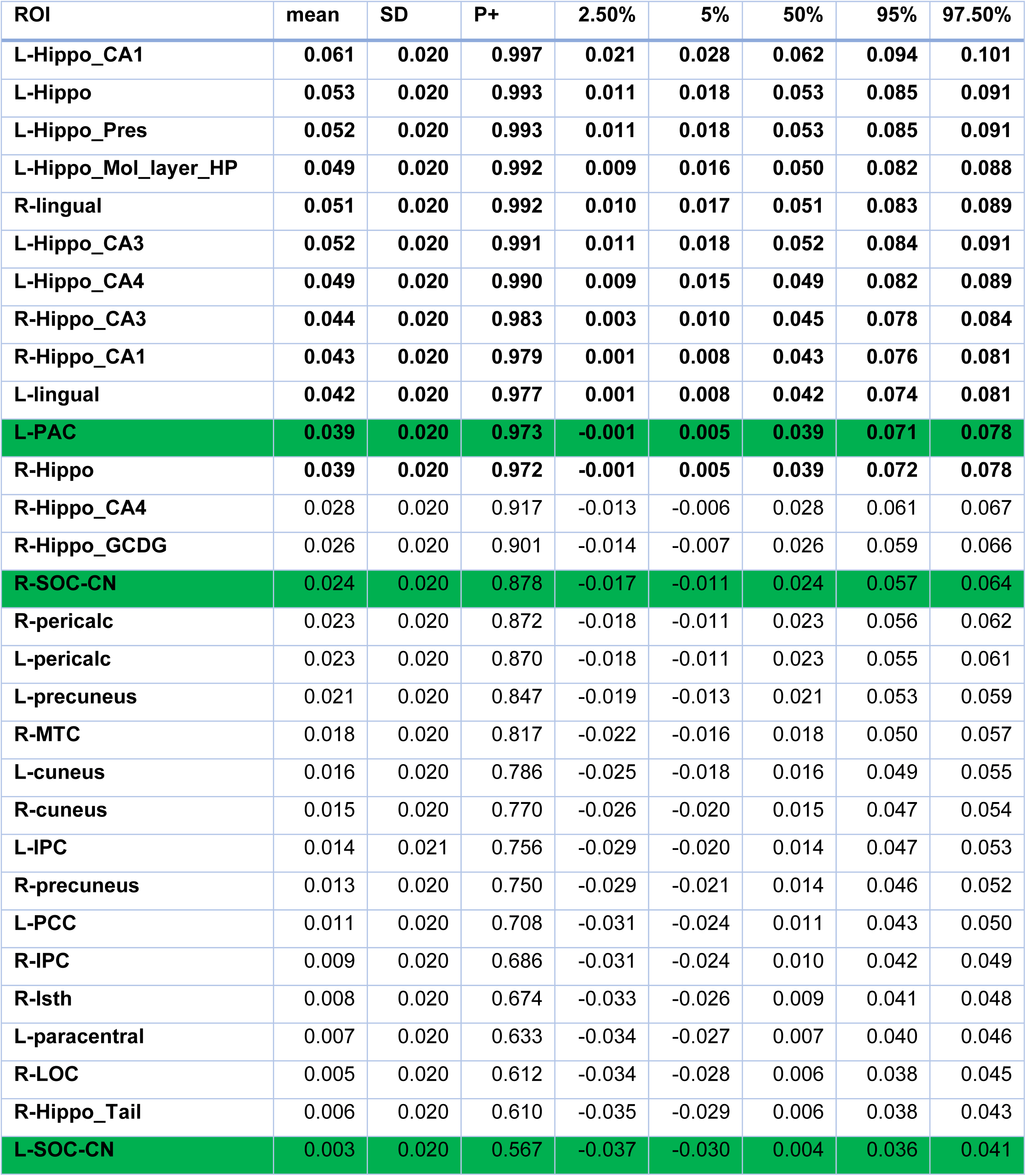
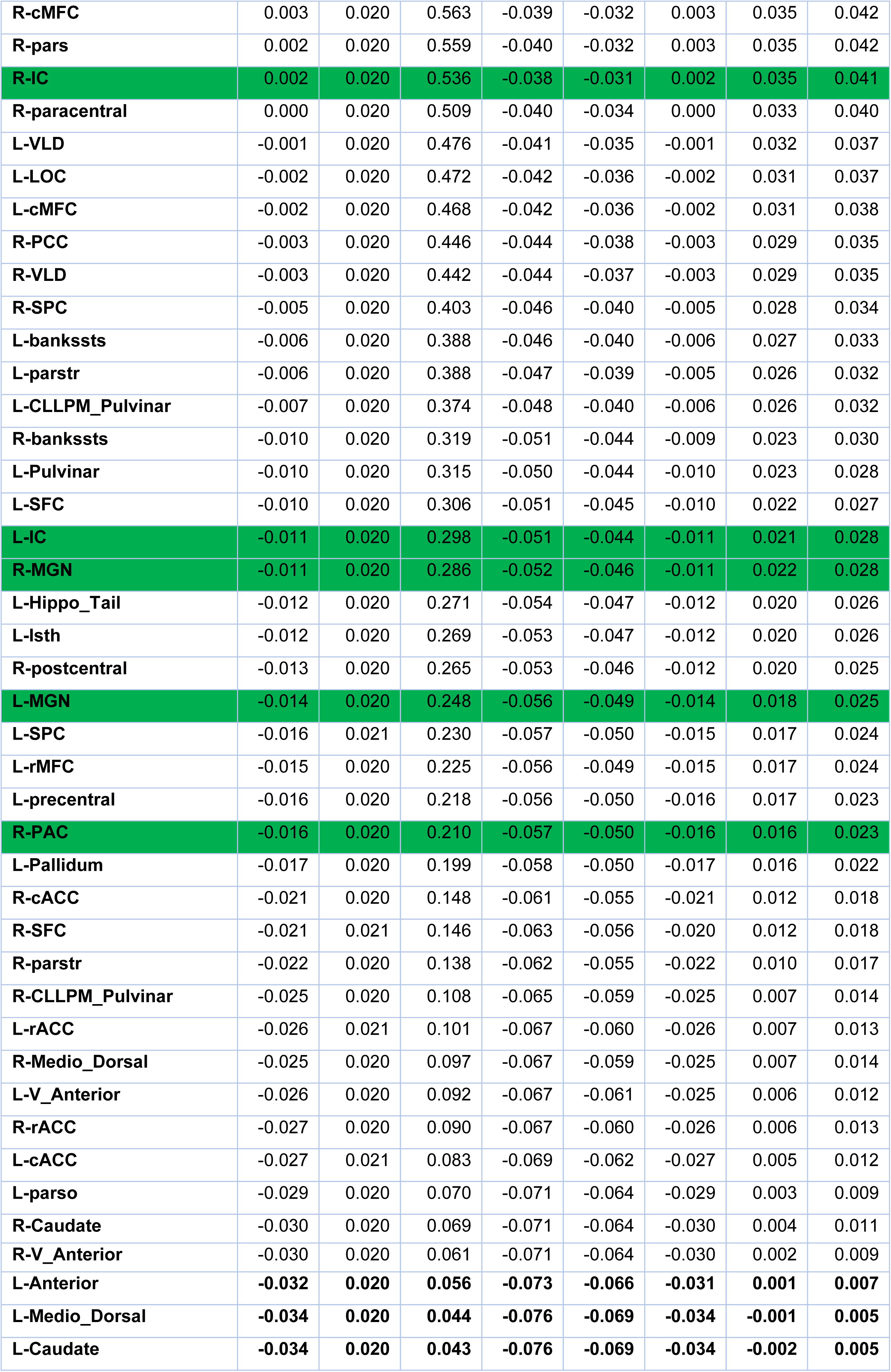
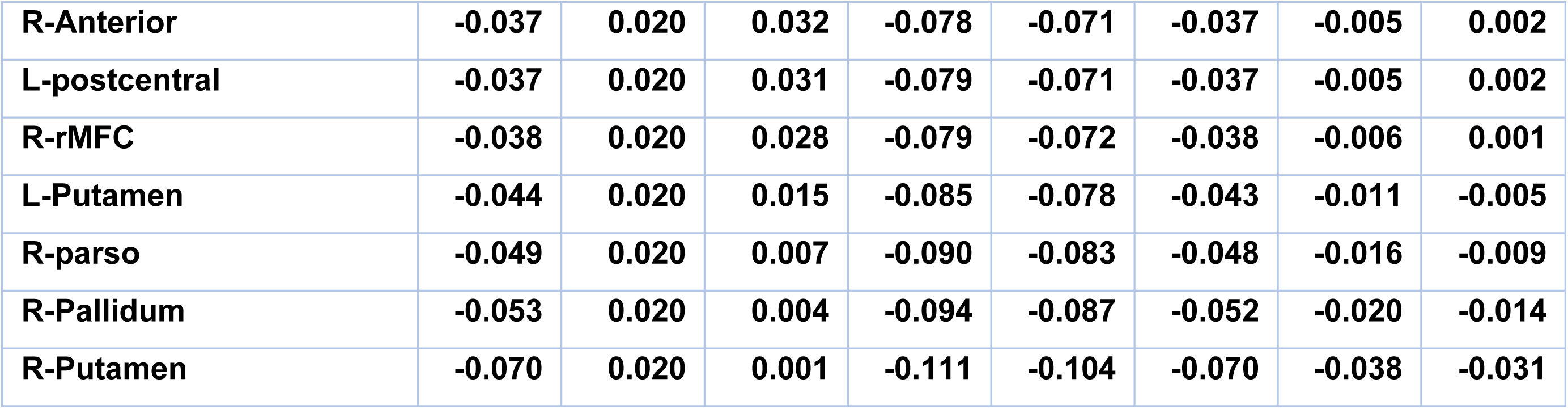
ROI effect estimates and their uncertainties for the comparison CHUU-CPHIV. The first 3 columns are difference between group mean FCs (Fisher’s Z-values), standard deviations, and the posterior probability of the effect being positive respectively. The preceding columns represent uncertainty intervals. The ROIs are organized in P+ descending order. Auditory ROI rows are marked in green and ROIs showing strong evidence of altered FC (P+ >= 0.95 | P+<= 0.05) are in bold.

**Table 4:**
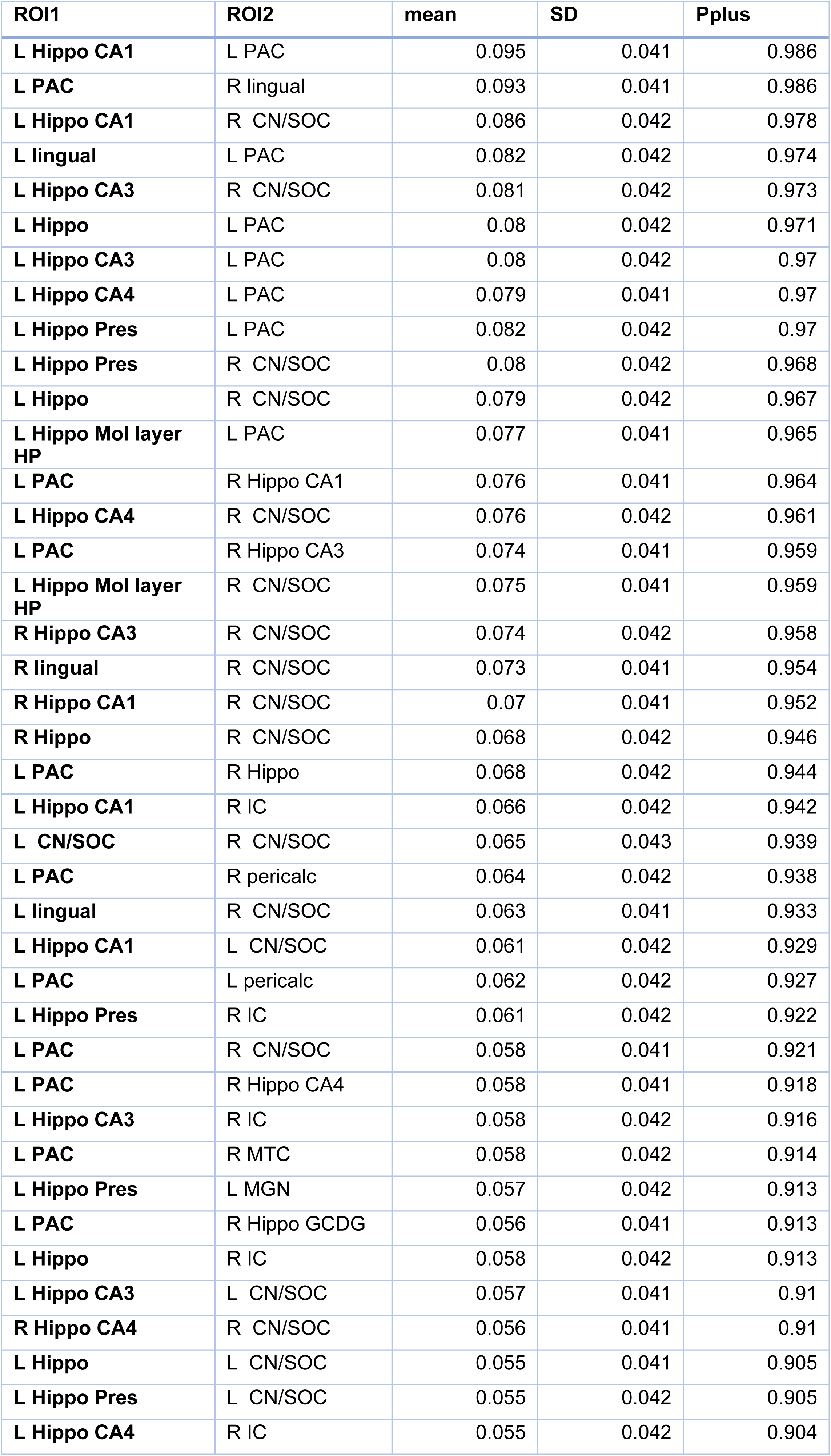
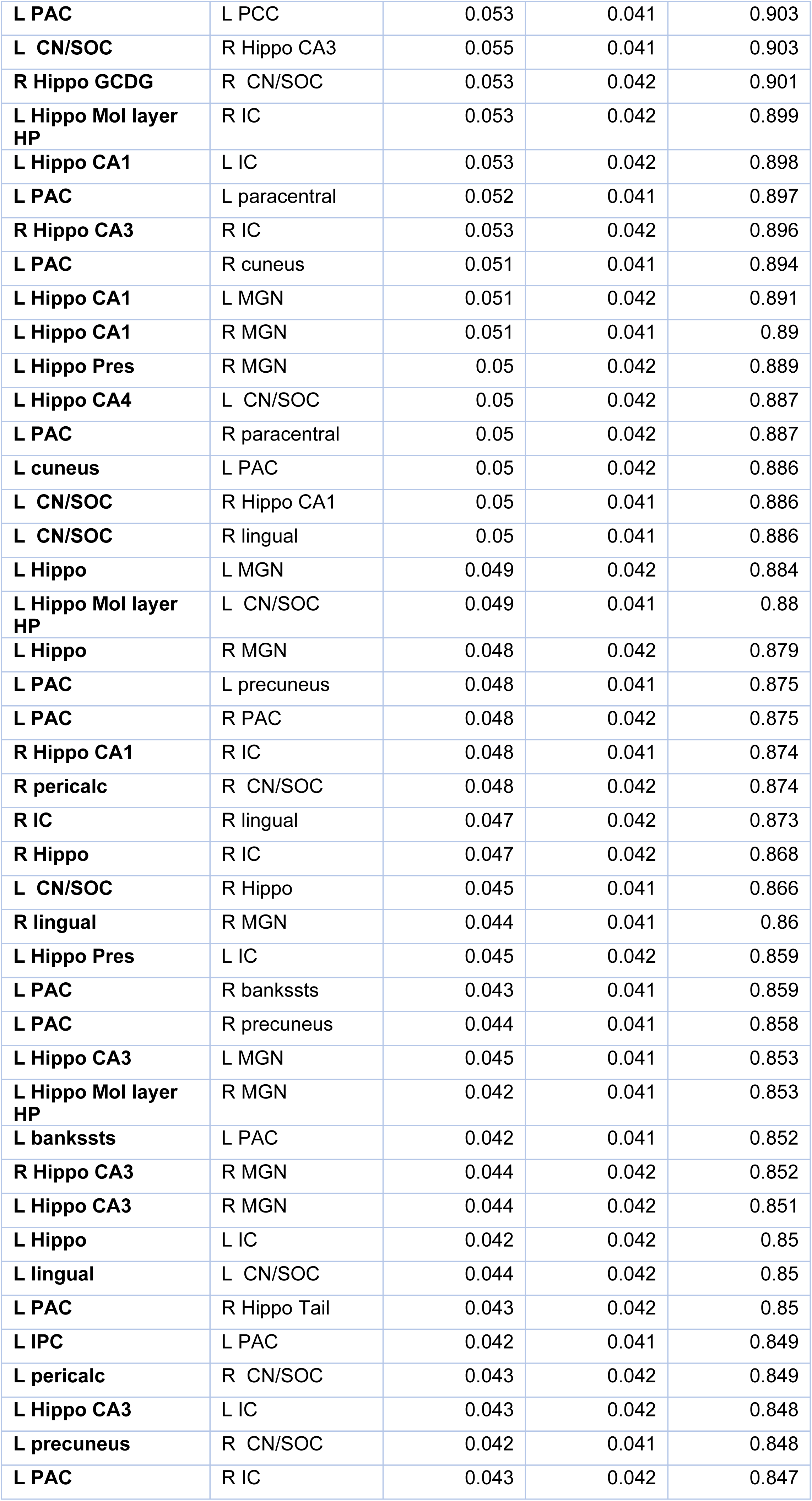
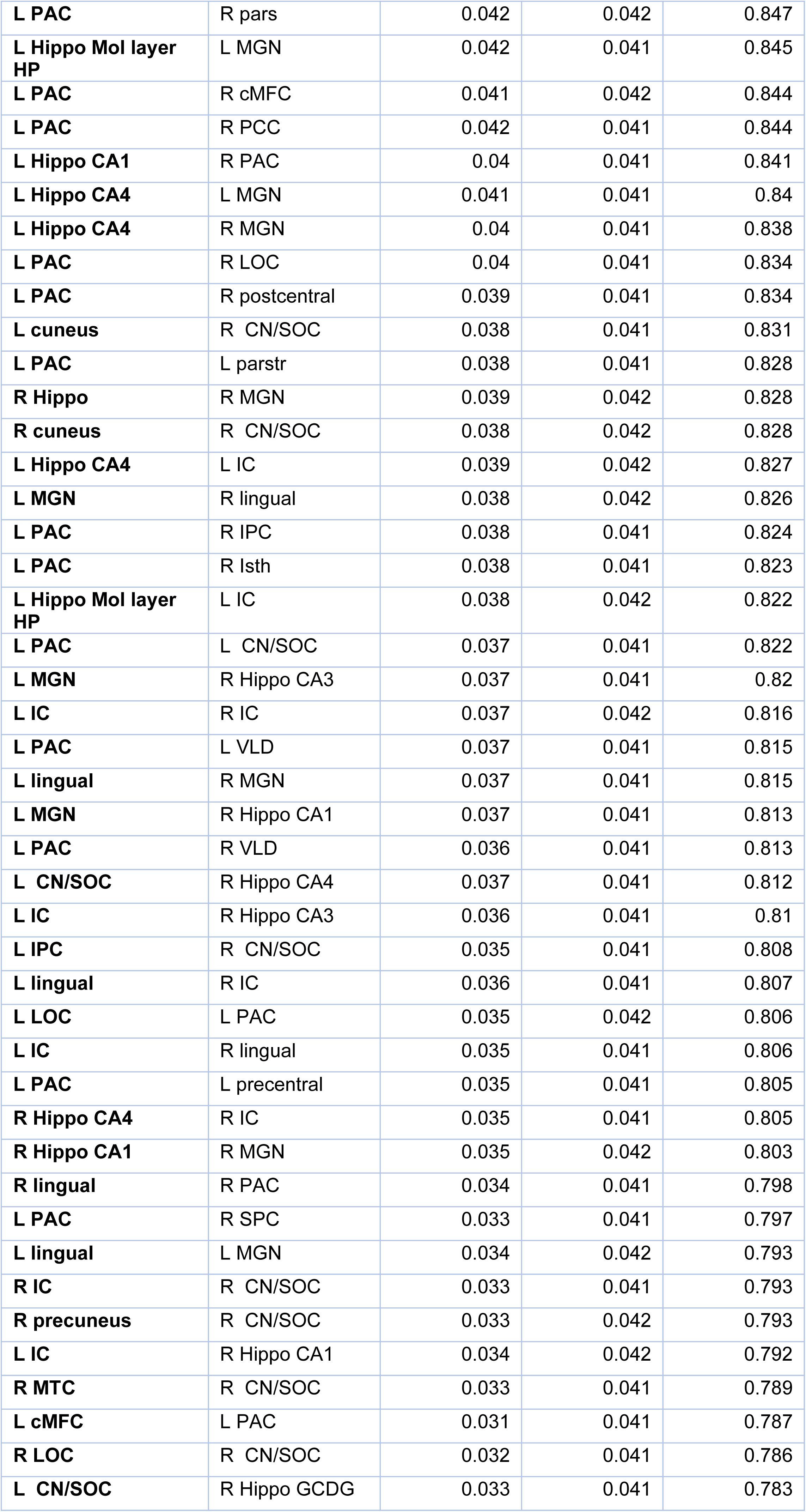
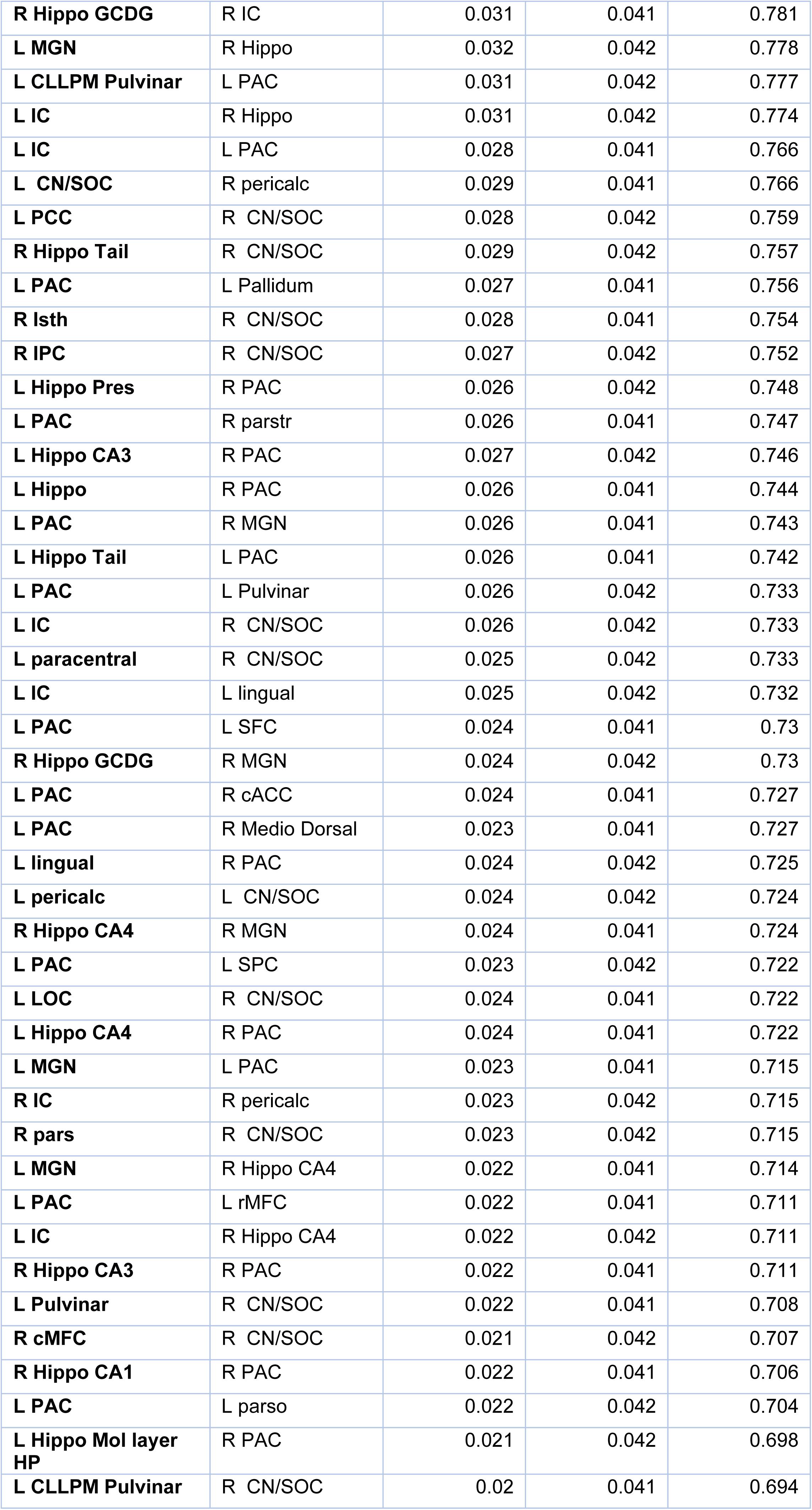
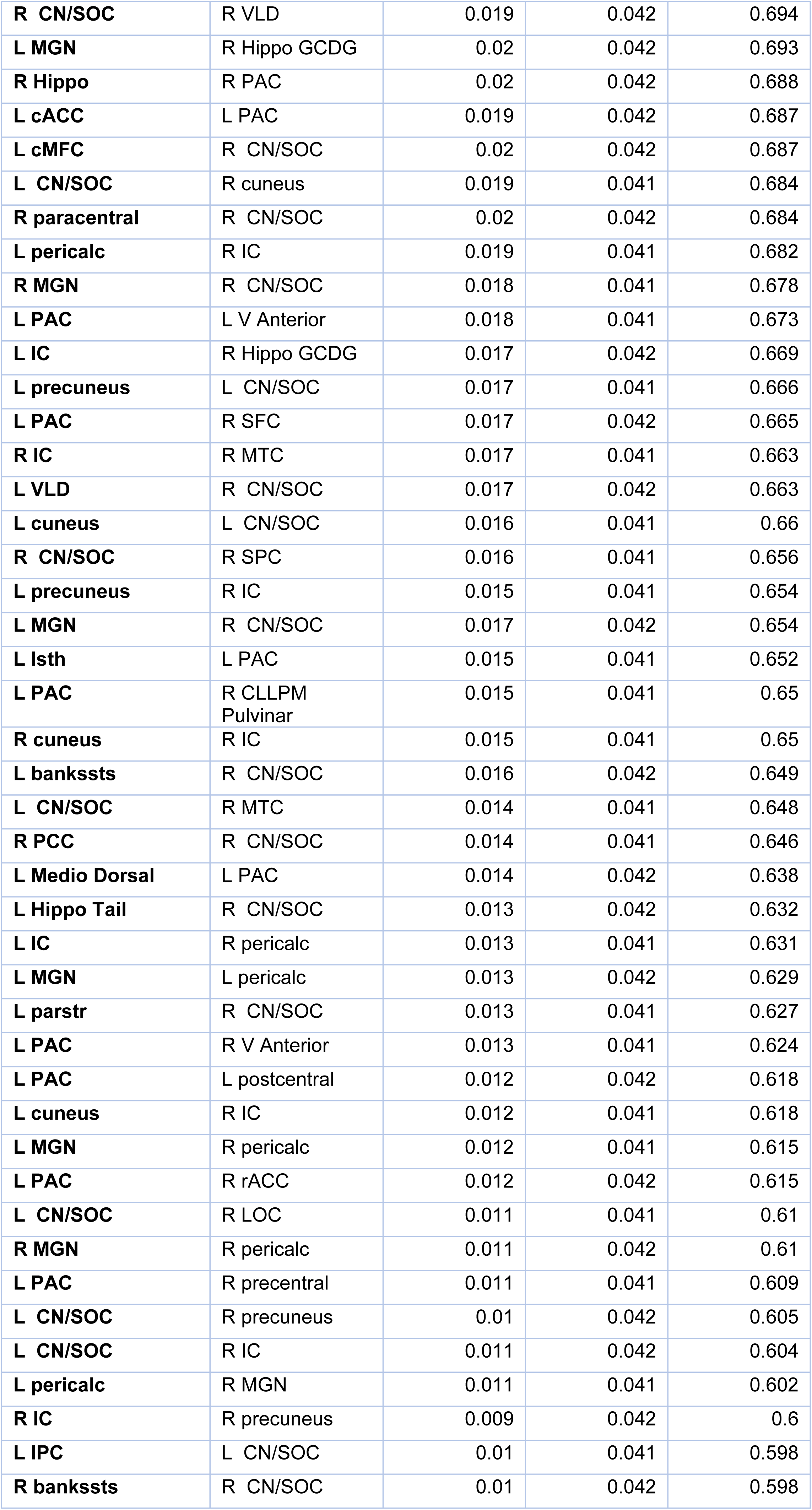
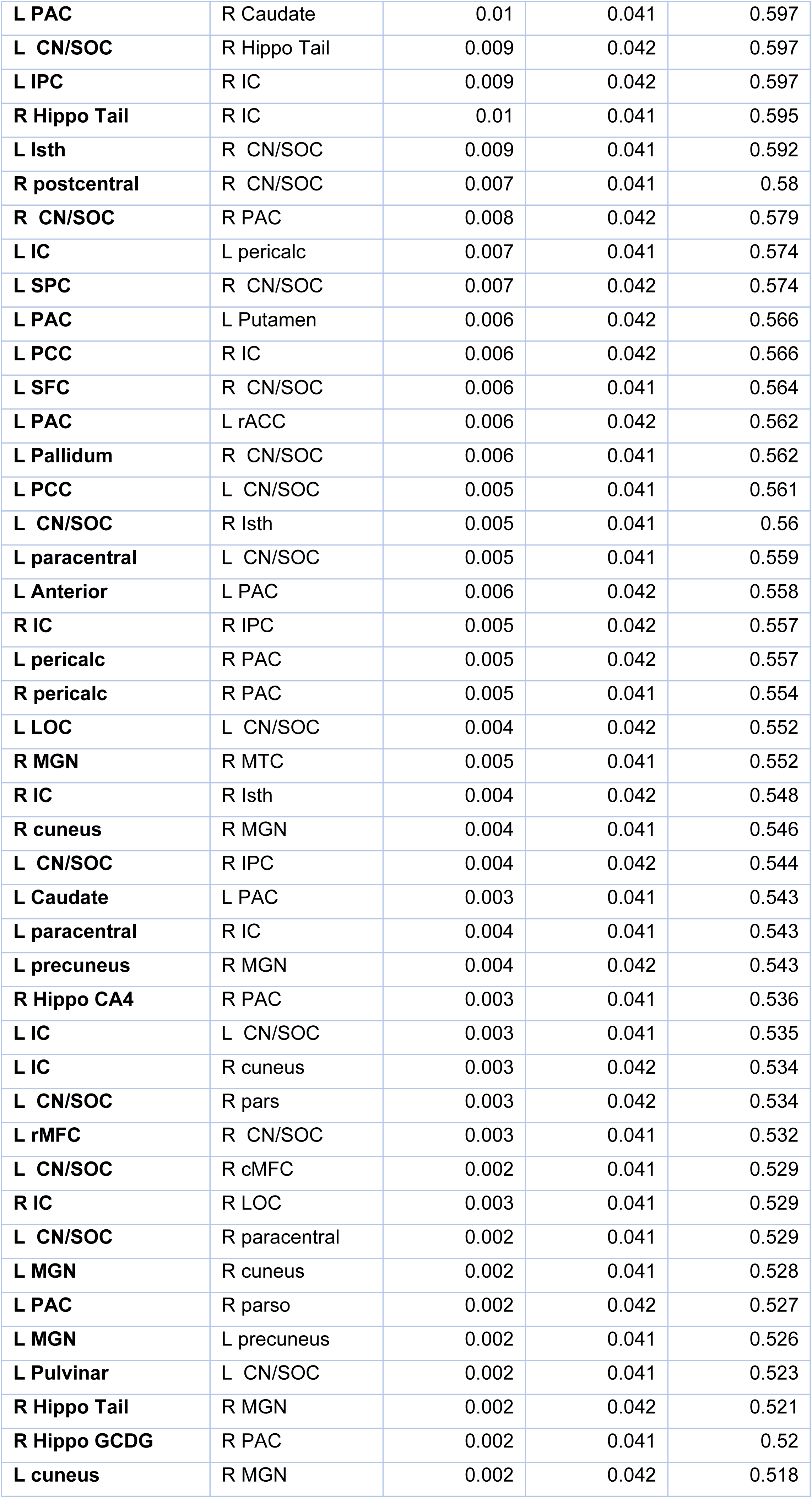
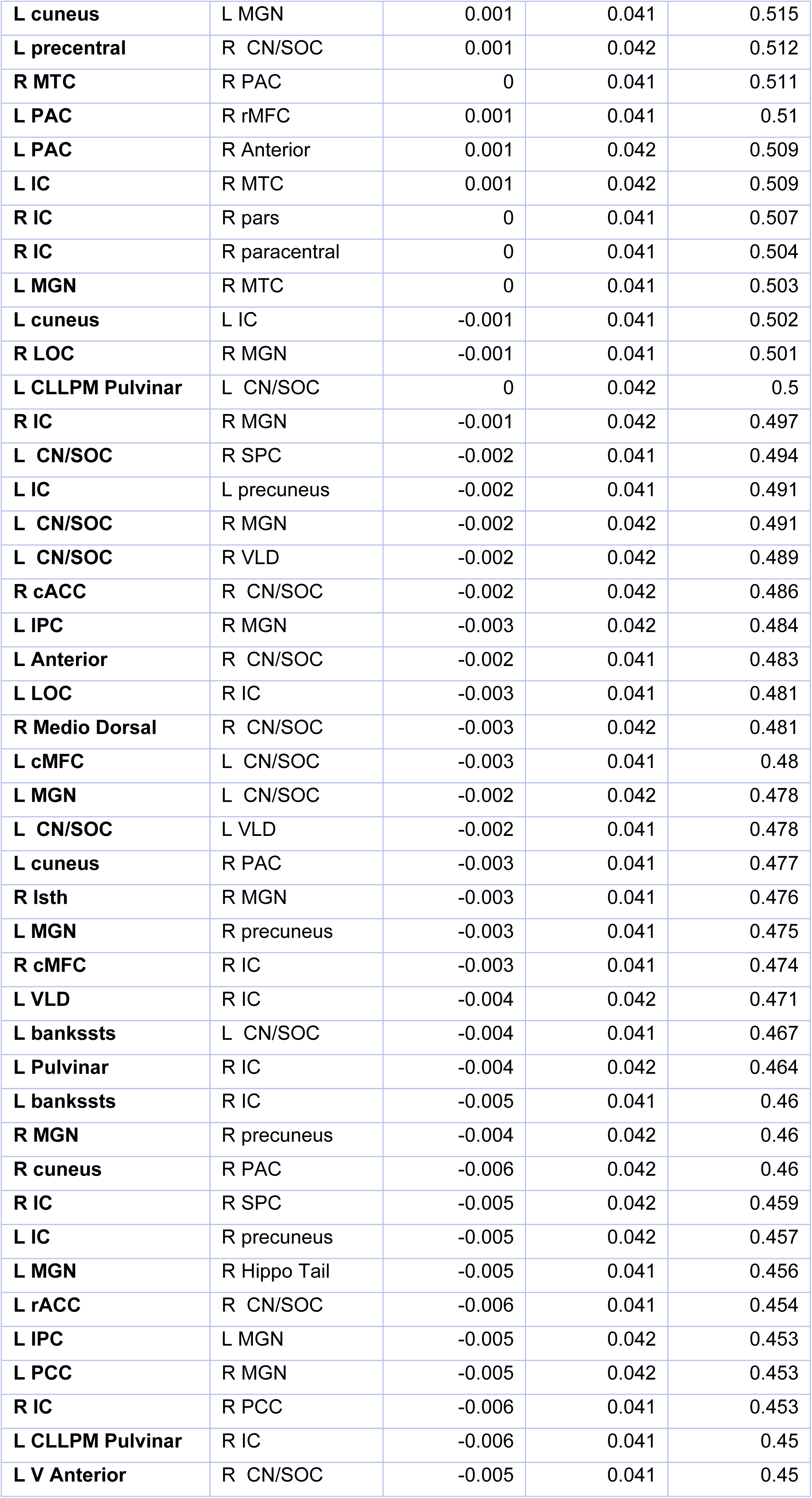
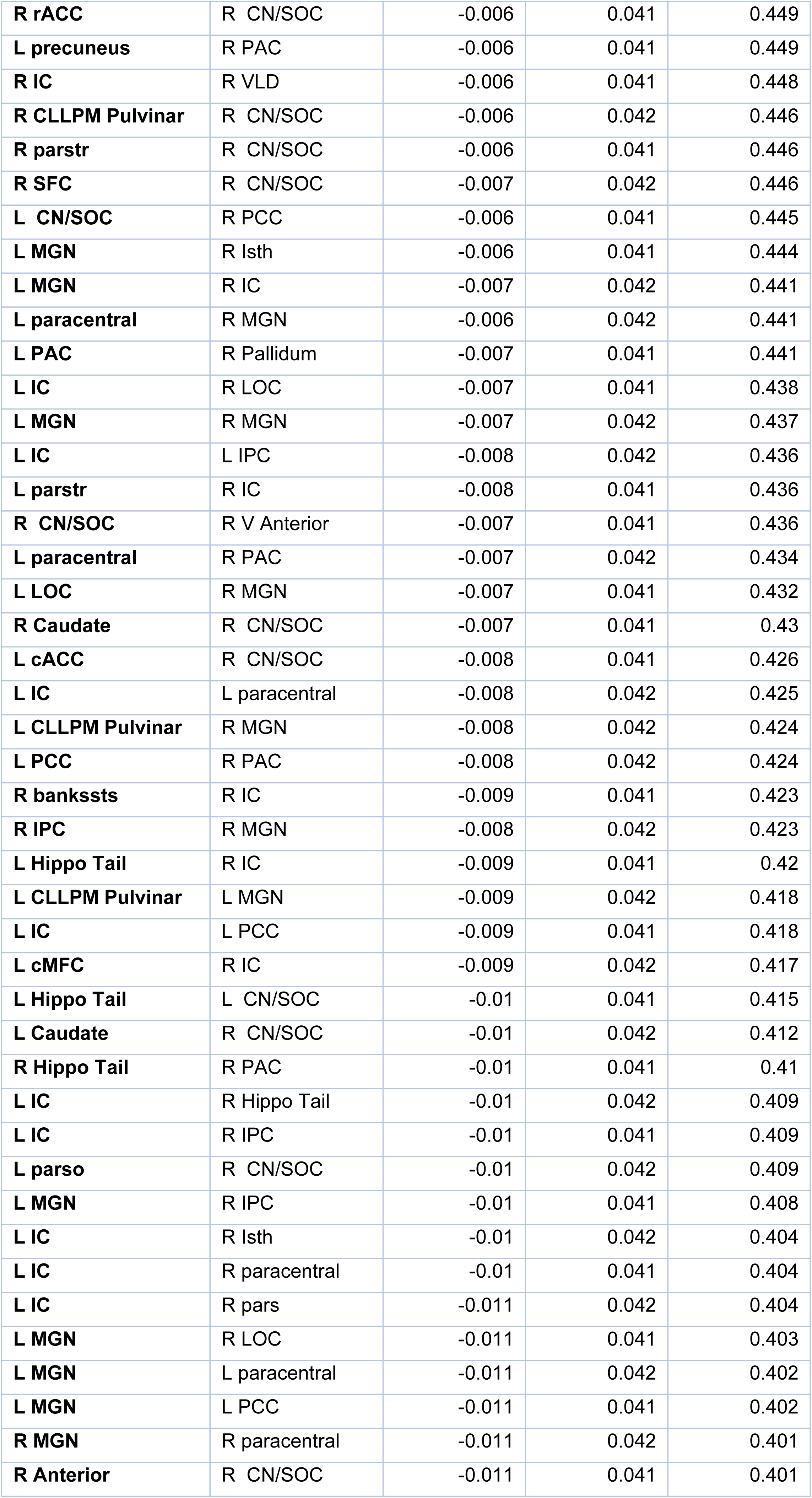
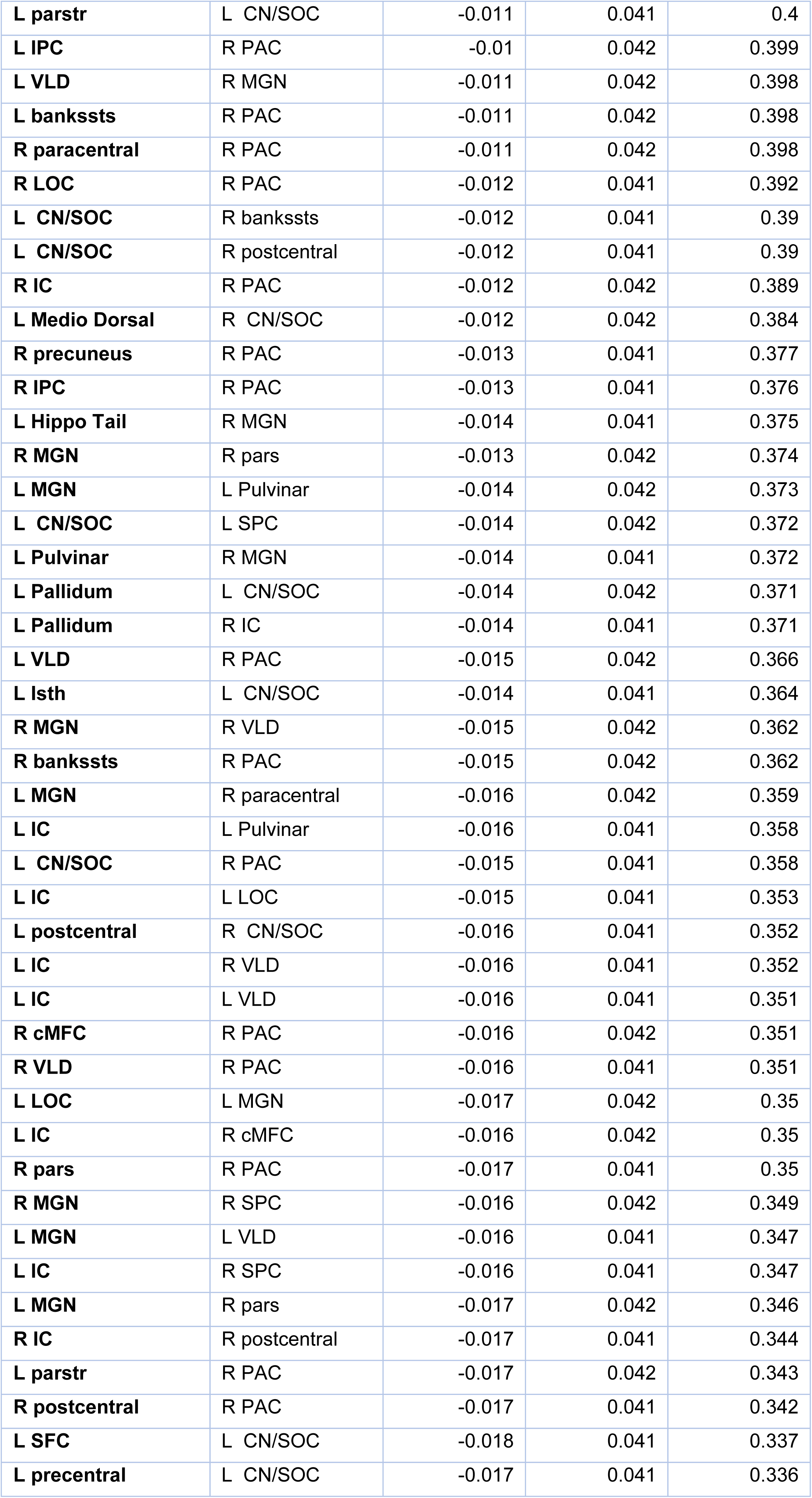
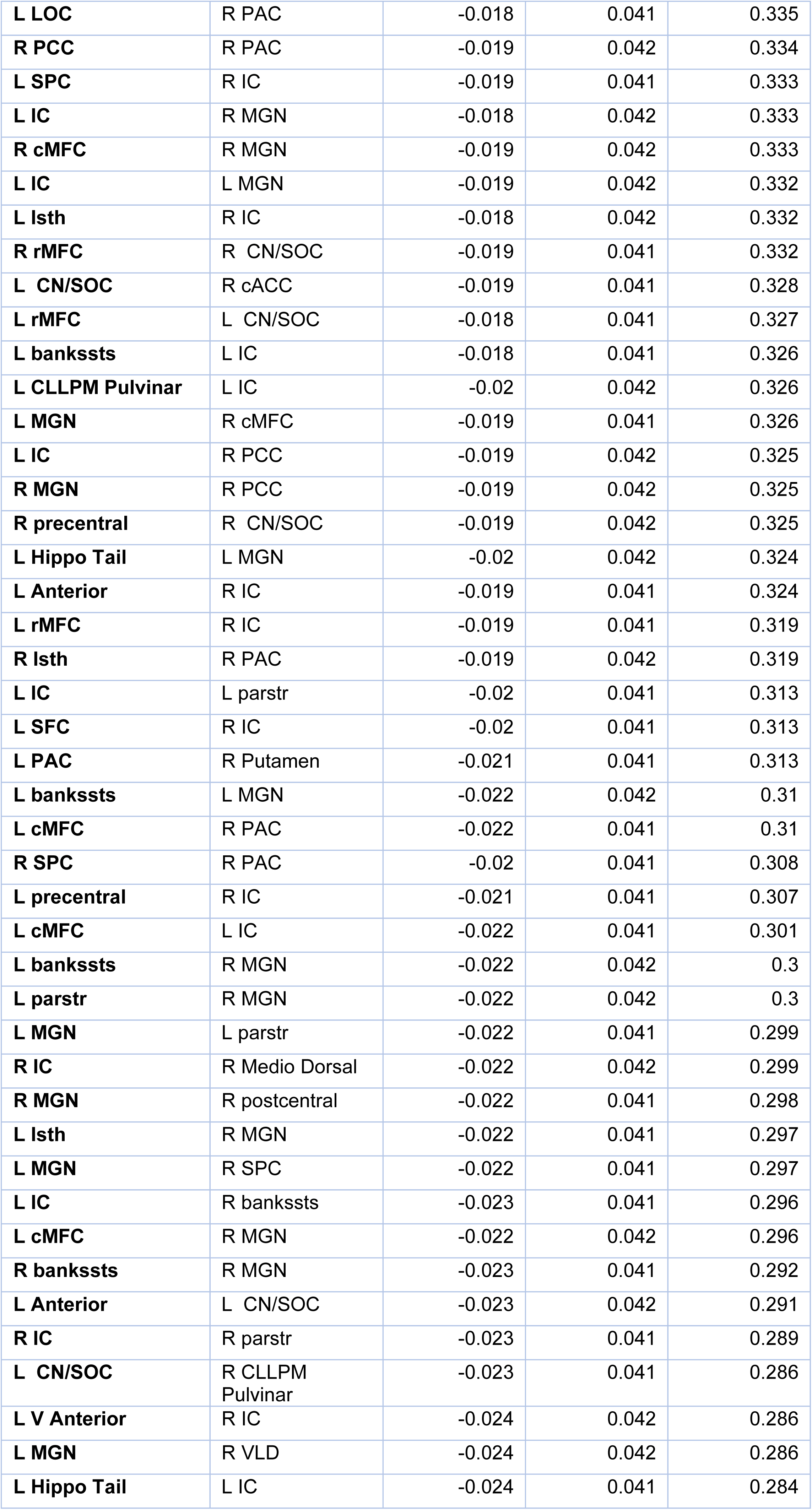
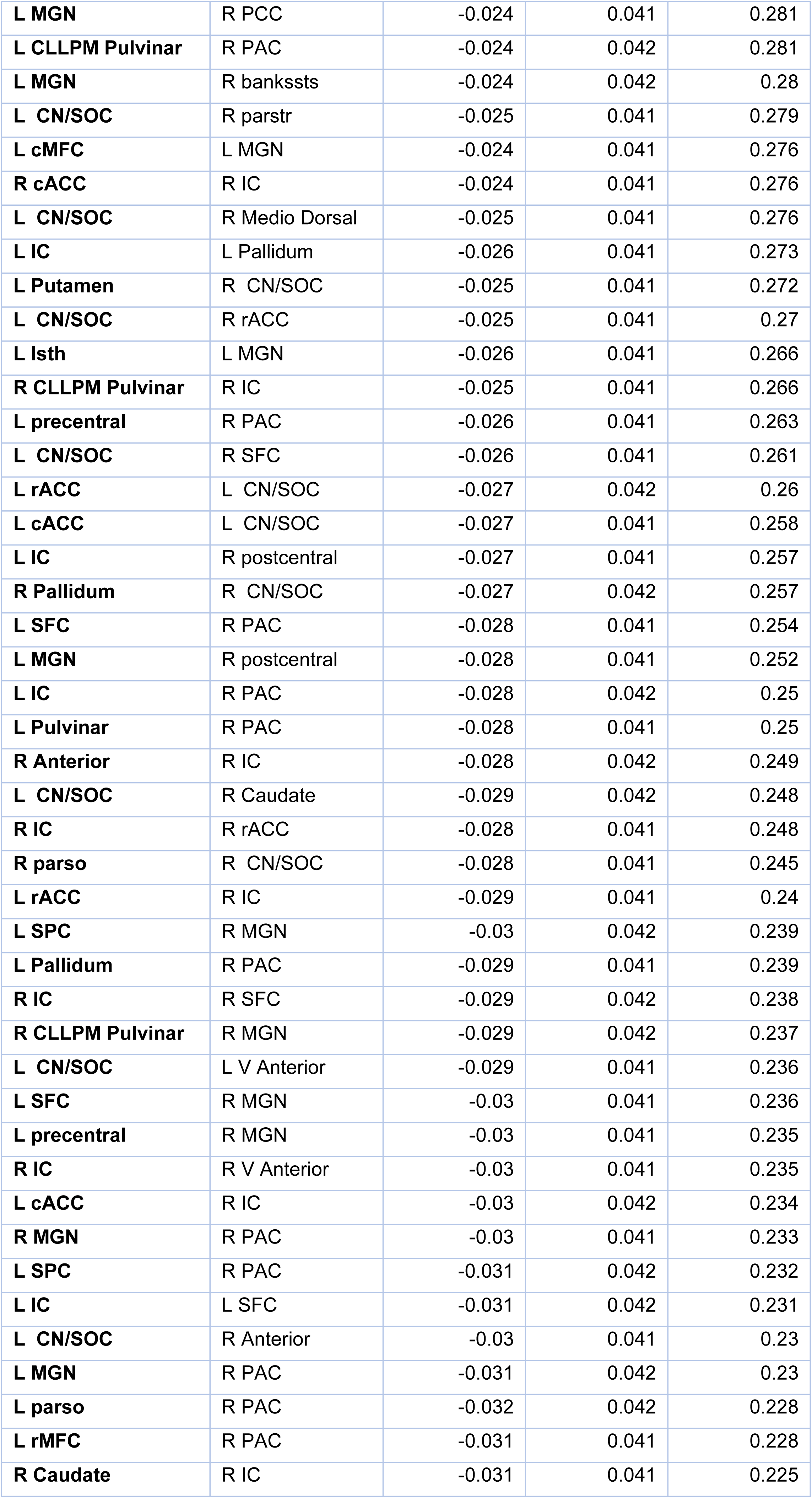
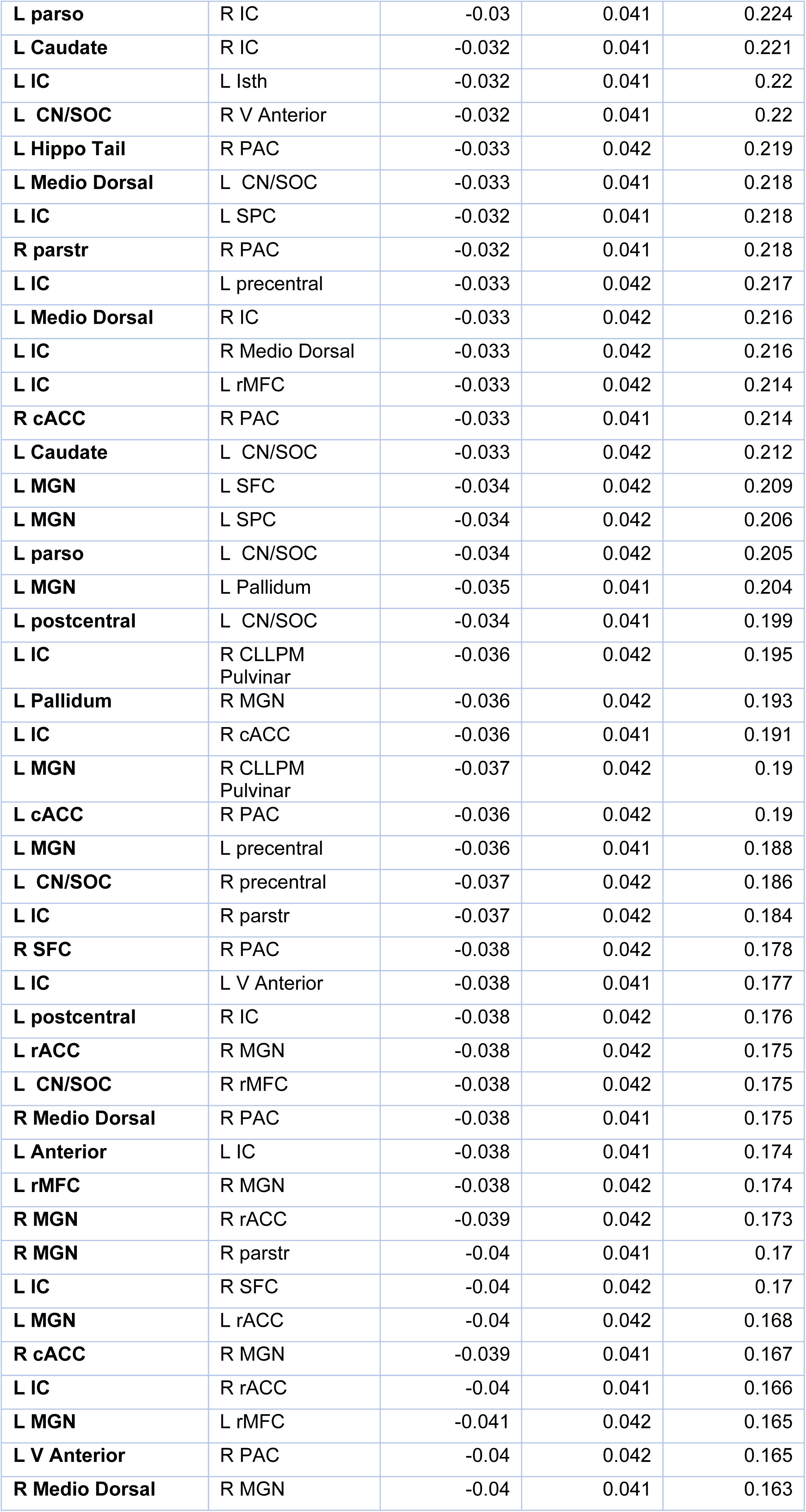
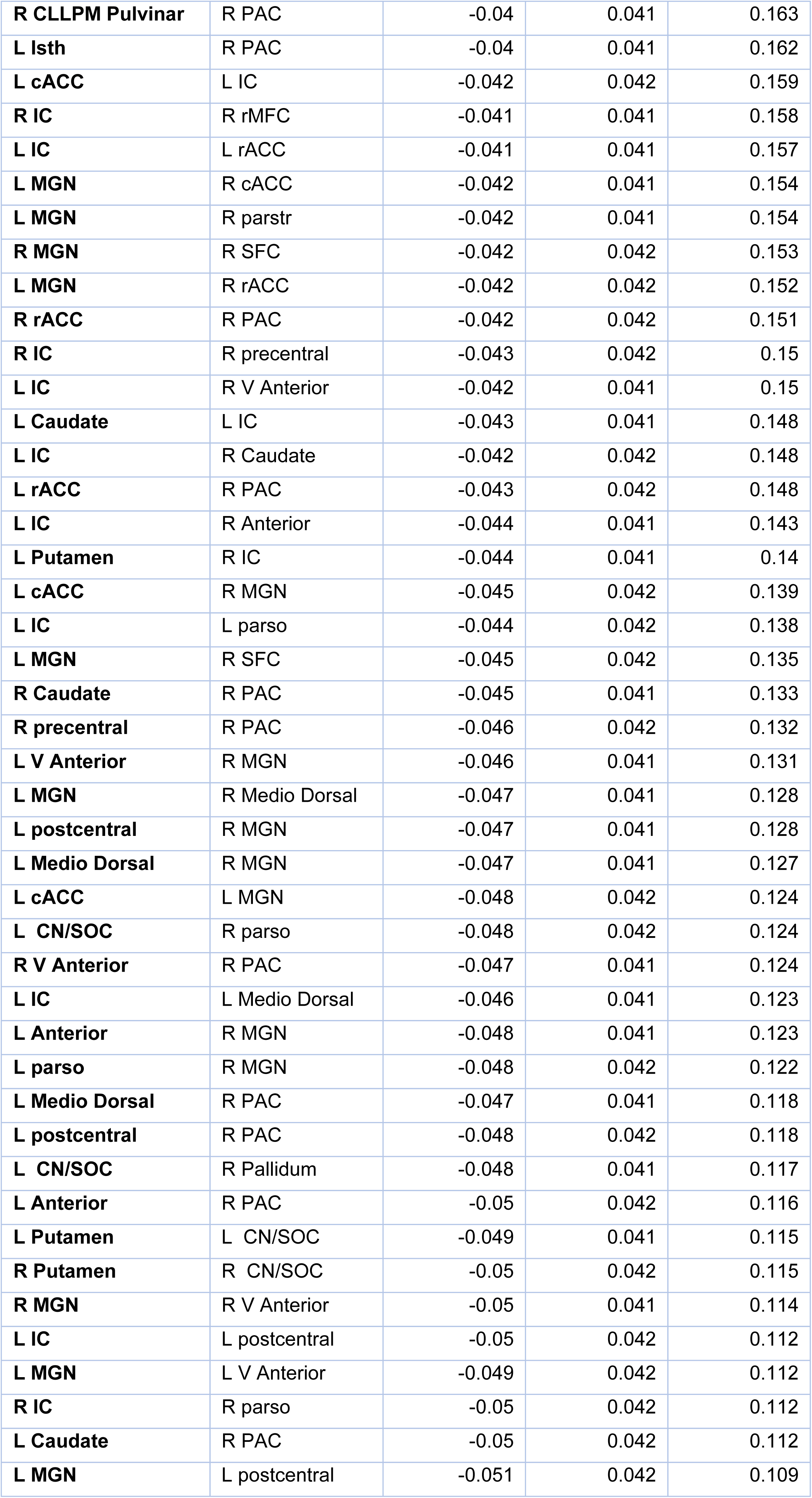
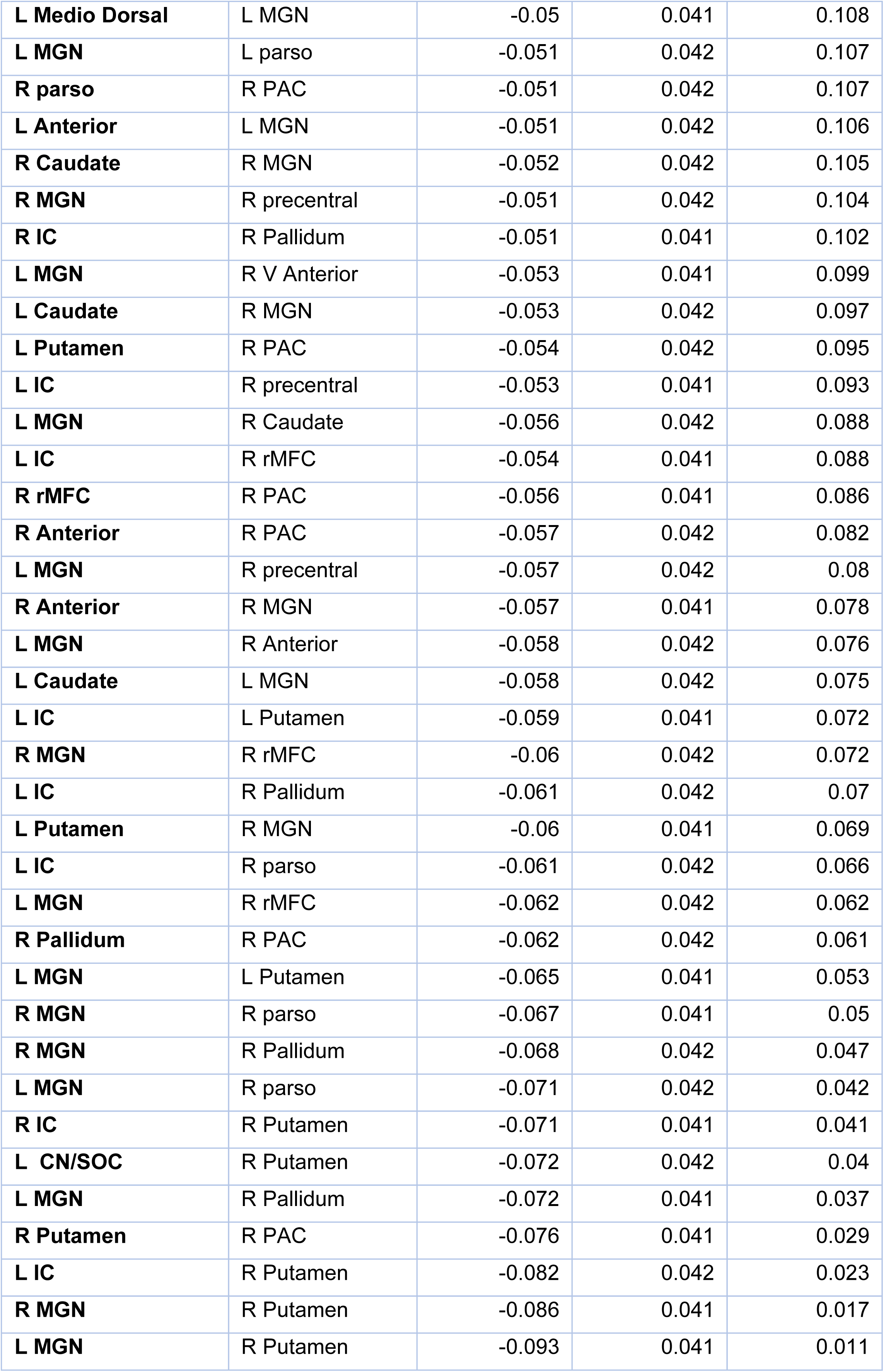
Effect estimates for the ROI pairs comparison between CHUU and CPHIV. P+ values are arranged in descending order.

### Appendix D: Summary statistics of functional network nodal measures, CHUU-CPHIV

**Table 5:**
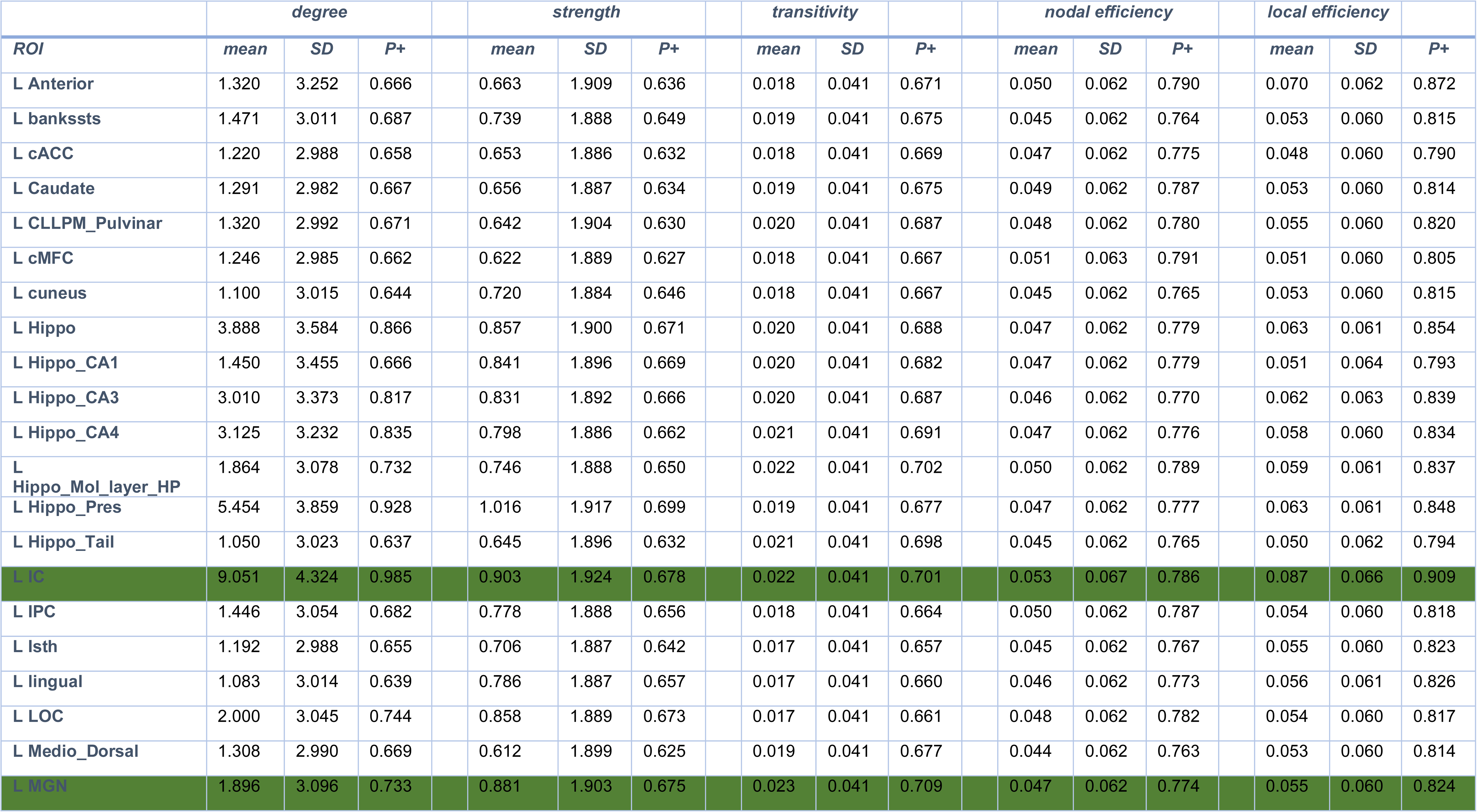
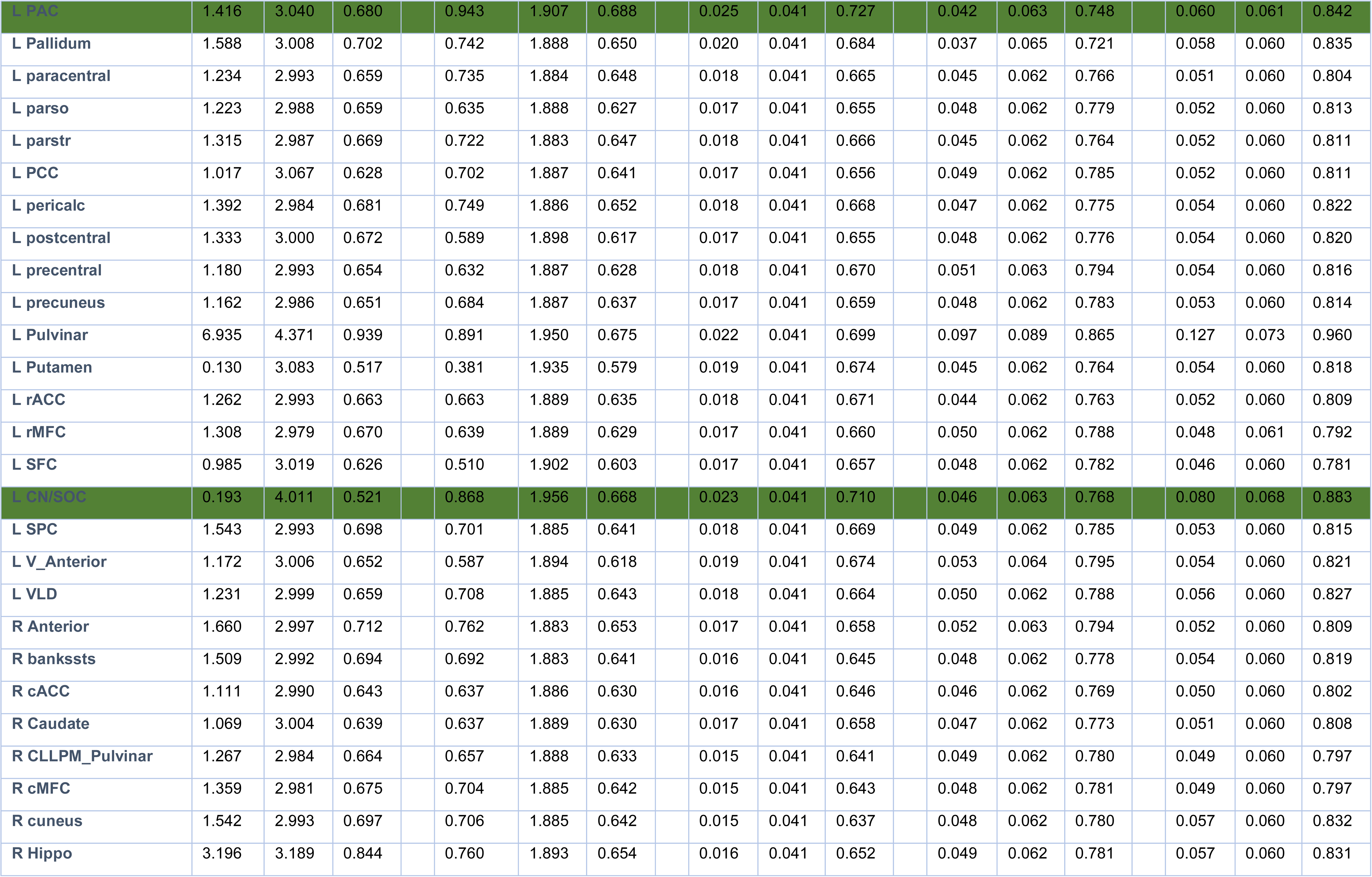
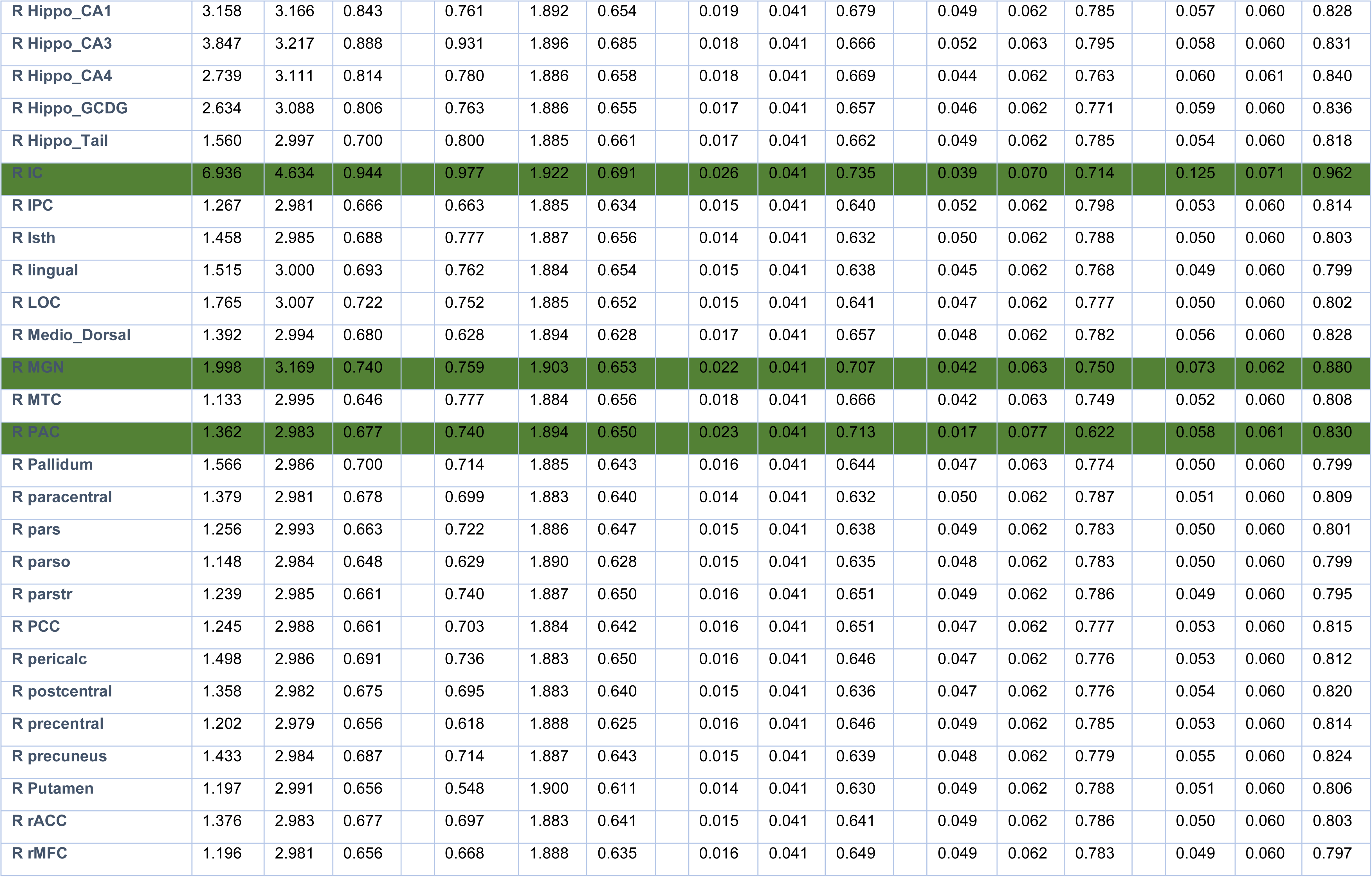
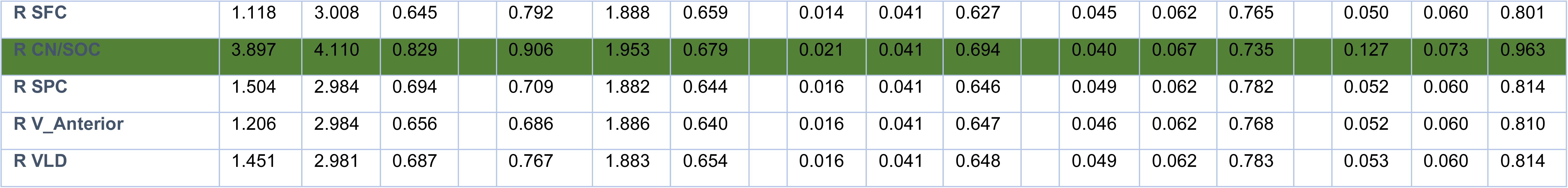
Summary statistics for CHUU vs CPHIV for the nodal graph measures degree, strength, transitivity, nodal and local efficiency. Auditory ROI rows are marked in green.

