## Appendix A: Automatically segmented regions of interest. for "Evidence of functional connectivity disruptions between auditory and non-auditory regions in adolescents living with HIV"

### Appendices

#### Appendix A: Automatically segmented regions of interest.

*Table 1: 126 automatically segmented ROIs. Asterisks denote the 54 regions excluded due to overlapping with manually traced regions in at least participant. R – right, L – left.*

| ROI | ROI abbreviation |
|---|---|
| R-lateralorbitofrontal* | R-lOFC |
| **R-parsorbitalis** | **R-pars** |
| R-frontalpole* | R-FP |
| R-medialorbitofrontal* | R-mOFC |
| **R-parstriangularis** | **R-parstr** |
| **R-parsopercularis** | **R-parso** |
| **R-rostralmiddlefrontal** | **R-rMFC** |
| **R-superiorfrontal** | **R-SFC** |
| **R-caudalmiddlefrontal** | **R-cMFC** |
| **R-precentral** | **R-precentral** |
| **R-paracentral** | **R-paracentral** |
| **R-rostralanteriorcingulate** | **R-rACC** |
| **R-caudalanteriorcingulate** | **R-cACC** |
| **R-posteriorcingulate** | **R-PCC** |
| **R-isthmuscingulate** | **R-Isth** |
| **R-postcentral** | **R-postcentral** |
| R-supramarginal* | R-supramarginal |
| **R-superiorparietal** | **R-SPC** |
| **R-inferiorparietal** | **R-IPC** |
| **R-precuneus** | **R-precuneus** |
| **R-cuneus** | **R-cuneus** |
| **R-pericalcarine** | **R-pericalc** |
| **R-lateraloccipital** | **R-LOC** |
| **R-lingual** | **R-lingual** |
| R-fusiform* | R-fusiform |
| R-parahippocampal* | R-paraHippo |
| R-entorhinal* | R-entorhinal |
| R-temporalpole* | R-TP |
| R-inferiortemporal* | R-ITC |
| **R-middletemporal** | **R-MTC** |
| **R-bankssts** | **R-bankssts** |
| R-superiortemporal* | R-STC |
| R-transversetemporal* | R-TTC |
| R-insula* | R-insula |

| | |
|---|---|
| R-Pulvinar* | R-Pulvinar |
| **R-Anterior** | **R-Anterior** |
| **R-Medio_Dorsal** | **R-Medio_Dorsal** |
| **R-Ventral_Latero_Dorsal** | **R-VLD** |
| **R-Central_Lateral-Lateral_Posterior-Medial_Pulvinar** | **R-CLLPM_Pulvinar** |
| **R-Ventral_Anterior** | **R-Ventral_Anterior** |
| R-Ventral_Latero_Ventral* | R-VLV |
| **R-Caudate** | **R-Caudate** |
| **R-Putamen** | **R-Putamen** |
| **R-Pallidum** | **R-Pallidum** |
| R-Accumbens_area* | R-Accumbens |
| R-Amygdala* | R-Amygdala |
| **R-Hippocampus** | **R-Hippo** |
| R-Hippocampus_Parasubiculum* | R-Hippo_Para |
| R-Hippocampus_Presubiculum* | R-Hippo_Pres |
| R-Hippocampus_Subiculum* | R-Hippo_Sub |
| **R-Hippocampus_CA1** | **R-Hippo_CA1** |
| **R-Hippocampus_CA3** | **R-Hippo_CA3** |
| **R-Hippocampus_CA4** | **R-Hippo_CA4** |
| **R-Hippocampus_GCDG** | **R-Hippo_GCDG** |
| R-Hippocampus_HATA* | R-Hippo_HATA |
| R-Hippocampus_Fimbria* | R-Hippo_Fimb |
| R-Hippocampus_Molecular_layer_HP* | R-Hippo_Mol_layer_HP |
| R-Hippocampus_Hippocampal_fissure* | R-Hippo_fissure |
| **R-Hippocampus_Tail** | **R-Hippo_Tail** |
| R-VentralDC* | R-VDC |
| R-Hypothalamus* | R-Hypothalamus |
| L-lateralorbitofrontal* | L-lOFC |
| L-parsorbitalis* | L-pars |
| L-frontalpole* | L-FP |
| L-medialorbitofrontal* | L-mOFC |
| **L-parstriangularis** | **L-parstr** |
| **L-parsopercularis** | **L-parso** |
| **L-rostralmiddlefrontal** | **L-rMFC** |
| **L-superiorfrontal** | **L-SFC** |
| **L-caudalmiddlefrontal** | **L-cMFC** |
| **L-precentral** | **L-precentral** |
| **L-paracentral** | **L-paracentral** |
| **L-rostralanteriorcingulate** | **L-rACC** |
| **L-caudalanteriorcingulate** | **L-cACC** |
| **L-posteriorcingulate** | **L-PCC** |
| **L-isthmuscingulate** | **L-Isth** |

| | |
|---|---|
| **L-postcentral** | **L-postcentral** |
| L-supramarginal* | L-supramarginal |
| **L-superiorparietal** | **L-SPC** |
| **L-inferiorparietal** | **L-IPC** |
| **L-precuneus** | **L-precuneus** |
| **L-cuneus** | **L-cuneus** |
| **L-pericalcarine** | **L-pericalc** |
| **L-lateraloccipital** | **L-LOC** |
| **L-lingual** | **L-lingual** |
| L-fusiform* | L-fusiform |
| L-parahippocampal* | L-paraHippo |
| L-entorhinal* | L-entorhinal |
| L-temporalpole* | L-TP |
| L-inferiortemporal* | L-ITC |
| L-middletemporal* | L-MTC |
| **L-bankssts** | **L-bankssts** |
| L-superiortemporal* | L-STC |
| L-transversetemporal* | L-TTC |
| L-insula* | L-insula |
| **L-Pulvinar** | **L-Pulvinar** |
| **L-Anterior** | **L-Anterior** |
| **L-Medio_Dorsal** | **L-Medio_Dorsal** |
| **L-Ventral_Latero_Dorsal** | **L-VLD** |
| **L-Central_Lateral-Lateral_Posterior-Medial_Pulvinar** | **L-CLLPM_Pulvinar** |
| **L-Ventral_Anterior** | **L-Ventral_Anterior** |
| L-Ventral_Latero_Ventral* | L-VLV |
| **L-Caudate** | **L-Caudate** |
| **L-Putamen** | **L-Putamen** |
| **L-Pallidum** | **L-Pallidum** |
| L-Accumbens_area* | L-Accumbens |
| L-Amygdala* | L-Amygdala |
| **L-Hippocampus** | **L-Hippo** |
| L-Hippocampus_Parasubiculum* | L-Hippo_Para |
| **L-Hippocampus_Presubiculum** | **L-Hippo_Pres** |
| L-Hippocampus_Subiculum* | L-Hippo_Sub |
| **L-Hippocampus_CA1** | **L-Hippo_CA1** |
| **L-Hippocampus_CA3** | **L-Hippo_CA3** |
| **L-Hippocampus_CA4** | **L-Hippo_CA4** |
| L-Hippocampus_GCDG* | L-Hippo_GCDG |
| L-Hippocampus_HATA* | L-Hippo_HATA |
| L-Hippocampus_Fimbria* | L-Hippo_Fimb |
| **L-Hippocampus_Molecular_layer_HP** | **L-Hippo_Mol_layer_HP** |

| | |
|---|---|
| L-Hippocampus_Hippocampal_fissure* | L-Hippo_fissure |
| **L-Hippocampus_Tail** | **L-Hippo_Tail** |
| L-VentralDC* | L-VDC |
| L-Hypothalamus* | **L-Hypothalamus** |
| Brain_Stem-Midbrain* | BS-mid |
| Brain_Stem-Pons* | BS-pons |
| Brain_Stem-Medulla* | BS-medulla |
| Brain_Stem-SCP* | BS-SCP |

#### Appendix B: Summary information of model results for FC of CHUU vs CPHIV

| **Group-level effects** | | | | | | | |
|---|---|---|---|---|---|---|---|
| | **Estimate** | **Est.Error** | **l-95% CI** | **u-95% CI** | **Rhat** | **Bulk_ESS** | **Tail_ESS** |
| **ROIs (Number of levels: 80):** | | | | | | | |
| **sd(Intercept)** | 0.05 | 0.00 | 0.04 | 0.06 | 1.01 | 779 | 1780 |
| **sd(group1)** | 0.01 | 0 | 0.01 | 0.02 | 1 | 889 | 1317 |
| **sd(sex1)** | 0.01 | 0 | 0.01 | 0.01 | 1 | 1109 | 2256 |
| **cor(Intercept,group1)** | -0.09 | 0.12 | -0.32 | 0.16 | 1 | 413 | 1054 |
| **cor(Intercept,sex1)** | 0.13 | 0.12 | -0.1 | 0.37 | 1 | 860 | 1554 |
| **cor(group1,sex1)** | 0.56 | 0.08 | 0.39 | 0.71 | 1 | 1062 | 2111 |
| **ROI pairs (Number of levels: 3160):** | | | | | | | |
| **sd(Intercept)** | 0.16 | 0 | 0.16 | 0.17 | 1.01 | 646 | 1271 |
| **sd(group1)** | 0.01 | 0 | 0.01 | 0.01 | 1 | 688 | 1365 |
| **sd(sex1)** | 0 | 0 | 0 | 0.01 | 1 | 2828 | 3575 |
| **cor(Intercept,group1)** | 0.79 | 0.08 | 0.64 | 0.96 | 1 | 568 | 811 |
| **cor(Intercept,sex1)** | 0.85 | 0.09 | 0.64 | 0.98 | 1 | 2015 | 3198 |
| **cor(group1,sex1)** | 0.88 | 0.09 | 0.65 | 0.99 | 1 | 2012 | 3286 |
| **Subjects (Number of levels: 78):** | | | | | | | |
| **sd(Intercept)** | 0.1 | 0.01 | 0.09 | 0.12 | 1.01 | 460 | 993 |
| **Population-Level Effects** | | | | | | | |
| **Intercept** | 0.17 | 0.02 | 0.14 | 0.2 | 1.01 | 407 | 870 |
| **group1** | 0 | 0.01 | -0.03 | 0.02 | 1 | 412 | 781 |

| | | | | | | | |
|---|---|---|---|---|---|---|---|
| **sex1** | 0.01 | 0.01 | -0.02 | 0.03 | 1.01 | 345 | 712 |
| **Family Specific Parameters:** | | | | | | | |
| **sigma** | 0.21 | 0 | 0.21 | 0.21 | 1 | 5829 | 5049 |
| **nu** | 16.08 | 0.47 | 15.19 | 17.07 | 1 | 6658 | 4757 |

*Samples were drawn using sample (hmc). For each parameter, Bulk_ESS and Tail_ESS are effective sample size measures, and Rhat is the potential scale reduction factor on split chains (at convergence, Rhat = 1).*

#### Appendix C: Region and region pair effect estimates

*Table 2: ROI effect estimates and their uncertainties for the comparison CHUU-CPHIV. The first 3 columns are difference between group mean FCs (Fisher's Z-values), standard deviations, and the posterior probability of the effect being positive respectively. The preceding columns represent uncertainty intervals. The ROIs are organized in P+ descending order. Auditory ROI rows are marked in green and ROIs showing strong evidence of altered FC (P+ >= 0.95 | P+<= 0.05) are in bold.*

| ROI | mean | SD | P+ | 2.50% | 5% | 50% | 95% | 97.50% |
|---|---|---|---|---|---|---|---|---|
| **L-Hippo_CA1** | **0.061** | **0.020** | **0.997** | **0.021** | **0.028** | **0.062** | **0.094** | **0.101** |
| **L-Hippo** | **0.053** | **0.020** | **0.993** | **0.011** | **0.018** | **0.053** | **0.085** | **0.091** |
| **L-Hippo_Pres** | **0.052** | **0.020** | **0.993** | **0.011** | **0.018** | **0.053** | **0.085** | **0.091** |
| **L-Hippo_Mol_layer_HP** | **0.049** | **0.020** | **0.992** | **0.009** | **0.016** | **0.050** | **0.082** | **0.088** |
| **R-lingual** | **0.051** | **0.020** | **0.992** | **0.010** | **0.017** | **0.051** | **0.083** | **0.089** |
| **L-Hippo_CA3** | **0.052** | **0.020** | **0.991** | **0.011** | **0.018** | **0.052** | **0.084** | **0.091** |
| **L-Hippo_CA4** | **0.049** | **0.020** | **0.990** | **0.009** | **0.015** | **0.049** | **0.082** | **0.089** |
| **R-Hippo_CA3** | **0.044** | **0.020** | **0.983** | **0.003** | **0.010** | **0.045** | **0.078** | **0.084** |
| **R-Hippo_CA1** | **0.043** | **0.020** | **0.979** | **0.001** | **0.008** | **0.043** | **0.076** | **0.081** |
| **L-lingual** | **0.042** | **0.020** | **0.977** | **0.001** | **0.008** | **0.042** | **0.074** | **0.081** |
| **L-PAC** | **0.039** | **0.020** | **0.973** | **-0.001** | **0.005** | **0.039** | **0.071** | **0.078** |
| **R-Hippo** | **0.039** | **0.020** | **0.972** | **-0.001** | **0.005** | **0.039** | **0.072** | **0.078** |
| **R-Hippo_CA4** | 0.028 | 0.020 | 0.917 | -0.013 | -0.006 | 0.028 | 0.061 | 0.067 |
| **R-Hippo_GCDG** | 0.026 | 0.020 | 0.901 | -0.014 | -0.007 | 0.026 | 0.059 | 0.066 |
| **R-SOC-CN** | 0.024 | 0.020 | 0.878 | -0.017 | -0.011 | 0.024 | 0.057 | 0.064 |
| **R-pericalc** | 0.023 | 0.020 | 0.872 | -0.018 | -0.011 | 0.023 | 0.056 | 0.062 |
| **L-pericalc** | 0.023 | 0.020 | 0.870 | -0.018 | -0.011 | 0.023 | 0.055 | 0.061 |
| **L-precuneus** | 0.021 | 0.020 | 0.847 | -0.019 | -0.013 | 0.021 | 0.053 | 0.059 |
| **R-MTC** | 0.018 | 0.020 | 0.817 | -0.022 | -0.016 | 0.018 | 0.050 | 0.057 |
| **L-cuneus** | 0.016 | 0.020 | 0.786 | -0.025 | -0.018 | 0.016 | 0.049 | 0.055 |
| **R-cuneus** | 0.015 | 0.020 | 0.770 | -0.026 | -0.020 | 0.015 | 0.047 | 0.054 |
| **L-IPC** | 0.014 | 0.021 | 0.756 | -0.029 | -0.020 | 0.014 | 0.047 | 0.053 |
| **R-precuneus** | 0.013 | 0.020 | 0.750 | -0.029 | -0.021 | 0.014 | 0.046 | 0.052 |
| **L-PCC** | 0.011 | 0.020 | 0.708 | -0.031 | -0.024 | 0.011 | 0.043 | 0.050 |
| **R-IPC** | 0.009 | 0.020 | 0.686 | -0.031 | -0.024 | 0.010 | 0.042 | 0.049 |
| **R-Isth** | 0.008 | 0.020 | 0.674 | -0.033 | -0.026 | 0.009 | 0.041 | 0.048 |

| | | | | | | | | |
|---|---|---|---|---|---|---|---|---|
| **L-paracentral** | 0.007 | 0.020 | 0.633 | -0.034 | -0.027 | 0.007 | 0.040 | 0.046 |
| **R-LOC** | 0.005 | 0.020 | 0.612 | -0.034 | -0.028 | 0.006 | 0.038 | 0.045 |
| **R-Hippo_Tail** | 0.006 | 0.020 | 0.610 | -0.035 | -0.029 | 0.006 | 0.038 | 0.043 |
| **L-SOC-CN** | 0.003 | 0.020 | 0.567 | -0.037 | -0.030 | 0.004 | 0.036 | 0.041 |
| **R-cMFC** | 0.003 | 0.020 | 0.563 | -0.039 | -0.032 | 0.003 | 0.035 | 0.042 |
| **R-pars** | 0.002 | 0.020 | 0.559 | -0.040 | -0.032 | 0.003 | 0.035 | 0.042 |
| **R-IC** | 0.002 | 0.020 | 0.536 | -0.038 | -0.031 | 0.002 | 0.035 | 0.041 |
| **R-paracentral** | 0.000 | 0.020 | 0.509 | -0.040 | -0.034 | 0.000 | 0.033 | 0.040 |
| **L-VLD** | -0.001 | 0.020 | 0.476 | -0.041 | -0.035 | -0.001 | 0.032 | 0.037 |
| **L-LOC** | -0.002 | 0.020 | 0.472 | -0.042 | -0.036 | -0.002 | 0.031 | 0.037 |
| **L-cMFC** | -0.002 | 0.020 | 0.468 | -0.042 | -0.036 | -0.002 | 0.031 | 0.038 |
| **R-PCC** | -0.003 | 0.020 | 0.446 | -0.044 | -0.038 | -0.003 | 0.029 | 0.035 |
| **R-VLD** | -0.003 | 0.020 | 0.442 | -0.044 | -0.037 | -0.003 | 0.029 | 0.035 |
| **R-SPC** | -0.005 | 0.020 | 0.403 | -0.046 | -0.040 | -0.005 | 0.028 | 0.034 |
| **L-bankssts** | -0.006 | 0.020 | 0.388 | -0.046 | -0.040 | -0.006 | 0.027 | 0.033 |
| **L-parstr** | -0.006 | 0.020 | 0.388 | -0.047 | -0.039 | -0.005 | 0.026 | 0.032 |
| **L-CLLPM_Pulvinar** | -0.007 | 0.020 | 0.374 | -0.048 | -0.040 | -0.006 | 0.026 | 0.032 |
| **R-bankssts** | -0.010 | 0.020 | 0.319 | -0.051 | -0.044 | -0.009 | 0.023 | 0.030 |
| **L-Pulvinar** | -0.010 | 0.020 | 0.315 | -0.050 | -0.044 | -0.010 | 0.023 | 0.028 |
| **L-SFC** | -0.010 | 0.020 | 0.306 | -0.051 | -0.045 | -0.010 | 0.022 | 0.027 |
| **L-IC** | -0.011 | 0.020 | 0.298 | -0.051 | -0.044 | -0.011 | 0.021 | 0.028 |
| **R-MGN** | -0.011 | 0.020 | 0.286 | -0.052 | -0.046 | -0.011 | 0.022 | 0.028 |
| **L-Hippo_Tail** | -0.012 | 0.020 | 0.271 | -0.054 | -0.047 | -0.012 | 0.020 | 0.026 |
| **L-Isth** | -0.012 | 0.020 | 0.269 | -0.053 | -0.047 | -0.012 | 0.020 | 0.026 |
| **R-postcentral** | -0.013 | 0.020 | 0.265 | -0.053 | -0.046 | -0.012 | 0.020 | 0.025 |
| **L-MGN** | -0.014 | 0.020 | 0.248 | -0.056 | -0.049 | -0.014 | 0.018 | 0.025 |
| **L-SPC** | -0.016 | 0.021 | 0.230 | -0.057 | -0.050 | -0.015 | 0.017 | 0.024 |
| **L-rMFC** | -0.015 | 0.020 | 0.225 | -0.056 | -0.049 | -0.015 | 0.017 | 0.024 |
| **L-precentral** | -0.016 | 0.020 | 0.218 | -0.056 | -0.050 | -0.016 | 0.017 | 0.023 |
| **R-PAC** | -0.016 | 0.020 | 0.210 | -0.057 | -0.050 | -0.016 | 0.016 | 0.023 |
| **L-Pallidum** | -0.017 | 0.020 | 0.199 | -0.058 | -0.050 | -0.017 | 0.016 | 0.022 |
| **R-cACC** | -0.021 | 0.020 | 0.148 | -0.061 | -0.055 | -0.021 | 0.012 | 0.018 |
| **R-SFC** | -0.021 | 0.021 | 0.146 | -0.063 | -0.056 | -0.020 | 0.012 | 0.018 |
| **R-parstr** | -0.022 | 0.020 | 0.138 | -0.062 | -0.055 | -0.022 | 0.010 | 0.017 |
| **R-CLLPM_Pulvinar** | -0.025 | 0.020 | 0.108 | -0.065 | -0.059 | -0.025 | 0.007 | 0.014 |
| **L-rACC** | -0.026 | 0.021 | 0.101 | -0.067 | -0.060 | -0.026 | 0.007 | 0.013 |
| **R-Medio_Dorsal** | -0.025 | 0.020 | 0.097 | -0.067 | -0.059 | -0.025 | 0.007 | 0.014 |
| **L-V_Anterior** | -0.026 | 0.020 | 0.092 | -0.067 | -0.061 | -0.025 | 0.006 | 0.012 |
| **R-rACC** | -0.027 | 0.020 | 0.090 | -0.067 | -0.060 | -0.026 | 0.006 | 0.013 |
| **L-cACC** | -0.027 | 0.021 | 0.083 | -0.069 | -0.062 | -0.027 | 0.005 | 0.012 |
| **L-parso** | -0.029 | 0.020 | 0.070 | -0.071 | -0.064 | -0.029 | 0.003 | 0.009 |
| **R-Caudate** | -0.030 | 0.020 | 0.069 | -0.071 | -0.064 | -0.030 | 0.004 | 0.011 |

| | | | | | | | | |
|---|---|---|---|---|---|---|---|---|
| **R-V_Anterior** | -0.030 | 0.020 | 0.061 | -0.071 | -0.064 | -0.030 | 0.002 | 0.009 |
| **L-Anterior** | **-0.032** | **0.020** | **0.056** | **-0.073** | **-0.066** | **-0.031** | **0.001** | **0.007** |
| **L-Medio_Dorsal** | **-0.034** | **0.020** | **0.044** | **-0.076** | **-0.069** | **-0.034** | **-0.001** | **0.005** |
| **L-Caudate** | **-0.034** | **0.020** | **0.043** | **-0.076** | **-0.069** | **-0.034** | **-0.002** | **0.005** |
| **R-Anterior** | **-0.037** | **0.020** | **0.032** | **-0.078** | **-0.071** | **-0.037** | **-0.005** | **0.002** |
| **L-postcentral** | **-0.037** | **0.020** | **0.031** | **-0.079** | **-0.071** | **-0.037** | **-0.005** | **0.002** |
| **R-rMFC** | **-0.038** | **0.020** | **0.028** | **-0.079** | **-0.072** | **-0.038** | **-0.006** | **0.001** |
| **L-Putamen** | **-0.044** | **0.020** | **0.015** | **-0.085** | **-0.078** | **-0.043** | **-0.011** | **-0.005** |
| **R-parso** | **-0.049** | **0.020** | **0.007** | **-0.090** | **-0.083** | **-0.048** | **-0.016** | **-0.009** |
| **R-Pallidum** | **-0.053** | **0.020** | **0.004** | **-0.094** | **-0.087** | **-0.052** | **-0.020** | **-0.014** |
| **R-Putamen** | **-0.070** | **0.020** | **0.001** | **-0.111** | **-0.104** | **-0.070** | **-0.038** | **-0.031** |

*Table 3: Effect estimates for the ROI pairs comparison between CHUU and CPHIV. P+ values are arranged in descending order.*

| ROI1 | ROI2 | mean | SD | Pplus |
|---|---|---|---|---|
| **L Hippo CA1** | L PAC | 0.095 | 0.041 | 0.986 |
| **L PAC** | R lingual | 0.093 | 0.041 | 0.986 |
| **L Hippo CA1** | R CN/SOC | 0.086 | 0.042 | 0.978 |
| **L lingual** | L PAC | 0.082 | 0.042 | 0.974 |
| **L Hippo CA3** | R CN/SOC | 0.081 | 0.042 | 0.973 |
| **L Hippo** | L PAC | 0.08 | 0.042 | 0.971 |
| **L Hippo CA3** | L PAC | 0.08 | 0.042 | 0.97 |
| **L Hippo CA4** | L PAC | 0.079 | 0.041 | 0.97 |
| **L Hippo Pres** | L PAC | 0.082 | 0.042 | 0.97 |
| **L Hippo Pres** | R CN/SOC | 0.08 | 0.042 | 0.968 |
| **L Hippo** | R CN/SOC | 0.079 | 0.042 | 0.967 |
| **L Hippo Mol layer HP** | L PAC | 0.077 | 0.041 | 0.965 |
| **L PAC** | R Hippo CA1 | 0.076 | 0.041 | 0.964 |
| **L Hippo CA4** | R CN/SOC | 0.076 | 0.042 | 0.961 |
| **L PAC** | R Hippo CA3 | 0.074 | 0.041 | 0.959 |
| **L Hippo Mol layer HP** | R CN/SOC | 0.075 | 0.041 | 0.959 |
| **R Hippo CA3** | R CN/SOC | 0.074 | 0.042 | 0.958 |
| **R lingual** | R CN/SOC | 0.073 | 0.041 | 0.954 |
| **R Hippo CA1** | R CN/SOC | 0.07 | 0.041 | 0.952 |
| **R Hippo** | R CN/SOC | 0.068 | 0.042 | 0.946 |
| **L PAC** | R Hippo | 0.068 | 0.042 | 0.944 |
| **L Hippo CA1** | R IC | 0.066 | 0.042 | 0.942 |
| **L CN/SOC** | R CN/SOC | 0.065 | 0.043 | 0.939 |
| **L PAC** | R pericalc | 0.064 | 0.042 | 0.938 |
| **L lingual** | R CN/SOC | 0.063 | 0.041 | 0.933 |
| **L Hippo CA1** | L CN/SOC | 0.061 | 0.042 | 0.929 |
| **L PAC** | L pericalc | 0.062 | 0.042 | 0.927 |
| **L Hippo Pres** | R IC | 0.061 | 0.042 | 0.922 |
| **L PAC** | R CN/SOC | 0.058 | 0.041 | 0.921 |
| **L PAC** | R Hippo CA4 | 0.058 | 0.041 | 0.918 |
| **L Hippo CA3** | R IC | 0.058 | 0.042 | 0.916 |
| **L PAC** | R MTC | 0.058 | 0.042 | 0.914 |
| **L Hippo Pres** | L MGN | 0.057 | 0.042 | 0.913 |
| **L PAC** | R Hippo GCDG | 0.056 | 0.041 | 0.913 |
| **L Hippo** | R IC | 0.058 | 0.042 | 0.913 |
| **L Hippo CA3** | L CN/SOC | 0.057 | 0.041 | 0.91 |
| **R Hippo CA4** | R CN/SOC | 0.056 | 0.041 | 0.91 |
| **L Hippo** | L CN/SOC | 0.055 | 0.041 | 0.905 |
| **L Hippo Pres** | L CN/SOC | 0.055 | 0.042 | 0.905 |
| **L Hippo CA4** | R IC | 0.055 | 0.042 | 0.904 |

| | | | | |
|---|---|---|---|---|
| **L PAC** | L PCC | 0.053 | 0.041 | 0.903 |
| **L CN/SOC** | R Hippo CA3 | 0.055 | 0.041 | 0.903 |
| **R Hippo GCDG** | R CN/SOC | 0.053 | 0.042 | 0.901 |
| **L Hippo Mol layer HP** | R IC | 0.053 | 0.042 | 0.899 |
| **L Hippo CA1** | L IC | 0.053 | 0.042 | 0.898 |
| **L PAC** | L paracentral | 0.052 | 0.041 | 0.897 |
| **R Hippo CA3** | R IC | 0.053 | 0.042 | 0.896 |
| **L PAC** | R cuneus | 0.051 | 0.041 | 0.894 |
| **L Hippo CA1** | L MGN | 0.051 | 0.042 | 0.891 |
| **L Hippo CA1** | R MGN | 0.051 | 0.041 | 0.89 |
| **L Hippo Pres** | R MGN | 0.05 | 0.042 | 0.889 |
| **L Hippo CA4** | L CN/SOC | 0.05 | 0.042 | 0.887 |
| **L PAC** | R paracentral | 0.05 | 0.042 | 0.887 |
| **L cuneus** | L PAC | 0.05 | 0.042 | 0.886 |
| **L CN/SOC** | R Hippo CA1 | 0.05 | 0.041 | 0.886 |
| **L CN/SOC** | R lingual | 0.05 | 0.041 | 0.886 |
| **L Hippo** | L MGN | 0.049 | 0.042 | 0.884 |
| **L Hippo Mol layer HP** | L CN/SOC | 0.049 | 0.041 | 0.88 |
| **L Hippo** | R MGN | 0.048 | 0.042 | 0.879 |
| **L PAC** | L precuneus | 0.048 | 0.041 | 0.875 |
| **L PAC** | R PAC | 0.048 | 0.042 | 0.875 |
| **R Hippo CA1** | R IC | 0.048 | 0.041 | 0.874 |
| **R pericalc** | R CN/SOC | 0.048 | 0.042 | 0.874 |
| **R IC** | R lingual | 0.047 | 0.042 | 0.873 |
| **R Hippo** | R IC | 0.047 | 0.042 | 0.868 |
| **L CN/SOC** | R Hippo | 0.045 | 0.041 | 0.866 |
| **R lingual** | R MGN | 0.044 | 0.041 | 0.86 |
| **L Hippo Pres** | L IC | 0.045 | 0.042 | 0.859 |
| **L PAC** | R bankssts | 0.043 | 0.041 | 0.859 |
| **L PAC** | R precuneus | 0.044 | 0.041 | 0.858 |
| **L Hippo CA3** | L MGN | 0.045 | 0.041 | 0.853 |
| **L Hippo Mol layer HP** | R MGN | 0.042 | 0.041 | 0.853 |
| **L bankssts** | L PAC | 0.042 | 0.041 | 0.852 |
| **R Hippo CA3** | R MGN | 0.044 | 0.042 | 0.852 |
| **L Hippo CA3** | R MGN | 0.044 | 0.042 | 0.851 |
| **L Hippo** | L IC | 0.042 | 0.042 | 0.85 |
| **L lingual** | L CN/SOC | 0.044 | 0.042 | 0.85 |
| **L PAC** | R Hippo Tail | 0.043 | 0.042 | 0.85 |
| **L IPC** | L PAC | 0.042 | 0.041 | 0.849 |
| **L pericalc** | R CN/SOC | 0.043 | 0.042 | 0.849 |
| **L Hippo CA3** | L IC | 0.043 | 0.042 | 0.848 |
| **L precuneus** | R CN/SOC | 0.042 | 0.041 | 0.848 |
| **L PAC** | R IC | 0.043 | 0.042 | 0.847 |

| | | | | |
|---|---|---|---|---|
| **L PAC** | R pars | 0.042 | 0.042 | 0.847 |
| **L Hippo Mol layer HP** | L MGN | 0.042 | 0.041 | 0.845 |
| **L PAC** | R cMFC | 0.041 | 0.042 | 0.844 |
| **L PAC** | R PCC | 0.042 | 0.041 | 0.844 |
| **L Hippo CA1** | R PAC | 0.04 | 0.041 | 0.841 |
| **L Hippo CA4** | L MGN | 0.041 | 0.041 | 0.84 |
| **L Hippo CA4** | R MGN | 0.04 | 0.041 | 0.838 |
| **L PAC** | R LOC | 0.04 | 0.041 | 0.834 |
| **L PAC** | R postcentral | 0.039 | 0.041 | 0.834 |
| **L cuneus** | R CN/SOC | 0.038 | 0.041 | 0.831 |
| **L PAC** | L parstr | 0.038 | 0.041 | 0.828 |
| **R Hippo** | R MGN | 0.039 | 0.042 | 0.828 |
| **R cuneus** | R CN/SOC | 0.038 | 0.042 | 0.828 |
| **L Hippo CA4** | L IC | 0.039 | 0.042 | 0.827 |
| **L MGN** | R lingual | 0.038 | 0.042 | 0.826 |
| **L PAC** | R IPC | 0.038 | 0.041 | 0.824 |
| **L PAC** | R Isth | 0.038 | 0.041 | 0.823 |
| **L Hippo Mol layer HP** | L IC | 0.038 | 0.042 | 0.822 |
| **L PAC** | L CN/SOC | 0.037 | 0.041 | 0.822 |
| **L MGN** | R Hippo CA3 | 0.037 | 0.041 | 0.82 |
| **L IC** | R IC | 0.037 | 0.042 | 0.816 |
| **L PAC** | L VLD | 0.037 | 0.041 | 0.815 |
| **L lingual** | R MGN | 0.037 | 0.041 | 0.815 |
| **L MGN** | R Hippo CA1 | 0.037 | 0.041 | 0.813 |
| **L PAC** | R VLD | 0.036 | 0.041 | 0.813 |
| **L CN/SOC** | R Hippo CA4 | 0.037 | 0.041 | 0.812 |
| **L IC** | R Hippo CA3 | 0.036 | 0.041 | 0.81 |
| **L IPC** | R CN/SOC | 0.035 | 0.041 | 0.808 |
| **L lingual** | R IC | 0.036 | 0.041 | 0.807 |
| **L LOC** | L PAC | 0.035 | 0.042 | 0.806 |
| **L IC** | R lingual | 0.035 | 0.041 | 0.806 |
| **L PAC** | L precentral | 0.035 | 0.041 | 0.805 |
| **R Hippo CA4** | R IC | 0.035 | 0.041 | 0.805 |
| **R Hippo CA1** | R MGN | 0.035 | 0.042 | 0.803 |
| **R lingual** | R PAC | 0.034 | 0.041 | 0.798 |
| **L PAC** | R SPC | 0.033 | 0.041 | 0.797 |
| **L lingual** | L MGN | 0.034 | 0.042 | 0.793 |
| **R IC** | R CN/SOC | 0.033 | 0.041 | 0.793 |
| **R precuneus** | R CN/SOC | 0.033 | 0.042 | 0.793 |
| **L IC** | R Hippo CA1 | 0.034 | 0.042 | 0.792 |
| **R MTC** | R CN/SOC | 0.033 | 0.041 | 0.789 |
| **L cMFC** | L PAC | 0.031 | 0.041 | 0.787 |
| **R LOC** | R CN/SOC | 0.032 | 0.041 | 0.786 |
| **L CN/SOC** | R Hippo GCDG | 0.033 | 0.041 | 0.783 |

| | | | | |
|---|---|---|---|---|
| **R Hippo GCDG** | R IC | 0.031 | 0.041 | 0.781 |
| **L MGN** | R Hippo | 0.032 | 0.042 | 0.778 |
| **L CLLPM Pulvinar** | L PAC | 0.031 | 0.042 | 0.777 |
| **L IC** | R Hippo | 0.031 | 0.042 | 0.774 |
| **L IC** | L PAC | 0.028 | 0.041 | 0.766 |
| **L CN/SOC** | R pericalc | 0.029 | 0.041 | 0.766 |
| **L PCC** | R CN/SOC | 0.028 | 0.042 | 0.759 |
| **R Hippo Tail** | R CN/SOC | 0.029 | 0.042 | 0.757 |
| **L PAC** | L Pallidum | 0.027 | 0.041 | 0.756 |
| **R Isth** | R CN/SOC | 0.028 | 0.041 | 0.754 |
| **R IPC** | R CN/SOC | 0.027 | 0.042 | 0.752 |
| **L Hippo Pres** | R PAC | 0.026 | 0.042 | 0.748 |
| **L PAC** | R parstr | 0.026 | 0.041 | 0.747 |
| **L Hippo CA3** | R PAC | 0.027 | 0.042 | 0.746 |
| **L Hippo** | R PAC | 0.026 | 0.041 | 0.744 |
| **L PAC** | R MGN | 0.026 | 0.041 | 0.743 |
| **L Hippo Tail** | L PAC | 0.026 | 0.041 | 0.742 |
| **L PAC** | L Pulvinar | 0.026 | 0.042 | 0.733 |
| **L IC** | R CN/SOC | 0.026 | 0.042 | 0.733 |
| **L paracentral** | R CN/SOC | 0.025 | 0.042 | 0.733 |
| **L IC** | L lingual | 0.025 | 0.042 | 0.732 |
| **L PAC** | L SFC | 0.024 | 0.041 | 0.73 |
| **R Hippo GCDG** | R MGN | 0.024 | 0.042 | 0.73 |
| **L PAC** | R cACC | 0.024 | 0.041 | 0.727 |
| **L PAC** | R Medio Dorsal | 0.023 | 0.041 | 0.727 |
| **L lingual** | R PAC | 0.024 | 0.042 | 0.725 |
| **L pericalc** | L CN/SOC | 0.024 | 0.042 | 0.724 |
| **R Hippo CA4** | R MGN | 0.024 | 0.041 | 0.724 |
| **L PAC** | L SPC | 0.023 | 0.042 | 0.722 |
| **L LOC** | R CN/SOC | 0.024 | 0.041 | 0.722 |
| **L Hippo CA4** | R PAC | 0.024 | 0.041 | 0.722 |
| **L MGN** | L PAC | 0.023 | 0.041 | 0.715 |
| **R IC** | R pericalc | 0.023 | 0.042 | 0.715 |
| **R pars** | R CN/SOC | 0.023 | 0.042 | 0.715 |
| **L MGN** | R Hippo CA4 | 0.022 | 0.041 | 0.714 |
| **L PAC** | L rMFC | 0.022 | 0.041 | 0.711 |
| **L IC** | R Hippo CA4 | 0.022 | 0.042 | 0.711 |
| **R Hippo CA3** | R PAC | 0.022 | 0.041 | 0.711 |
| **L Pulvinar** | R CN/SOC | 0.022 | 0.041 | 0.708 |
| **R cMFC** | R CN/SOC | 0.021 | 0.042 | 0.707 |
| **R Hippo CA1** | R PAC | 0.022 | 0.041 | 0.706 |
| **L PAC** | L parso | 0.022 | 0.042 | 0.704 |
| **L Hippo Mol layer HP** | R PAC | 0.021 | 0.042 | 0.698 |
| **L CLLPM Pulvinar** | R CN/SOC | 0.02 | 0.041 | 0.694 |

| | | | | |
|---|---|---|---|---|
| **R CN/SOC** | R VLD | 0.019 | 0.042 | 0.694 |
| **L MGN** | R Hippo GCDG | 0.02 | 0.042 | 0.693 |
| **R Hippo** | R PAC | 0.02 | 0.042 | 0.688 |
| **L cACC** | L PAC | 0.019 | 0.042 | 0.687 |
| **L cMFC** | R CN/SOC | 0.02 | 0.042 | 0.687 |
| **L CN/SOC** | R cuneus | 0.019 | 0.041 | 0.684 |
| **R paracentral** | R CN/SOC | 0.02 | 0.042 | 0.684 |
| **L pericalc** | R IC | 0.019 | 0.041 | 0.682 |
| **R MGN** | R CN/SOC | 0.018 | 0.041 | 0.678 |
| **L PAC** | L V Anterior | 0.018 | 0.041 | 0.673 |
| **L IC** | R Hippo GCDG | 0.017 | 0.042 | 0.669 |
| **L precuneus** | L CN/SOC | 0.017 | 0.041 | 0.666 |
| **L PAC** | R SFC | 0.017 | 0.042 | 0.665 |
| **R IC** | R MTC | 0.017 | 0.041 | 0.663 |
| **L VLD** | R CN/SOC | 0.017 | 0.042 | 0.663 |
| **L cuneus** | L CN/SOC | 0.016 | 0.041 | 0.66 |
| **R CN/SOC** | R SPC | 0.016 | 0.041 | 0.656 |
| **L precuneus** | R IC | 0.015 | 0.041 | 0.654 |
| **L MGN** | R CN/SOC | 0.017 | 0.042 | 0.654 |
| **L Isth** | L PAC | 0.015 | 0.041 | 0.652 |
| **L PAC** | R CLLPM Pulvinar | 0.015 | 0.041 | 0.65 |
| **R cuneus** | R IC | 0.015 | 0.041 | 0.65 |
| **L bankssts** | R CN/SOC | 0.016 | 0.042 | 0.649 |
| **L CN/SOC** | R MTC | 0.014 | 0.041 | 0.648 |
| **R PCC** | R CN/SOC | 0.014 | 0.041 | 0.646 |
| **L Medio Dorsal** | L PAC | 0.014 | 0.042 | 0.638 |
| **L Hippo Tail** | R CN/SOC | 0.013 | 0.042 | 0.632 |
| **L IC** | R pericalc | 0.013 | 0.041 | 0.631 |
| **L MGN** | L pericalc | 0.013 | 0.042 | 0.629 |
| **L parstr** | R CN/SOC | 0.013 | 0.041 | 0.627 |
| **L PAC** | R V Anterior | 0.013 | 0.041 | 0.624 |
| **L PAC** | L postcentral | 0.012 | 0.042 | 0.618 |
| **L cuneus** | R IC | 0.012 | 0.041 | 0.618 |
| **L MGN** | R pericalc | 0.012 | 0.041 | 0.615 |
| **L PAC** | R rACC | 0.012 | 0.042 | 0.615 |
| **L CN/SOC** | R LOC | 0.011 | 0.041 | 0.61 |
| **R MGN** | R pericalc | 0.011 | 0.042 | 0.61 |
| **L PAC** | R precentral | 0.011 | 0.041 | 0.609 |
| **L CN/SOC** | R precuneus | 0.01 | 0.042 | 0.605 |
| **L CN/SOC** | R IC | 0.011 | 0.042 | 0.604 |
| **L pericalc** | R MGN | 0.011 | 0.041 | 0.602 |
| **R IC** | R precuneus | 0.009 | 0.042 | 0.6 |
| **L IPC** | L CN/SOC | 0.01 | 0.041 | 0.598 |
| **R bankssts** | R CN/SOC | 0.01 | 0.042 | 0.598 |

| | | | | |
|---|---|---|---|---|
| **L PAC** | R Caudate | 0.01 | 0.041 | 0.597 |
| **L CN/SOC** | R Hippo Tail | 0.009 | 0.042 | 0.597 |
| **L IPC** | R IC | 0.009 | 0.042 | 0.597 |
| **R Hippo Tail** | R IC | 0.01 | 0.041 | 0.595 |
| **L Isth** | R CN/SOC | 0.009 | 0.041 | 0.592 |
| **R postcentral** | R CN/SOC | 0.007 | 0.041 | 0.58 |
| **R CN/SOC** | R PAC | 0.008 | 0.042 | 0.579 |
| **L IC** | L pericalc | 0.007 | 0.041 | 0.574 |
| **L SPC** | R CN/SOC | 0.007 | 0.042 | 0.574 |
| **L PAC** | L Putamen | 0.006 | 0.042 | 0.566 |
| **L PCC** | R IC | 0.006 | 0.042 | 0.566 |
| **L SFC** | R CN/SOC | 0.006 | 0.041 | 0.564 |
| **L PAC** | L rACC | 0.006 | 0.042 | 0.562 |
| **L Pallidum** | R CN/SOC | 0.006 | 0.041 | 0.562 |
| **L PCC** | L CN/SOC | 0.005 | 0.041 | 0.561 |
| **L CN/SOC** | R Isth | 0.005 | 0.041 | 0.56 |
| **L paracentral** | L CN/SOC | 0.005 | 0.041 | 0.559 |
| **L Anterior** | L PAC | 0.006 | 0.042 | 0.558 |
| **R IC** | R IPC | 0.005 | 0.042 | 0.557 |
| **L pericalc** | R PAC | 0.005 | 0.042 | 0.557 |
| **R pericalc** | R PAC | 0.005 | 0.041 | 0.554 |
| **L LOC** | L CN/SOC | 0.004 | 0.042 | 0.552 |
| **R MGN** | R MTC | 0.005 | 0.041 | 0.552 |
| **R IC** | R Isth | 0.004 | 0.042 | 0.548 |
| **R cuneus** | R MGN | 0.004 | 0.041 | 0.546 |
| **L CN/SOC** | R IPC | 0.004 | 0.042 | 0.544 |
| **L Caudate** | L PAC | 0.003 | 0.041 | 0.543 |
| **L paracentral** | R IC | 0.004 | 0.041 | 0.543 |
| **L precuneus** | R MGN | 0.004 | 0.042 | 0.543 |
| **R Hippo CA4** | R PAC | 0.003 | 0.041 | 0.536 |
| **L IC** | L CN/SOC | 0.003 | 0.041 | 0.535 |
| **L IC** | R cuneus | 0.003 | 0.042 | 0.534 |
| **L CN/SOC** | R pars | 0.003 | 0.042 | 0.534 |
| **L rMFC** | R CN/SOC | 0.003 | 0.041 | 0.532 |
| **L CN/SOC** | R cMFC | 0.002 | 0.041 | 0.529 |
| **R IC** | R LOC | 0.003 | 0.041 | 0.529 |
| **L CN/SOC** | R paracentral | 0.002 | 0.041 | 0.529 |
| **L MGN** | R cuneus | 0.002 | 0.041 | 0.528 |
| **L PAC** | R parso | 0.002 | 0.042 | 0.527 |
| **L MGN** | L precuneus | 0.002 | 0.041 | 0.526 |
| **L Pulvinar** | L CN/SOC | 0.002 | 0.041 | 0.523 |
| **R Hippo Tail** | R MGN | 0.002 | 0.042 | 0.521 |
| **R Hippo GCDG** | R PAC | 0.002 | 0.041 | 0.52 |
| **L cuneus** | R MGN | 0.002 | 0.042 | 0.518 |

| | | | | |
|---|---|---|---|---|
| **L cuneus** | L MGN | 0.001 | 0.041 | 0.515 |
| **L precentral** | R CN/SOC | 0.001 | 0.042 | 0.512 |
| **R MTC** | R PAC | 0 | 0.041 | 0.511 |
| **L PAC** | R rMFC | 0.001 | 0.041 | 0.51 |
| **L PAC** | R Anterior | 0.001 | 0.042 | 0.509 |
| **L IC** | R MTC | 0.001 | 0.042 | 0.509 |
| **R IC** | R pars | 0 | 0.041 | 0.507 |
| **R IC** | R paracentral | 0 | 0.041 | 0.504 |
| **L MGN** | R MTC | 0 | 0.041 | 0.503 |
| **L cuneus** | L IC | -0.001 | 0.041 | 0.502 |
| **R LOC** | R MGN | -0.001 | 0.041 | 0.501 |
| **L CLLPM Pulvinar** | L CN/SOC | 0 | 0.042 | 0.5 |
| **R IC** | R MGN | -0.001 | 0.042 | 0.497 |
| **L CN/SOC** | R SPC | -0.002 | 0.041 | 0.494 |
| **L IC** | L precuneus | -0.002 | 0.041 | 0.491 |
| **L CN/SOC** | R MGN | -0.002 | 0.042 | 0.491 |
| **L CN/SOC** | R VLD | -0.002 | 0.042 | 0.489 |
| **R cACC** | R CN/SOC | -0.002 | 0.042 | 0.486 |
| **L IPC** | R MGN | -0.003 | 0.042 | 0.484 |
| **L Anterior** | R CN/SOC | -0.002 | 0.041 | 0.483 |
| **L LOC** | R IC | -0.003 | 0.041 | 0.481 |
| **R Medio Dorsal** | R CN/SOC | -0.003 | 0.042 | 0.481 |
| **L cMFC** | L CN/SOC | -0.003 | 0.041 | 0.48 |
| **L MGN** | L CN/SOC | -0.002 | 0.042 | 0.478 |
| **L CN/SOC** | L VLD | -0.002 | 0.041 | 0.478 |
| **L cuneus** | R PAC | -0.003 | 0.041 | 0.477 |
| **R Isth** | R MGN | -0.003 | 0.041 | 0.476 |
| **L MGN** | R precuneus | -0.003 | 0.041 | 0.475 |
| **R cMFC** | R IC | -0.003 | 0.041 | 0.474 |
| **L VLD** | R IC | -0.004 | 0.042 | 0.471 |
| **L bankssts** | L CN/SOC | -0.004 | 0.041 | 0.467 |
| **L Pulvinar** | R IC | -0.004 | 0.042 | 0.464 |
| **L bankssts** | R IC | -0.005 | 0.041 | 0.46 |
| **R MGN** | R precuneus | -0.004 | 0.042 | 0.46 |
| **R cuneus** | R PAC | -0.006 | 0.042 | 0.46 |
| **R IC** | R SPC | -0.005 | 0.042 | 0.459 |
| **L IC** | R precuneus | -0.005 | 0.042 | 0.457 |
| **L MGN** | R Hippo Tail | -0.005 | 0.041 | 0.456 |
| **L rACC** | R CN/SOC | -0.006 | 0.041 | 0.454 |
| **L IPC** | L MGN | -0.005 | 0.042 | 0.453 |
| **L PCC** | R MGN | -0.005 | 0.042 | 0.453 |
| **R IC** | R PCC | -0.006 | 0.041 | 0.453 |
| **L CLLPM Pulvinar** | R IC | -0.006 | 0.041 | 0.45 |
| **L V Anterior** | R CN/SOC | -0.005 | 0.041 | 0.45 |

| | | | | |
|---|---|---|---|---|
| **R rACC** | R CN/SOC | -0.006 | 0.041 | 0.449 |
| **L precuneus** | R PAC | -0.006 | 0.041 | 0.449 |
| **R IC** | R VLD | -0.006 | 0.041 | 0.448 |
| **R CLLPM Pulvinar** | R CN/SOC | -0.006 | 0.042 | 0.446 |
| **R parstr** | R CN/SOC | -0.006 | 0.041 | 0.446 |
| **R SFC** | R CN/SOC | -0.007 | 0.042 | 0.446 |
| **L CN/SOC** | R PCC | -0.006 | 0.041 | 0.445 |
| **L MGN** | R Isth | -0.006 | 0.041 | 0.444 |
| **L MGN** | R IC | -0.007 | 0.042 | 0.441 |
| **L paracentral** | R MGN | -0.006 | 0.042 | 0.441 |
| **L PAC** | R Pallidum | -0.007 | 0.041 | 0.441 |
| **L IC** | R LOC | -0.007 | 0.041 | 0.438 |
| **L MGN** | R MGN | -0.007 | 0.042 | 0.437 |
| **L IC** | L IPC | -0.008 | 0.042 | 0.436 |
| **L parstr** | R IC | -0.008 | 0.041 | 0.436 |
| **R CN/SOC** | R V Anterior | -0.007 | 0.041 | 0.436 |
| **L paracentral** | R PAC | -0.007 | 0.042 | 0.434 |
| **L LOC** | R MGN | -0.007 | 0.041 | 0.432 |
| **R Caudate** | R CN/SOC | -0.007 | 0.041 | 0.43 |
| **L cACC** | R CN/SOC | -0.008 | 0.041 | 0.426 |
| **L IC** | L paracentral | -0.008 | 0.042 | 0.425 |
| **L CLLPM Pulvinar** | R MGN | -0.008 | 0.042 | 0.424 |
| **L PCC** | R PAC | -0.008 | 0.042 | 0.424 |
| **R bankssts** | R IC | -0.009 | 0.041 | 0.423 |
| **R IPC** | R MGN | -0.008 | 0.042 | 0.423 |
| **L Hippo Tail** | R IC | -0.009 | 0.041 | 0.42 |
| **L CLLPM Pulvinar** | L MGN | -0.009 | 0.042 | 0.418 |
| **L IC** | L PCC | -0.009 | 0.041 | 0.418 |
| **L cMFC** | R IC | -0.009 | 0.042 | 0.417 |
| **L Hippo Tail** | L CN/SOC | -0.01 | 0.041 | 0.415 |
| **L Caudate** | R CN/SOC | -0.01 | 0.042 | 0.412 |
| **R Hippo Tail** | R PAC | -0.01 | 0.041 | 0.41 |
| **L IC** | R Hippo Tail | -0.01 | 0.042 | 0.409 |
| **L IC** | R IPC | -0.01 | 0.041 | 0.409 |
| **L parso** | R CN/SOC | -0.01 | 0.042 | 0.409 |
| **L MGN** | R IPC | -0.01 | 0.041 | 0.408 |
| **L IC** | R Isth | -0.01 | 0.042 | 0.404 |
| **L IC** | R paracentral | -0.01 | 0.041 | 0.404 |
| **L IC** | R pars | -0.011 | 0.042 | 0.404 |
| **L MGN** | R LOC | -0.011 | 0.041 | 0.403 |
| **L MGN** | L paracentral | -0.011 | 0.042 | 0.402 |
| **L MGN** | L PCC | -0.011 | 0.041 | 0.402 |
| **R MGN** | R paracentral | -0.011 | 0.042 | 0.401 |
| **R Anterior** | R CN/SOC | -0.011 | 0.041 | 0.401 |

| | | | | |
|---|---|---|---|---|
| **L parstr** | L CN/SOC | -0.011 | 0.041 | 0.4 |
| **L IPC** | R PAC | -0.01 | 0.042 | 0.399 |
| **L VLD** | R MGN | -0.011 | 0.042 | 0.398 |
| **L bankssts** | R PAC | -0.011 | 0.042 | 0.398 |
| **R paracentral** | R PAC | -0.011 | 0.042 | 0.398 |
| **R LOC** | R PAC | -0.012 | 0.041 | 0.392 |
| **L CN/SOC** | R bankssts | -0.012 | 0.041 | 0.39 |
| **L CN/SOC** | R postcentral | -0.012 | 0.041 | 0.39 |
| **R IC** | R PAC | -0.012 | 0.042 | 0.389 |
| **L Medio Dorsal** | R CN/SOC | -0.012 | 0.042 | 0.384 |
| **R precuneus** | R PAC | -0.013 | 0.041 | 0.377 |
| **R IPC** | R PAC | -0.013 | 0.041 | 0.376 |
| **L Hippo Tail** | R MGN | -0.014 | 0.041 | 0.375 |
| **R MGN** | R pars | -0.013 | 0.042 | 0.374 |
| **L MGN** | L Pulvinar | -0.014 | 0.042 | 0.373 |
| **L CN/SOC** | L SPC | -0.014 | 0.042 | 0.372 |
| **L Pulvinar** | R MGN | -0.014 | 0.041 | 0.372 |
| **L Pallidum** | L CN/SOC | -0.014 | 0.042 | 0.371 |
| **L Pallidum** | R IC | -0.014 | 0.041 | 0.371 |
| **L VLD** | R PAC | -0.015 | 0.042 | 0.366 |
| **L Isth** | L CN/SOC | -0.014 | 0.041 | 0.364 |
| **R MGN** | R VLD | -0.015 | 0.042 | 0.362 |
| **R bankssts** | R PAC | -0.015 | 0.042 | 0.362 |
| **L MGN** | R paracentral | -0.016 | 0.042 | 0.359 |
| **L IC** | L Pulvinar | -0.016 | 0.041 | 0.358 |
| **L CN/SOC** | R PAC | -0.015 | 0.041 | 0.358 |
| **L IC** | L LOC | -0.015 | 0.041 | 0.353 |
| **L postcentral** | R CN/SOC | -0.016 | 0.041 | 0.352 |
| **L IC** | R VLD | -0.016 | 0.041 | 0.352 |
| **L IC** | L VLD | -0.016 | 0.041 | 0.351 |
| **R cMFC** | R PAC | -0.016 | 0.042 | 0.351 |
| **R VLD** | R PAC | -0.016 | 0.041 | 0.351 |
| **L LOC** | L MGN | -0.017 | 0.042 | 0.35 |
| **L IC** | R cMFC | -0.016 | 0.042 | 0.35 |
| **R pars** | R PAC | -0.017 | 0.041 | 0.35 |
| **R MGN** | R SPC | -0.016 | 0.042 | 0.349 |
| **L MGN** | L VLD | -0.016 | 0.041 | 0.347 |
| **L IC** | R SPC | -0.016 | 0.041 | 0.347 |
| **L MGN** | R pars | -0.017 | 0.042 | 0.346 |
| **R IC** | R postcentral | -0.017 | 0.041 | 0.344 |
| **L parstr** | R PAC | -0.017 | 0.042 | 0.343 |
| **R postcentral** | R PAC | -0.017 | 0.041 | 0.342 |
| **L SFC** | L CN/SOC | -0.018 | 0.041 | 0.337 |
| **L precentral** | L CN/SOC | -0.017 | 0.041 | 0.336 |

| | | | | |
|---|---|---|---|---|
| **L LOC** | R PAC | -0.018 | 0.041 | 0.335 |
| **R PCC** | R PAC | -0.019 | 0.042 | 0.334 |
| **L SPC** | R IC | -0.019 | 0.041 | 0.333 |
| **L IC** | R MGN | -0.018 | 0.042 | 0.333 |
| **R cMFC** | R MGN | -0.019 | 0.042 | 0.333 |
| **L IC** | L MGN | -0.019 | 0.042 | 0.332 |
| **L Isth** | R IC | -0.018 | 0.042 | 0.332 |
| **R rMFC** | R CN/SOC | -0.019 | 0.041 | 0.332 |
| **L CN/SOC** | R cACC | -0.019 | 0.041 | 0.328 |
| **L rMFC** | L CN/SOC | -0.018 | 0.041 | 0.327 |
| **L bankssts** | L IC | -0.018 | 0.041 | 0.326 |
| **L CLLPM Pulvinar** | L IC | -0.02 | 0.042 | 0.326 |
| **L MGN** | R cMFC | -0.019 | 0.041 | 0.326 |
| **L IC** | R PCC | -0.019 | 0.042 | 0.325 |
| **R MGN** | R PCC | -0.019 | 0.042 | 0.325 |
| **R precentral** | R CN/SOC | -0.019 | 0.042 | 0.325 |
| **L Hippo Tail** | L MGN | -0.02 | 0.042 | 0.324 |
| **L Anterior** | R IC | -0.019 | 0.041 | 0.324 |
| **L rMFC** | R IC | -0.019 | 0.041 | 0.319 |
| **R Isth** | R PAC | -0.019 | 0.042 | 0.319 |
| **L IC** | L parstr | -0.02 | 0.041 | 0.313 |
| **L SFC** | R IC | -0.02 | 0.041 | 0.313 |
| **L PAC** | R Putamen | -0.021 | 0.041 | 0.313 |
| **L bankssts** | L MGN | -0.022 | 0.042 | 0.31 |
| **L cMFC** | R PAC | -0.022 | 0.041 | 0.31 |
| **R SPC** | R PAC | -0.02 | 0.041 | 0.308 |
| **L precentral** | R IC | -0.021 | 0.041 | 0.307 |
| **L cMFC** | L IC | -0.022 | 0.041 | 0.301 |
| **L bankssts** | R MGN | -0.022 | 0.042 | 0.3 |
| **L parstr** | R MGN | -0.022 | 0.042 | 0.3 |
| **L MGN** | L parstr | -0.022 | 0.041 | 0.299 |
| **R IC** | R Medio Dorsal | -0.022 | 0.042 | 0.299 |
| **R MGN** | R postcentral | -0.022 | 0.041 | 0.298 |
| **L Isth** | R MGN | -0.022 | 0.041 | 0.297 |
| **L MGN** | R SPC | -0.022 | 0.041 | 0.297 |
| **L IC** | R bankssts | -0.023 | 0.041 | 0.296 |
| **L cMFC** | R MGN | -0.022 | 0.042 | 0.296 |
| **R bankssts** | R MGN | -0.023 | 0.041 | 0.292 |
| **L Anterior** | L CN/SOC | -0.023 | 0.042 | 0.291 |
| **R IC** | R parstr | -0.023 | 0.041 | 0.289 |
| **L CN/SOC** | R CLLPM Pulvinar | -0.023 | 0.041 | 0.286 |
| **L V Anterior** | R IC | -0.024 | 0.042 | 0.286 |
| **L MGN** | R VLD | -0.024 | 0.042 | 0.286 |
| **L Hippo Tail** | L IC | -0.024 | 0.041 | 0.284 |

| | | | | |
|---|---|---|---|---|
| **L MGN** | R PCC | -0.024 | 0.041 | 0.281 |
| **L CLLPM Pulvinar** | R PAC | -0.024 | 0.042 | 0.281 |
| **L MGN** | R bankssts | -0.024 | 0.042 | 0.28 |
| **L CN/SOC** | R parstr | -0.025 | 0.041 | 0.279 |
| **L cMFC** | L MGN | -0.024 | 0.041 | 0.276 |
| **R cACC** | R IC | -0.024 | 0.041 | 0.276 |
| **L CN/SOC** | R Medio Dorsal | -0.025 | 0.041 | 0.276 |
| **L IC** | L Pallidum | -0.026 | 0.041 | 0.273 |
| **L Putamen** | R CN/SOC | -0.025 | 0.041 | 0.272 |
| **L CN/SOC** | R rACC | -0.025 | 0.041 | 0.27 |
| **L Isth** | L MGN | -0.026 | 0.041 | 0.266 |
| **R CLLPM Pulvinar** | R IC | -0.025 | 0.041 | 0.266 |
| **L precentral** | R PAC | -0.026 | 0.041 | 0.263 |
| **L CN/SOC** | R SFC | -0.026 | 0.041 | 0.261 |
| **L rACC** | L CN/SOC | -0.027 | 0.042 | 0.26 |
| **L cACC** | L CN/SOC | -0.027 | 0.041 | 0.258 |
| **L IC** | R postcentral | -0.027 | 0.041 | 0.257 |
| **R Pallidum** | R CN/SOC | -0.027 | 0.042 | 0.257 |
| **L SFC** | R PAC | -0.028 | 0.041 | 0.254 |
| **L MGN** | R postcentral | -0.028 | 0.041 | 0.252 |
| **L IC** | R PAC | -0.028 | 0.042 | 0.25 |
| **L Pulvinar** | R PAC | -0.028 | 0.041 | 0.25 |
| **R Anterior** | R IC | -0.028 | 0.042 | 0.249 |
| **L CN/SOC** | R Caudate | -0.029 | 0.042 | 0.248 |
| **R IC** | R rACC | -0.028 | 0.041 | 0.248 |
| **R parso** | R CN/SOC | -0.028 | 0.041 | 0.245 |
| **L rACC** | R IC | -0.029 | 0.041 | 0.24 |
| **L SPC** | R MGN | -0.03 | 0.042 | 0.239 |
| **L Pallidum** | R PAC | -0.029 | 0.041 | 0.239 |
| **R IC** | R SFC | -0.029 | 0.042 | 0.238 |
| **R CLLPM Pulvinar** | R MGN | -0.029 | 0.042 | 0.237 |
| **L CN/SOC** | L V Anterior | -0.029 | 0.041 | 0.236 |
| **L SFC** | R MGN | -0.03 | 0.041 | 0.236 |
| **L precentral** | R MGN | -0.03 | 0.041 | 0.235 |
| **R IC** | R V Anterior | -0.03 | 0.041 | 0.235 |
| **L cACC** | R IC | -0.03 | 0.042 | 0.234 |
| **R MGN** | R PAC | -0.03 | 0.041 | 0.233 |
| **L SPC** | R PAC | -0.031 | 0.042 | 0.232 |
| **L IC** | L SFC | -0.031 | 0.042 | 0.231 |
| **L CN/SOC** | R Anterior | -0.03 | 0.041 | 0.23 |
| **L MGN** | R PAC | -0.031 | 0.042 | 0.23 |
| **L parso** | R PAC | -0.032 | 0.042 | 0.228 |
| **L rMFC** | R PAC | -0.031 | 0.041 | 0.228 |
| **R Caudate** | R IC | -0.031 | 0.041 | 0.225 |

| | | | | |
|---|---|---|---|---|
| **L parso** | R IC | -0.03 | 0.041 | 0.224 |
| **L Caudate** | R IC | -0.032 | 0.041 | 0.221 |
| **L IC** | L Isth | -0.032 | 0.041 | 0.22 |
| **L CN/SOC** | R V Anterior | -0.032 | 0.041 | 0.22 |
| **L Hippo Tail** | R PAC | -0.033 | 0.042 | 0.219 |
| **L Medio Dorsal** | L CN/SOC | -0.033 | 0.041 | 0.218 |
| **L IC** | L SPC | -0.032 | 0.041 | 0.218 |
| **R parstr** | R PAC | -0.032 | 0.041 | 0.218 |
| **L IC** | L precentral | -0.033 | 0.042 | 0.217 |
| **L Medio Dorsal** | R IC | -0.033 | 0.042 | 0.216 |
| **L IC** | R Medio Dorsal | -0.033 | 0.042 | 0.216 |
| **L IC** | L rMFC | -0.033 | 0.042 | 0.214 |
| **R cACC** | R PAC | -0.033 | 0.041 | 0.214 |
| **L Caudate** | L CN/SOC | -0.033 | 0.042 | 0.212 |
| **L MGN** | L SFC | -0.034 | 0.042 | 0.209 |
| **L MGN** | L SPC | -0.034 | 0.042 | 0.206 |
| **L parso** | L CN/SOC | -0.034 | 0.042 | 0.205 |
| **L MGN** | L Pallidum | -0.035 | 0.041 | 0.204 |
| **L postcentral** | L CN/SOC | -0.034 | 0.041 | 0.199 |
| **L IC** | R CLLPM Pulvinar | -0.036 | 0.042 | 0.195 |
| **L Pallidum** | R MGN | -0.036 | 0.042 | 0.193 |
| **L IC** | R cACC | -0.036 | 0.041 | 0.191 |
| **L MGN** | R CLLPM Pulvinar | -0.037 | 0.042 | 0.19 |
| **L cACC** | R PAC | -0.036 | 0.042 | 0.19 |
| **L MGN** | L precentral | -0.036 | 0.041 | 0.188 |
| **L CN/SOC** | R precentral | -0.037 | 0.042 | 0.186 |
| **L IC** | R parstr | -0.037 | 0.042 | 0.184 |
| **R SFC** | R PAC | -0.038 | 0.042 | 0.178 |
| **L IC** | L V Anterior | -0.038 | 0.041 | 0.177 |
| **L postcentral** | R IC | -0.038 | 0.042 | 0.176 |
| **L rACC** | R MGN | -0.038 | 0.042 | 0.175 |
| **L CN/SOC** | R rMFC | -0.038 | 0.042 | 0.175 |
| **R Medio Dorsal** | R PAC | -0.038 | 0.041 | 0.175 |
| **L Anterior** | L IC | -0.038 | 0.041 | 0.174 |
| **L rMFC** | R MGN | -0.038 | 0.042 | 0.174 |
| **R MGN** | R rACC | -0.039 | 0.042 | 0.173 |
| **R MGN** | R parstr | -0.04 | 0.041 | 0.17 |
| **L IC** | R SFC | -0.04 | 0.042 | 0.17 |
| **L MGN** | L rACC | -0.04 | 0.042 | 0.168 |
| **R cACC** | R MGN | -0.039 | 0.041 | 0.167 |
| **L IC** | R rACC | -0.04 | 0.041 | 0.166 |
| **L MGN** | L rMFC | -0.041 | 0.042 | 0.165 |
| **L V Anterior** | R PAC | -0.04 | 0.042 | 0.165 |
| **R Medio Dorsal** | R MGN | -0.04 | 0.041 | 0.163 |

| | | | | |
|---|---|---|---|---|
| **R CLLPM Pulvinar** | R PAC | -0.04 | 0.041 | 0.163 |
| **L Isth** | R PAC | -0.04 | 0.041 | 0.162 |
| **L cACC** | L IC | -0.042 | 0.042 | 0.159 |
| **R IC** | R rMFC | -0.041 | 0.041 | 0.158 |
| **L IC** | L rACC | -0.041 | 0.041 | 0.157 |
| **L MGN** | R cACC | -0.042 | 0.041 | 0.154 |
| **L MGN** | R parstr | -0.042 | 0.041 | 0.154 |
| **R MGN** | R SFC | -0.042 | 0.042 | 0.153 |
| **L MGN** | R rACC | -0.042 | 0.042 | 0.152 |
| **R rACC** | R PAC | -0.042 | 0.042 | 0.151 |
| **R IC** | R precentral | -0.043 | 0.042 | 0.15 |
| **L IC** | R V Anterior | -0.042 | 0.041 | 0.15 |
| **L Caudate** | L IC | -0.043 | 0.041 | 0.148 |
| **L IC** | R Caudate | -0.042 | 0.042 | 0.148 |
| **L rACC** | R PAC | -0.043 | 0.042 | 0.148 |
| **L IC** | R Anterior | -0.044 | 0.041 | 0.143 |
| **L Putamen** | R IC | -0.044 | 0.041 | 0.14 |
| **L cACC** | R MGN | -0.045 | 0.042 | 0.139 |
| **L IC** | L parso | -0.044 | 0.042 | 0.138 |
| **L MGN** | R SFC | -0.045 | 0.042 | 0.135 |
| **R Caudate** | R PAC | -0.045 | 0.041 | 0.133 |
| **R precentral** | R PAC | -0.046 | 0.042 | 0.132 |
| **L V Anterior** | R MGN | -0.046 | 0.041 | 0.131 |
| **L MGN** | R Medio Dorsal | -0.047 | 0.041 | 0.128 |
| **L postcentral** | R MGN | -0.047 | 0.041 | 0.128 |
| **L Medio Dorsal** | R MGN | -0.047 | 0.041 | 0.127 |
| **L cACC** | L MGN | -0.048 | 0.042 | 0.124 |
| **L CN/SOC** | R parso | -0.048 | 0.042 | 0.124 |
| **R V Anterior** | R PAC | -0.047 | 0.041 | 0.124 |
| **L IC** | L Medio Dorsal | -0.046 | 0.041 | 0.123 |
| **L Anterior** | R MGN | -0.048 | 0.041 | 0.123 |
| **L parso** | R MGN | -0.048 | 0.042 | 0.122 |
| **L Medio Dorsal** | R PAC | -0.047 | 0.041 | 0.118 |
| **L postcentral** | R PAC | -0.048 | 0.042 | 0.118 |
| **L CN/SOC** | R Pallidum | -0.048 | 0.041 | 0.117 |
| **L Anterior** | R PAC | -0.05 | 0.042 | 0.116 |
| **L Putamen** | L CN/SOC | -0.049 | 0.041 | 0.115 |
| **R Putamen** | R CN/SOC | -0.05 | 0.042 | 0.115 |
| **R MGN** | R V Anterior | -0.05 | 0.041 | 0.114 |
| **L IC** | L postcentral | -0.05 | 0.042 | 0.112 |
| **L MGN** | L V Anterior | -0.049 | 0.042 | 0.112 |
| **R IC** | R parso | -0.05 | 0.042 | 0.112 |
| **L Caudate** | R PAC | -0.05 | 0.042 | 0.112 |
| **L MGN** | L postcentral | -0.051 | 0.042 | 0.109 |

| | | | | |
|---|---|---|---|---|
| **L Medio Dorsal** | L MGN | -0.05 | 0.041 | 0.108 |
| **L MGN** | L parso | -0.051 | 0.042 | 0.107 |
| **R parso** | R PAC | -0.051 | 0.042 | 0.107 |
| **L Anterior** | L MGN | -0.051 | 0.042 | 0.106 |
| **R Caudate** | R MGN | -0.052 | 0.042 | 0.105 |
| **R MGN** | R precentral | -0.051 | 0.042 | 0.104 |
| **R IC** | R Pallidum | -0.051 | 0.041 | 0.102 |
| **L MGN** | R V Anterior | -0.053 | 0.041 | 0.099 |
| **L Caudate** | R MGN | -0.053 | 0.042 | 0.097 |
| **L Putamen** | R PAC | -0.054 | 0.042 | 0.095 |
| **L IC** | R precentral | -0.053 | 0.041 | 0.093 |
| **L MGN** | R Caudate | -0.056 | 0.042 | 0.088 |
| **L IC** | R rMFC | -0.054 | 0.041 | 0.088 |
| **R rMFC** | R PAC | -0.056 | 0.041 | 0.086 |
| **R Anterior** | R PAC | -0.057 | 0.042 | 0.082 |
| **L MGN** | R precentral | -0.057 | 0.042 | 0.08 |
| **R Anterior** | R MGN | -0.057 | 0.041 | 0.078 |
| **L MGN** | R Anterior | -0.058 | 0.042 | 0.076 |
| **L Caudate** | L MGN | -0.058 | 0.042 | 0.075 |
| **L IC** | L Putamen | -0.059 | 0.041 | 0.072 |
| **R MGN** | R rMFC | -0.06 | 0.042 | 0.072 |
| **L IC** | R Pallidum | -0.061 | 0.042 | 0.07 |
| **L Putamen** | R MGN | -0.06 | 0.041 | 0.069 |
| **L IC** | R parso | -0.061 | 0.042 | 0.066 |
| **L MGN** | R rMFC | -0.062 | 0.042 | 0.062 |
| **R Pallidum** | R PAC | -0.062 | 0.042 | 0.061 |
| **L MGN** | L Putamen | -0.065 | 0.041 | 0.053 |
| **R MGN** | R parso | -0.067 | 0.041 | 0.05 |
| **R MGN** | R Pallidum | -0.068 | 0.042 | 0.047 |
| **L MGN** | R parso | -0.071 | 0.042 | 0.042 |
| **R IC** | R Putamen | -0.071 | 0.041 | 0.041 |
| **L CN/SOC** | R Putamen | -0.072 | 0.042 | 0.04 |
| **L MGN** | R Pallidum | -0.072 | 0.041 | 0.037 |
| **R Putamen** | R PAC | -0.076 | 0.041 | 0.029 |
| **L IC** | R Putamen | -0.082 | 0.042 | 0.023 |
| **R MGN** | R Putamen | -0.086 | 0.041 | 0.017 |
| **L MGN** | R Putamen | -0.093 | 0.041 | 0.011 |

#### Appendix D: Summary statistics of functional network nodal measures, CHUU-CPHIV.

*Table 4: Summary statistics for CHUU vs CPHIV for the nodal graph measures degree, strength, transitivity, nodal and local efficiency. Auditory ROI rows are marked in green.*

| | ***degree*** | | | | ***strength*** | | | | ***transitivity*** | | | | ***nodal efficiency*** | | | | ***local efficiency*** | | |
|---|---|---|---|---|---|---|---|---|---|---|---|---|---|---|---|---|---|---|---|
| ***ROI*** | ***mean*** | ***SD*** | ***P+*** | | ***mean*** | ***SD*** | ***P+*** | | ***mean*** | ***SD*** | ***P+*** | | ***mean*** | ***SD*** | ***P+*** | | ***mean*** | ***SD*** | ***P+*** |
| **L Anterior** | 1.320 | 3.252 | 0.666 | | 0.663 | 1.909 | 0.636 | | 0.018 | 0.041 | 0.671 | | 0.050 | 0.062 | 0.790 | | 0.070 | 0.062 | 0.872 |
| **L bankssts** | 1.471 | 3.011 | 0.687 | | 0.739 | 1.888 | 0.649 | | 0.019 | 0.041 | 0.675 | | 0.045 | 0.062 | 0.764 | | 0.053 | 0.060 | 0.815 |
| **L cACC** | 1.220 | 2.988 | 0.658 | | 0.653 | 1.886 | 0.632 | | 0.018 | 0.041 | 0.669 | | 0.047 | 0.062 | 0.775 | | 0.048 | 0.060 | 0.790 |
| **L Caudate** | 1.291 | 2.982 | 0.667 | | 0.656 | 1.887 | 0.634 | | 0.019 | 0.041 | 0.675 | | 0.049 | 0.062 | 0.787 | | 0.053 | 0.060 | 0.814 |
| **L CLLPM_Pulvinar** | 1.320 | 2.992 | 0.671 | | 0.642 | 1.904 | 0.630 | | 0.020 | 0.041 | 0.687 | | 0.048 | 0.062 | 0.780 | | 0.055 | 0.060 | 0.820 |
| **L cMFC** | 1.246 | 2.985 | 0.662 | | 0.622 | 1.889 | 0.627 | | 0.018 | 0.041 | 0.667 | | 0.051 | 0.063 | 0.791 | | 0.051 | 0.060 | 0.805 |
| **L cuneus** | 1.100 | 3.015 | 0.644 | | 0.720 | 1.884 | 0.646 | | 0.018 | 0.041 | 0.667 | | 0.045 | 0.062 | 0.765 | | 0.053 | 0.060 | 0.815 |
| **L Hippo** | 3.888 | 3.584 | 0.866 | | 0.857 | 1.900 | 0.671 | | 0.020 | 0.041 | 0.688 | | 0.047 | 0.062 | 0.779 | | 0.063 | 0.061 | 0.854 |
| **L Hippo_CA1** | 1.450 | 3.455 | 0.666 | | 0.841 | 1.896 | 0.669 | | 0.020 | 0.041 | 0.682 | | 0.047 | 0.062 | 0.779 | | 0.051 | 0.064 | 0.793 |
| **L Hippo_CA3** | 3.010 | 3.373 | 0.817 | | 0.831 | 1.892 | 0.666 | | 0.020 | 0.041 | 0.687 | | 0.046 | 0.062 | 0.770 | | 0.062 | 0.063 | 0.839 |
| **L Hippo_CA4** | 3.125 | 3.232 | 0.835 | | 0.798 | 1.886 | 0.662 | | 0.021 | 0.041 | 0.691 | | 0.047 | 0.062 | 0.776 | | 0.058 | 0.060 | 0.834 |
| **L Hippo_Mol_layer_HP** | 1.864 | 3.078 | 0.732 | | 0.746 | 1.888 | 0.650 | | 0.022 | 0.041 | 0.702 | | 0.050 | 0.062 | 0.789 | | 0.059 | 0.061 | 0.837 |
| **L Hippo_Pres** | 5.454 | 3.859 | 0.928 | | 1.016 | 1.917 | 0.699 | | 0.019 | 0.041 | 0.677 | | 0.047 | 0.062 | 0.777 | | 0.063 | 0.061 | 0.848 |
| **L Hippo_Tail** | 1.050 | 3.023 | 0.637 | | 0.645 | 1.896 | 0.632 | | 0.021 | 0.041 | 0.698 | | 0.045 | 0.062 | 0.765 | | 0.050 | 0.062 | 0.794 |
| **L IC** | 9.051 | 4.324 | 0.985 | | 0.903 | 1.924 | 0.678 | | 0.022 | 0.041 | 0.701 | | 0.053 | 0.067 | 0.786 | | 0.087 | 0.066 | 0.909 |
| **L IPC** | 1.446 | 3.054 | 0.682 | | 0.778 | 1.888 | 0.656 | | 0.018 | 0.041 | 0.664 | | 0.050 | 0.062 | 0.787 | | 0.054 | 0.060 | 0.818 |
| **L Isth** | 1.192 | 2.988 | 0.655 | | 0.706 | 1.887 | 0.642 | | 0.017 | 0.041 | 0.657 | | 0.045 | 0.062 | 0.767 | | 0.055 | 0.060 | 0.823 |
| **L lingual** | 1.083 | 3.014 | 0.639 | | 0.786 | 1.887 | 0.657 | | 0.017 | 0.041 | 0.660 | | 0.046 | 0.062 | 0.773 | | 0.056 | 0.061 | 0.826 |
| **L LOC** | 2.000 | 3.045 | 0.744 | | 0.858 | 1.889 | 0.673 | | 0.017 | 0.041 | 0.661 | | 0.048 | 0.062 | 0.782 | | 0.054 | 0.060 | 0.817 |
| **L Medio_Dorsal** | 1.308 | 2.990 | 0.669 | | 0.612 | 1.899 | 0.625 | | 0.019 | 0.041 | 0.677 | | 0.044 | 0.062 | 0.763 | | 0.053 | 0.060 | 0.814 |

| | | | | | | | | | | | | | | | | | | | |
|---|---|---|---|---|---|---|---|---|---|---|---|---|---|---|---|---|---|---|---|
| **L MGN** | 1.896 | 3.096 | 0.733 | | 0.881 | 1.903 | 0.675 | | 0.023 | 0.041 | 0.709 | | 0.047 | 0.062 | 0.774 | | 0.055 | 0.060 | 0.824 |
| **L PAC** | 1.416 | 3.040 | 0.680 | | 0.943 | 1.907 | 0.688 | | 0.025 | 0.041 | 0.727 | | 0.042 | 0.063 | 0.748 | | 0.060 | 0.061 | 0.842 |
| **L Pallidum** | 1.588 | 3.008 | 0.702 | | 0.742 | 1.888 | 0.650 | | 0.020 | 0.041 | 0.684 | | 0.037 | 0.065 | 0.721 | | 0.058 | 0.060 | 0.835 |
| **L paracentral** | 1.234 | 2.993 | 0.659 | | 0.735 | 1.884 | 0.648 | | 0.018 | 0.041 | 0.665 | | 0.045 | 0.062 | 0.766 | | 0.051 | 0.060 | 0.804 |
| **L parso** | 1.223 | 2.988 | 0.659 | | 0.635 | 1.888 | 0.627 | | 0.017 | 0.041 | 0.655 | | 0.048 | 0.062 | 0.779 | | 0.052 | 0.060 | 0.813 |
| **L parstr** | 1.315 | 2.987 | 0.669 | | 0.722 | 1.883 | 0.647 | | 0.018 | 0.041 | 0.666 | | 0.045 | 0.062 | 0.764 | | 0.052 | 0.060 | 0.811 |
| **L PCC** | 1.017 | 3.067 | 0.628 | | 0.702 | 1.887 | 0.641 | | 0.017 | 0.041 | 0.656 | | 0.049 | 0.062 | 0.785 | | 0.052 | 0.060 | 0.811 |
| **L pericalc** | 1.392 | 2.984 | 0.681 | | 0.749 | 1.886 | 0.652 | | 0.018 | 0.041 | 0.668 | | 0.047 | 0.062 | 0.775 | | 0.054 | 0.060 | 0.822 |
| **L postcentral** | 1.333 | 3.000 | 0.672 | | 0.589 | 1.898 | 0.617 | | 0.017 | 0.041 | 0.655 | | 0.048 | 0.062 | 0.776 | | 0.054 | 0.060 | 0.820 |
| **L precentral** | 1.180 | 2.993 | 0.654 | | 0.632 | 1.887 | 0.628 | | 0.018 | 0.041 | 0.670 | | 0.051 | 0.063 | 0.794 | | 0.054 | 0.060 | 0.816 |
| **L precuneus** | 1.162 | 2.986 | 0.651 | | 0.684 | 1.887 | 0.637 | | 0.017 | 0.041 | 0.659 | | 0.048 | 0.062 | 0.783 | | 0.053 | 0.060 | 0.814 |
| **L Pulvinar** | 6.935 | 4.371 | 0.939 | | 0.891 | 1.950 | 0.675 | | 0.022 | 0.041 | 0.699 | | 0.097 | 0.089 | 0.865 | | 0.127 | 0.073 | 0.960 |
| **L Putamen** | 0.130 | 3.083 | 0.517 | | 0.381 | 1.935 | 0.579 | | 0.019 | 0.041 | 0.674 | | 0.045 | 0.062 | 0.764 | | 0.054 | 0.060 | 0.818 |
| **L rACC** | 1.262 | 2.993 | 0.663 | | 0.663 | 1.889 | 0.635 | | 0.018 | 0.041 | 0.671 | | 0.044 | 0.062 | 0.763 | | 0.052 | 0.060 | 0.809 |
| **L rMFC** | 1.308 | 2.979 | 0.670 | | 0.639 | 1.889 | 0.629 | | 0.017 | 0.041 | 0.660 | | 0.050 | 0.062 | 0.788 | | 0.048 | 0.061 | 0.792 |
| **L SFC** | 0.985 | 3.019 | 0.626 | | 0.510 | 1.902 | 0.603 | | 0.017 | 0.041 | 0.657 | | 0.048 | 0.062 | 0.782 | | 0.046 | 0.060 | 0.781 |
| **L CN/SOC** | 0.193 | 4.011 | 0.521 | | 0.868 | 1.956 | 0.668 | | 0.023 | 0.041 | 0.710 | | 0.046 | 0.063 | 0.768 | | 0.080 | 0.068 | 0.883 |
| **L SPC** | 1.543 | 2.993 | 0.698 | | 0.701 | 1.885 | 0.641 | | 0.018 | 0.041 | 0.669 | | 0.049 | 0.062 | 0.785 | | 0.053 | 0.060 | 0.815 |
| **L V_Anterior** | 1.172 | 3.006 | 0.652 | | 0.587 | 1.894 | 0.618 | | 0.019 | 0.041 | 0.674 | | 0.053 | 0.064 | 0.795 | | 0.054 | 0.060 | 0.821 |
| **L VLD** | 1.231 | 2.999 | 0.659 | | 0.708 | 1.885 | 0.643 | | 0.018 | 0.041 | 0.664 | | 0.050 | 0.062 | 0.788 | | 0.056 | 0.060 | 0.827 |
| **R Anterior** | 1.660 | 2.997 | 0.712 | | 0.762 | 1.883 | 0.653 | | 0.017 | 0.041 | 0.658 | | 0.052 | 0.063 | 0.794 | | 0.052 | 0.060 | 0.809 |
| **R bankssts** | 1.509 | 2.992 | 0.694 | | 0.692 | 1.883 | 0.641 | | 0.016 | 0.041 | 0.645 | | 0.048 | 0.062 | 0.778 | | 0.054 | 0.060 | 0.819 |
| **R cACC** | 1.111 | 2.990 | 0.643 | | 0.637 | 1.886 | 0.630 | | 0.016 | 0.041 | 0.646 | | 0.046 | 0.062 | 0.769 | | 0.050 | 0.060 | 0.802 |
| **R Caudate** | 1.069 | 3.004 | 0.639 | | 0.637 | 1.889 | 0.630 | | 0.017 | 0.041 | 0.658 | | 0.047 | 0.062 | 0.773 | | 0.051 | 0.060 | 0.808 |
| **R CLLPM_Pulvinar** | 1.267 | 2.984 | 0.664 | | 0.657 | 1.888 | 0.633 | | 0.015 | 0.041 | 0.641 | | 0.049 | 0.062 | 0.780 | | 0.049 | 0.060 | 0.797 |
| **R cMFC** | 1.359 | 2.981 | 0.675 | | 0.704 | 1.885 | 0.642 | | 0.015 | 0.041 | 0.643 | | 0.048 | 0.062 | 0.781 | | 0.049 | 0.060 | 0.797 |
| **R cuneus** | 1.542 | 2.993 | 0.697 | | 0.706 | 1.885 | 0.642 | | 0.015 | 0.041 | 0.637 | | 0.048 | 0.062 | 0.780 | | 0.057 | 0.060 | 0.832 |

| | | | | | | | | | | | | | | | | | | | |
|---|---|---|---|---|---|---|---|---|---|---|---|---|---|---|---|---|---|---|---|
| **R Hippo** | 3.196 | 3.189 | 0.844 | | 0.760 | 1.893 | 0.654 | | 0.016 | 0.041 | 0.652 | | 0.049 | 0.062 | 0.781 | | 0.057 | 0.060 | 0.831 |
| **R Hippo_CA1** | 3.158 | 3.166 | 0.843 | | 0.761 | 1.892 | 0.654 | | 0.019 | 0.041 | 0.679 | | 0.049 | 0.062 | 0.785 | | 0.057 | 0.060 | 0.828 |
| **R Hippo_CA3** | 3.847 | 3.217 | 0.888 | | 0.931 | 1.896 | 0.685 | | 0.018 | 0.041 | 0.666 | | 0.052 | 0.063 | 0.795 | | 0.058 | 0.060 | 0.831 |
| **R Hippo_CA4** | 2.739 | 3.111 | 0.814 | | 0.780 | 1.886 | 0.658 | | 0.018 | 0.041 | 0.669 | | 0.044 | 0.062 | 0.763 | | 0.060 | 0.061 | 0.840 |
| **R Hippo_GCDG** | 2.634 | 3.088 | 0.806 | | 0.763 | 1.886 | 0.655 | | 0.017 | 0.041 | 0.657 | | 0.046 | 0.062 | 0.771 | | 0.059 | 0.060 | 0.836 |
| **R Hippo_Tail** | 1.560 | 2.997 | 0.700 | | 0.800 | 1.885 | 0.661 | | 0.017 | 0.041 | 0.662 | | 0.049 | 0.062 | 0.785 | | 0.054 | 0.060 | 0.818 |
| **R IC** | 6.936 | 4.634 | 0.944 | | 0.977 | 1.922 | 0.691 | | 0.026 | 0.041 | 0.735 | | 0.039 | 0.070 | 0.714 | | 0.125 | 0.071 | 0.962 |
| **R IPC** | 1.267 | 2.981 | 0.666 | | 0.663 | 1.885 | 0.634 | | 0.015 | 0.041 | 0.640 | | 0.052 | 0.062 | 0.798 | | 0.053 | 0.060 | 0.814 |
| **R Isth** | 1.458 | 2.985 | 0.688 | | 0.777 | 1.887 | 0.656 | | 0.014 | 0.041 | 0.632 | | 0.050 | 0.062 | 0.788 | | 0.050 | 0.060 | 0.803 |
| **R lingual** | 1.515 | 3.000 | 0.693 | | 0.762 | 1.884 | 0.654 | | 0.015 | 0.041 | 0.638 | | 0.045 | 0.062 | 0.768 | | 0.049 | 0.060 | 0.799 |
| **R LOC** | 1.765 | 3.007 | 0.722 | | 0.752 | 1.885 | 0.652 | | 0.015 | 0.041 | 0.641 | | 0.047 | 0.062 | 0.777 | | 0.050 | 0.060 | 0.802 |
| **R Medio_Dorsal** | 1.392 | 2.994 | 0.680 | | 0.628 | 1.894 | 0.628 | | 0.017 | 0.041 | 0.657 | | 0.048 | 0.062 | 0.782 | | 0.056 | 0.060 | 0.828 |
| **R MGN** | 1.998 | 3.169 | 0.740 | | 0.759 | 1.903 | 0.653 | | 0.022 | 0.041 | 0.707 | | 0.042 | 0.063 | 0.750 | | 0.073 | 0.062 | 0.880 |
| **R MTC** | 1.133 | 2.995 | 0.646 | | 0.777 | 1.884 | 0.656 | | 0.018 | 0.041 | 0.666 | | 0.042 | 0.063 | 0.749 | | 0.052 | 0.060 | 0.808 |
| **R PAC** | 1.362 | 2.983 | 0.677 | | 0.740 | 1.894 | 0.650 | | 0.023 | 0.041 | 0.713 | | 0.017 | 0.077 | 0.622 | | 0.058 | 0.061 | 0.830 |
| **R Pallidum** | 1.566 | 2.986 | 0.700 | | 0.714 | 1.885 | 0.643 | | 0.016 | 0.041 | 0.644 | | 0.047 | 0.063 | 0.774 | | 0.050 | 0.060 | 0.799 |
| **R paracentral** | 1.379 | 2.981 | 0.678 | | 0.699 | 1.883 | 0.640 | | 0.014 | 0.041 | 0.632 | | 0.050 | 0.062 | 0.787 | | 0.051 | 0.060 | 0.809 |
| **R pars** | 1.256 | 2.993 | 0.663 | | 0.722 | 1.886 | 0.647 | | 0.015 | 0.041 | 0.638 | | 0.049 | 0.062 | 0.783 | | 0.050 | 0.060 | 0.801 |
| **R parso** | 1.148 | 2.984 | 0.648 | | 0.629 | 1.890 | 0.628 | | 0.015 | 0.041 | 0.635 | | 0.048 | 0.062 | 0.783 | | 0.050 | 0.060 | 0.799 |
| **R parstr** | 1.239 | 2.985 | 0.661 | | 0.740 | 1.887 | 0.650 | | 0.016 | 0.041 | 0.651 | | 0.049 | 0.062 | 0.786 | | 0.049 | 0.060 | 0.795 |
| **R PCC** | 1.245 | 2.988 | 0.661 | | 0.703 | 1.884 | 0.642 | | 0.016 | 0.041 | 0.651 | | 0.047 | 0.062 | 0.777 | | 0.053 | 0.060 | 0.815 |
| **R pericalc** | 1.498 | 2.986 | 0.691 | | 0.736 | 1.883 | 0.650 | | 0.016 | 0.041 | 0.646 | | 0.047 | 0.062 | 0.776 | | 0.053 | 0.060 | 0.812 |
| **R postcentral** | 1.358 | 2.982 | 0.675 | | 0.695 | 1.883 | 0.640 | | 0.015 | 0.041 | 0.636 | | 0.047 | 0.062 | 0.776 | | 0.054 | 0.060 | 0.820 |
| **R precentral** | 1.202 | 2.979 | 0.656 | | 0.618 | 1.888 | 0.625 | | 0.016 | 0.041 | 0.646 | | 0.049 | 0.062 | 0.785 | | 0.053 | 0.060 | 0.814 |
| **R precuneus** | 1.433 | 2.984 | 0.687 | | 0.714 | 1.887 | 0.643 | | 0.015 | 0.041 | 0.639 | | 0.048 | 0.062 | 0.779 | | 0.055 | 0.060 | 0.824 |
| **R Putamen** | 1.197 | 2.991 | 0.656 | | 0.548 | 1.900 | 0.611 | | 0.014 | 0.041 | 0.630 | | 0.049 | 0.062 | 0.788 | | 0.051 | 0.060 | 0.806 |
| **R rACC** | 1.376 | 2.983 | 0.677 | | 0.697 | 1.883 | 0.641 | | 0.015 | 0.041 | 0.641 | | 0.049 | 0.062 | 0.786 | | 0.050 | 0.060 | 0.803 |

| | | | | | | | | | | | | | | | | | | | |
|---|---|---|---|---|---|---|---|---|---|---|---|---|---|---|---|---|---|---|---|
| **R rMFC** | 1.196 | 2.981 | 0.656 | | 0.668 | 1.888 | 0.635 | | 0.016 | 0.041 | 0.649 | | 0.049 | 0.062 | 0.783 | | 0.049 | 0.060 | 0.797 |
| **R SFC** | 1.118 | 3.008 | 0.645 | | 0.792 | 1.888 | 0.659 | | 0.014 | 0.041 | 0.627 | | 0.045 | 0.062 | 0.765 | | 0.050 | 0.060 | 0.801 |
| **R CN/SOC** | 3.897 | 4.110 | 0.829 | | 0.906 | 1.953 | 0.679 | | 0.021 | 0.041 | 0.694 | | 0.040 | 0.067 | 0.735 | | 0.127 | 0.073 | 0.963 |
| **R SPC** | 1.504 | 2.984 | 0.694 | | 0.709 | 1.882 | 0.644 | | 0.016 | 0.041 | 0.646 | | 0.049 | 0.062 | 0.782 | | 0.052 | 0.060 | 0.814 |
| **R V_Anterior** | 1.206 | 2.984 | 0.656 | | 0.686 | 1.886 | 0.640 | | 0.016 | 0.041 | 0.647 | | 0.046 | 0.062 | 0.768 | | 0.052 | 0.060 | 0.810 |
| **R VLD** | 1.451 | 2.981 | 0.687 | | 0.767 | 1.883 | 0.654 | | 0.016 | 0.041 | 0.648 | | 0.049 | 0.062 | 0.783 | | 0.053 | 0.060 | 0.814 |
